# Live-cell magnetic micromanipulation of recycling endosomes reveals their direct effect on actin-based protrusions to promote invasive migration

**DOI:** 10.1101/2024.11.11.622870

**Authors:** Jakub Gemperle, Domenik Liße, Marie Kappen, Emilie Secret, Mathieu Coppey, Martin Gregor, Christine Menager, Jacob Piehler, Patrick Caswell

## Abstract

Endocytic recycling pathways play key roles in the re-routing of cargoes through the cell to control a broad range of cellular processes, and many vesicle trafficking regulators are implicated in progression of disease such as cancer. The Rab11 family (Rab11a, Rab11b, and Rab25) control return of internalised cargoes to the plasma membrane, and Rab25 has been implicated in the aggressiveness of cancer by promoting invasive migration. However, whilst Rab25 vesicles distribute to the leading of edge of moving cells, how directly they contribute to cell protrusion is not clear. Here we adopt a magnetogenetic approach that allows direct manipulation of Rab25 positioning to show that localisation to the cell periphery drives the formation of F-actin protrusions. We demonstrate that endogenous Rab25 vesicles coordinate the positioning of key cargoes, including the actin regulator FMNL1 and integrin β1, with the activation of Rho GTPases at the plasma membrane to generate and maintain F-actin rich filopodial protrusions and promote cancer cell invasive migration in 3D matrix.

## Introduction

Endocytic trafficking controls how cells sense and respond to their environment (Jacquemet, Humphries, et al., 2013; Scita and Di Fiore, 2010) and accumulating evidence suggests a fundamental role in many physiological processes and pathological conditions (Golachowska et al., 2010; Jin et al., 2021; Kelly et al., 2012). Rab GTPases control specific steps within the intricate vesicular trafficking pathways, including how endosomes are positioned. It has been proposed that specific localisation of endosomes contributes to polarization, local growth and migration of cells, either through selective delivery of receptors, and/or through localized signalling (Eva et al., 2012; Golachowska et al., 2010; Higuchi et al., 2014; Sadowski *et al*., 2009; Vaidžiulyte *et al*., 2022). However, it is challenging to directly address the functional consequences of endosomal positioning, and it remains unclear whether Rab GTPases simply position vesicle and their receptor cargoes in a local area for utilisation at the membrane, or if they play an active role in determining the initiation of protrusions.

The Rab11 subfamily, comprised of Rab11a, Rab11b, and Rab25 (also known as Rab11c), are key regulators of endocytic recycling. They control the return of internalized membrane-associated cargos to the cell surface (Zerial and McBride, 2001) through recycling endosomes. Accumulating evidence suggests that polarized Rab11 trafficking contributes to aspects of tumorigenicity (Caswell *et al*., 2007; Cho and Lee, 2019; Gebhardt *et al*., 2005; Kelly *et al*., 2012; Mitra *et al*., 2012; Ray *et al*., 1997; Wang *et al*., 2017). Furthermore, Rab11 recycling endosomes have been demonstrated to nucleate F-actin from vesicles by recruiting actin nucleation factors to organise the actin network into tracks that allow for microtubule-independent vesicle movement (Pylypenko *et al*., 2016; Schuh, 2011). The Rab11 family has also been linked to cell motility. Rab11a/b has been demonstrated to facilitate cancer cell invasion by promoting the formation of filopodia-tipped protrusions (Gemperle *et al*., 2022; Jacquemet, Green, *et al*., 2013; Paul *et al*., 2015; Wilson *et al*., 2023), suggesting a link between the Rab11 family and F-actin nucleation that extends beyond formation of tracks for vesicles. Unlike the ubiquitous Rab11s (Rab11a and Rab11b), Rab25 has a limited expression profile in normal tissue (Goldenring *et al*., 1993) and has been shown to be upregulated in approximately half of ovarian and breast tumors, correlating with their aggressiveness both clinically and in mouse models (Cheng *et al*., 2004; Jeong *et al*., 2018). Rab25 directs the localization of integrin α5β1-recycling vesicles/endosomes to the leading edge of migrating cells to promote the formation of long actin-rich pseudopodia and enhance ability of tumor cells to invade the extracellular matrix (Caswell *et al*., 2007; Dozynkiewicz *et al*., 2012; Gemperle *et al*., 2022).

Whilst it is clear that recycling vesicles serve to deliver transmembrane cargoes such as integrins to the plasma membrane at the leading edge to facilitate cell migration, and the presence of recycling endosomes correlates with protrusion formation, whether recycling endosomes directly modulate F-actin regulation at the cell periphery to promote protrusion formation is not known.

The causal relationship between the directed localization of recycling endosomes and cell invasion could be dissected through active manipulation of their repositioning. Optogenetic control of Rab11 endosomes provided valuable insight into the role of Rab11 in neurite outgrowth in 2D (van Bergeijk *et al*., 2015). However, this approach depends on recruiting engineered cytoskeletal motor proteins such as kinesin, myosin, or dynein which are guided by an endogenous, polarized cytoskeleton. Blue light photo-toxicity, inadvertent background dark-state dimerisation binding, and motor overexpression side-effects are also confounding factors for this approach (Hoogenraad *et al*., 2001; Nijenhuis *et al*., 2020). In contrast to optogenetics, magnetic manipulation techniques are readily compatible with opaque specimens, are non-invasive and not limited by molecular motor properties (Keizer *et al*., 2022). Recent advances in magnetogenetics have led to the development of magnetic semisynthetic superparamagnetic nanoparticles combined with GFP that possess appropriate biological, physicochemical, and magnetic properties. These nanoparticles have demonstrated the ability to manipulate proteins and organelles fused to anti-GFP nanobodies inside living cells in 2D conditions with exceptional spatial and temporal resolution (Kappen *et al*., 2024; Keizer *et al*., 2022; Liße *et al*., 2017). Here, we have built on this approach to develop a magnetogenetic strategy to micromanipulate oncogenic Rab25 endosomes and vesicles with high spatiotemporal precision in migrating cells. We show that the local positioning of Rab25 vesicles to the plasma membrane is sufficient to initiate protrusion in 2D and physiologically relevant 3D matrices. The actin polymerising protein FMLN1 is a cargo of Rab25 vesicles which promotes the formation of both actin tracks and filopodia to generate new protrusions, and Rab25 acts to coordinate the delivery of FMNL1 and integrins with RhoA activity to promote protrusion formation.

## Results

### A magnetogenetic approach for remote manipulation of Rab25 endosomes demonstrates a direct role for endosome positioning in cell protrusion formation

To gain insight into the role of Rab25 in cancer cell invasion, we reactivated expression of Rab25 from the genomic locus of A2780 ovarian cancer cells using the DExCon approach (fig. S1A; (Gemperle *et al*., 2022)). This allows for the reactivation of silenced Rab25 gene in a doxycycline (dox)-dependent manner and enables the visualization of endogenous Rab25-positive endosomes using mCherry fluorescence. The anti-GFP nanobody (NB^GFP^) was included in frame with endogenous Rab25 to enable vesicle manipulations (fig. S1A-C). The functionality of NB^GFP^ was verified through colocalization with co-expressed GFP (fig. S1C), with induced Rab25 showing the expected perinuclear vesicle-enriched localization. Rab25 expression increased cell length, consistent with previous observations (Caswell *et al*., 2007), thereby demonstrating the functionality of our system (fig. S1DE). Additionally, we observed a positive correlation between the trafficking of Rab25 endosomes towards the plasma membrane (PM) and actin polymerisation/protrusion growth (see fig. S1FG). Similarly, new actin tracks were observed alongside Rab25 endosomes (fig. S1H), which also formed independently of microtubules (fig. S1I). These findings corroborate previous research (Caswell *et al*., 2007; Paul *et al*., 2015; Schuh, 2011), but do not distinguish the causality between endosome positioning and actin polymerisation/protrusion formation.

To determine the precise connection and causality between Rab25 recycling endosomes and protrusion growth, we developed a magnetogenetic approach for spatiotemporal control of Rab25 endosomes (Fig. 1A) using a magnetic tip attached to a remote-controlled micromanipulator (Fig. 1B). A2780 cells were microinjected with the superparamagnetic synthetic maghemite core particles coated with green fluorescent protein (mEGFP) fused with the iron binding site of Mms6 from magnetotactic bacteria (GFP-MNPs; Fig. 1A-C; (Kappen *et al*., 2024)). We previously found that these nanoparticles yield intracellular stealth properties and high magnetization, while also maintaining a small hydrodynamic diameter (19.6 ± 2.7 nm), allowing them to diffuse freely in the cytoplasm (Kappen *et al*., 2024). To enhance the usability and stability of the mEGFP coating (Kappen *et al*., 2024), mEGFP was cross-linked with paraformaldehyde after binding to nanoparticles. A rapid attraction gradient of freely moving GFP-MNPs towards the magnetic field was observed in all microinjected cells up to approximately 200 µm away from the micro-magnet covering the 120° angle (fig. S2A). The magnetic force exerted on particles at a distance of 30 to 150 µm from the micro-magnet was in the femto-Newton range, as previously determined using the Boltzmann law from the steady-state profile of GFP-MNPs distribution inside living cells (fig. S2B). By moving the micromagnet, it was possible to quantitatively concentrate GFP-MNPs to various subcellular, even centrally located positions in the cell (fig. S2C; Movie S1). Although GFP-MNPs have unhindered mobility in the cytoplasm (Kappen *et al*., 2024), they could not pass through the nuclear envelope and the nucleus provided an obstacle for their movement (fig. S2C; Movie S1). Microinjection of GFP-MNPs into the cytoplasm of cells stably expressing NB^GFP^-mCherry (control) resulted in the accumulation of NB^GFP^-mCherry in the cytoplasm. Magnetic relocalization of both GFP-MNPs and NB^GFP^-mCherry was found to be reversible within a matter of seconds (attraction time t_1/2_ ∼20 s; relaxation time t_1/2_ ∼10 s) (fig. S2D) and did not have any significant visible effect on the plasma membrane or F-actin despite sustained attraction of GFP-MNPs over a 10-60 min imaging period (Fig. 1CD; fig. S2EF).

**Figure 1.**
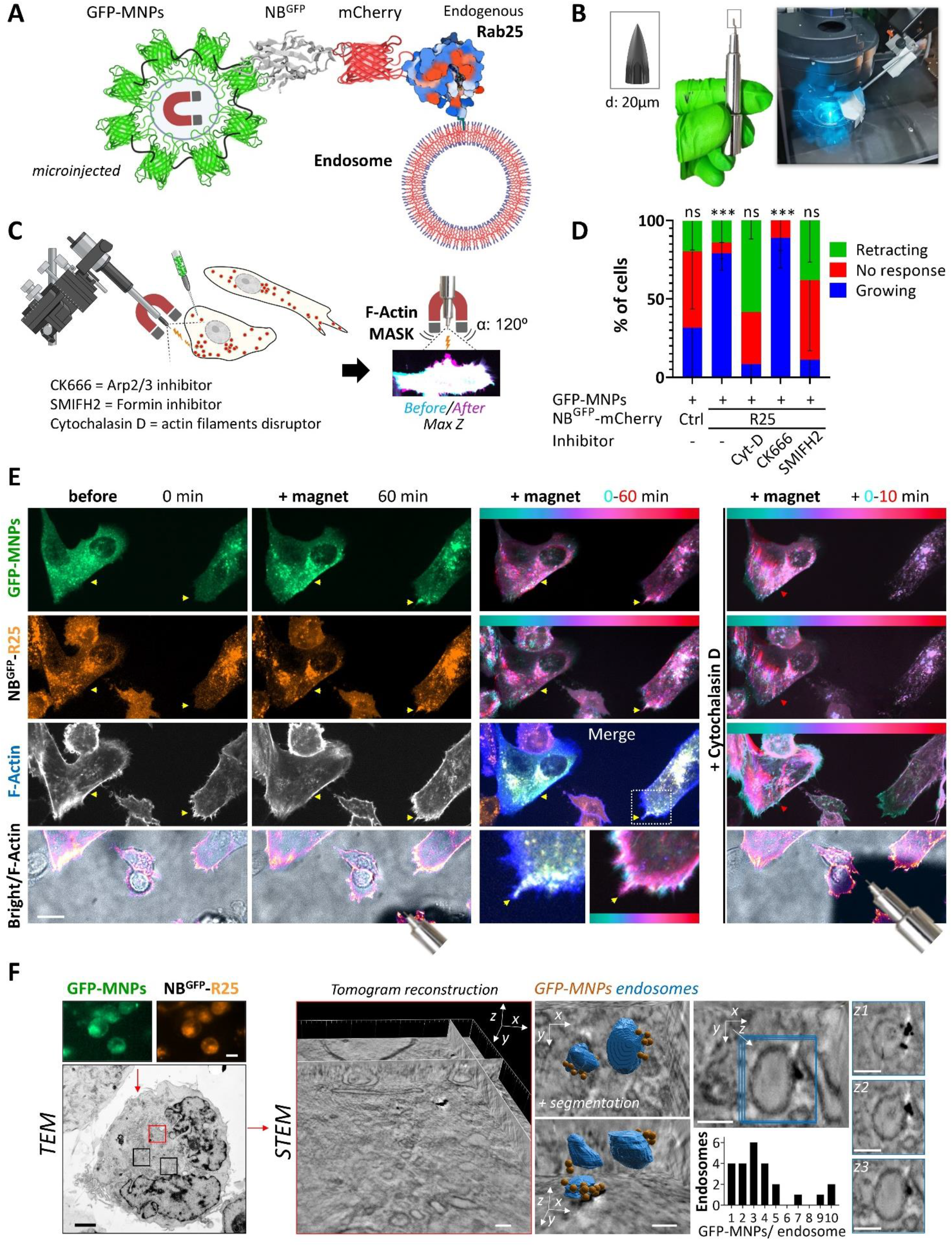
A magnetogenetic approach for remote manipulation of Rab25 demonstrates a direct role for endosome positioning in cell protrusion growth. **A)** Schematic diagram of magnetogenetic strategy to control Rab25 (R25) endosomes. **B)** Magnetic tip assembly composed of neodymium magnets and a fine wire tip attached to micromanipulation apparatus. **C)** Experimental set-up and **D)** quantification of protrusion growth: GFP-MNPs delivered by microinjection into randomly selected (independent of initial cell polarity) A2780 cells expressing control NB^GFP^-mCherry (Ctrl) or NB^GFP^-mCherry-Rab25 cells (DExCon-modified; Dox>94 h; 250 ng/ml) on FN-coated coverslips and MNPs attracted with magnetic tip. For each GFP-MNPs positive cell protrusion growth towards magnetic tip was analysed from maximum intensity projections (MIPs) of Lifeact-iRFP670 (F-actin) images; overlay masks before (0) and after (10-60 min) visible magnetic enrichment of GFP-MNPs, n = 19 (Ctrl; N = 5), n = 27 (R25; N = 8), n = 9 (R25; 200 nM Cytochalasin D; N = 2), n = 8 (R25; 100 μM CK666 N = 3), n = 13 (R25; 5 μM SMIFH2 N = 3), mean ± SD. Ordinary one-way ANOVA (GraphPad); ***P < 0.001. **E)** Representative spinning-disk confocal timelapse images of cells with magnetically attracted Rab25 endosomes before and after cytochalasin D treatment (see movie S2). Shadow in brightfield indicates magnetic tip. Yellow arrowheads and cyan-red LUT illustrates changes in GFP-MNPs and vesicle distribution and protrusion growth/filopodia (F-actin) over time through colour grading. Scale bar 20 μm. **F)** STEM tomography. GFP-MNPs delivered by electroporation (representative fluorescent images; scale bar 10 μm), TEM image (scale bar 2 μm) with corresponding STEM tomogram (red box, scale bar 2 μm)); zoom (right) with segmented tomogram and quantification of GFP/MNPs per recycling endosome [50-200 nm], scale bar 100 nm. Tomogram animation movie S16 accessible via https://doi.org/10.6084/m9.figshare.22155083. **A, C)** Created with BioRender.com

Next, GFP-MNPs were microinjected into cells engineered to express NB^GFP^-mCherry-Rab25 from the endogenous locus. GFP-MNPs localised to Rab25 endosomes, which were then re-localized using magnets. The re-localization was noticeable within several minutes and increased slowly over time (Fig. 1E; Movie S2). This suggests that the majority of endogenous Rab25 is likely to be vesicular, and not rapidly diffusible, consistent with previous observations (Gemperle *et al*., 2022). The direct re-localization of Rab25 resulted in the formation of small but robust and clearly discernible actin-rich protrusions at sites where vesicles had accumulated. This occurred in the direction of the magnetic field in the majority of microinjected cells, regardless of their polarity prior to positioning of the magnet (Fig. 1C-E; Movie S2). The effect of Rab25 on actin protrusions was blocked by cytochalasin-D or the small molecule inhibitor of formins (SMIFH2), but not by the Arp2/3 inhibitor CK666 (Fig. 1C-E; Movie S2; fig. S3AB). This suggests that the mechanism is dependent on formin-mediated actin polymerisation rather than Arp2/3. Unbiased AI-based quantification of un-manipulated vesicle dynamics revealed that the active movement of Rab25 endosomes in A2780 cells does not require Arp2/3 activity, as demonstrated by the CK666 inhibitor after 5 minutes of treatment (fig. S4A). In contrast, inhibition of formins by SMIFH2 had a negative impact on the mobility of Rab25 vesicles in the absence of magnetic manipulation (fig. S4B). However, magnetic relocalisation of Rab25 endosomes remained possible, although this did not result in any discernible effect on actin-based protrusions (fig. S4C). This is consistent with previously reported dose-dependent inhibition of A2780 cell migration by SMIFH2 (Hetmanski *et al*., 2021).

To our knowledge, the number of Rab molecules per recycling endosome has not been well documented. Therefore, to determine the average magnetic force exerted on Rab25-recycling endosomes (40-200 nm in size) and compare it with the force parameters related to molecular motors, we imaged GFP-MNPs that were delivered to Rab25 positive cells using scanning transmission electron microscopy (STEM) tomography. In the reconstituted and segmented tomograms, we observed up to 10 GFP-MNPs per recycling endosome usually enriched in a defined subdomain of the outer vesicular membrane (Fig. 1F). This observation indicates that Rab25 may not be distributed uniformly throughout the recycling endosome. Furthermore, it suggests that the total force per whole Rab25 endosome is up to 15 femto-Newtons (10 × 1.5 fN) at a distance of 50 µm from the micro-magnet which is two orders of magnitude below the proposed force of myosin and kinesin molecular motors (∼1-5 pN; (Cross, 2006; Schnitzer *et al*., 2000; Shukla *et al*., 2022)). Therefore, we hypothesized that a low but sustained magnetic force could alter the balance between positively and negatively directed motors and potentially repeatedly shift vesicles toward the micro-magnet when they temporarily detach from the cytoskeleton. This is consistent with our previously published findings, which demonstrated relatively fast relocation of genetically modified Rab25 endosomes to the outer mitochondrial membrane using knock-sideways (t_1/2_ = 476 s; (Gemperle et al., 2022)) and recent observation that Rab11 vesicles move to significant extent by passive diffusion/flow (Sittewelle and Royle, 2023).

Overall, these data demonstrate for the first time that it is feasible to actively re-localize membrane-bound Rab25 using magnets and that re-localization of Rab25 directly triggers protrusion outgrowth that is F-actin-and formin-dependent.

### Endosomes spatiotemporally modulate Rab25 localization to control protrusion outgrowth

To investigate Rab25’s role in inducing protrusion, we generated mutants of Rab25 that disrupted membrane anchorage (dC; fig. S5A) and/or mutated the GTP binding site (T26N; dominant negative (DN)). These NB^GFP^-mCherry fused Rab25 mutants (wt; dC; DN; or DN/dC) were introduced into A2780 cells (endogenous Rab25 expression non-detectable) using lentiviruses and sorted for comparable expression of mCherry. In comparison to the wild-type Rab25 (wt) protein, Rab25 dC, DN, and DN/dC mutants were more diffusely localized, suggesting a defect in the membrane recruitment and trafficking function for all these mutants (Fig. 2A). This observation is analogous to that previously demonstrated for Rab11b (Schlierf *et al*., 2000). GFP-MNPs were then microinjected to compare our ability to spatiotemporally control Rab25 variants using magnets. We found that Rab25 dC has fast and reversible re-localization kinetics, similar to those of the NB^GFP^-mCherry control (attraction time t_1/2_ ∼20s; relaxation time t_1/2_ ∼10s; Fig. 2BC). In contrast, membrane bound Rab25 wt yielded typical values of t_1/2_ ∼352 s with low reversibility (Fig. 2BC), consistent with previous optogenetic experiments (Nijenhuis *et al*., 2020). The quantification of the difference in re-localization kinetics between Rab25 dC and Rab25 wt provides further confirmation that it is possible to magnetically re-localize Rab25 endosomes, rather than just membrane-free Rab25. Additionally, the magnetic re-direction and enrichment of Rab25 wt and Rab25 dC, but not inactive Rab25 dC/DN, to the cell periphery promoted F-actin protrusion growth (Fig. 2DE; Movie S3). This suggests that the effect of Rab25 on protrusion, once localized, is direct and requires Rab25 activity, but at least on 2D substrates, vesicle association is not essential. Interestingly, a small proportion of Rab25 dC mutant associated with stress fibers (Fig. 2A), This was particularly evident in approximately 25 % of cells upon GFP-MNPs delivery by microinjection, and was not amenable to magnetic force, suggesting highly stable association (fig. S5C).

**Figure 2.**
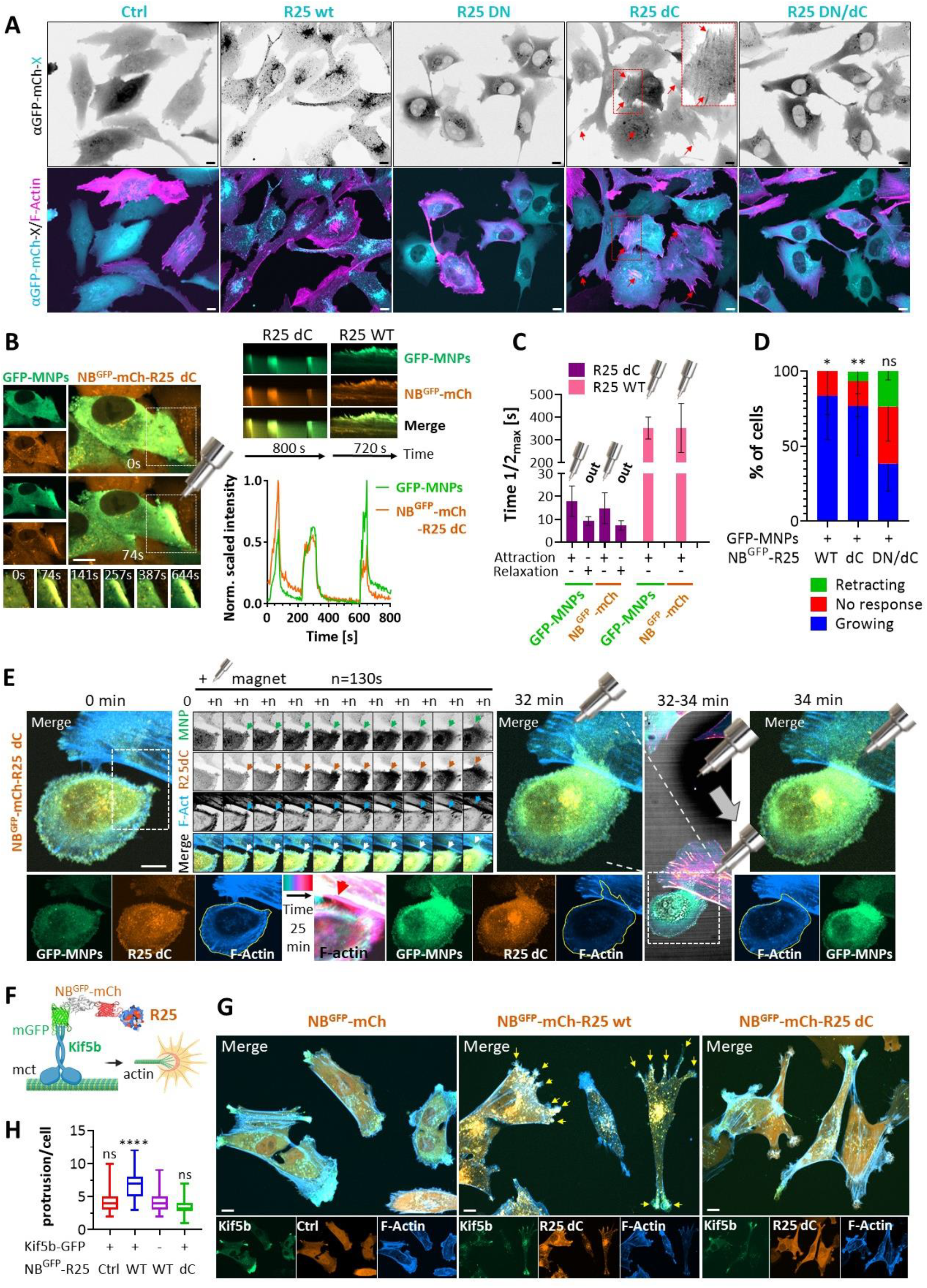
Endosomes spatiotemporally modulate Rab25 localization to control protrusion outgrowth. **A, B, E)** Representative images from confocal spinning-disk live cell imaging of A2780 stably expressing Lifeact-iRFP670 (F-actin) with NB^GFP^-mCherry-(Ctrl) or fused with different Rab25 mutants on FN. A) Boxed area ×2 magnified images. Arrows indicate stress fibers. Scale bar 10 μm. **B-C)** Magnetic attraction and release kinetics of NB^GFP^-mCherry-(X) and GFP-MNPs (delivered by microinjection); individual representative frames (scale bar 10 μm) from highlighted white box, kymographs and profiles determined from the changes in fluorescence intensity across black arrow (normalized 0-1 scaled fluorescent intensities shown), exponentially fitted and quantified in **C)**; n = 3 (N = 3). **D)** Protrusion growth quantified as in Fig. 1C, D; n = 14 (R25 WT; N = 3), n = 14 (R25 dC; N = 5), n = 22 (R25 DN/dC; N = 5), mean ± SD. Ordinary one-way ANOVA (GraphPad). **E)** Representative spinning-disk confocal timelapse images of cells on FN with magnetically attracted NB^GFP^-mCherry-Rab25 dC shown as MIPs, zoom inset from white box (1Z plane; arrows indicate changes). Shadow in brightfield indicates magnetic tip movement indicated by grey arrow and the corresponding alteration in cell shape outlined by yellow line. Colour-grade timelapse (cyan-red LUT, arrow). See movie S3. Scale bar 10 μm. **F)** Schematic diagram of alternative strategy to control Rab25 (R25) endosomes based on NB^GFP^ module and GFP-tagged C-terminally truncated molecular motor Kif5b [1-807] (created with BioRender.com). G) Representative confocal spinning-disk live cell images of A2780 on FN stably co-expressing Lifeact-iRFP670 (F-actin), NB^GFP^-mCherry-(Ctrl) or fused with different Rab25 mutants and GFP-Kif5b[1-807] sorted for as shown in fig. S6C (dox 48 h, 500 ng/ml). MIPs are shown. Scale bar 10 μm. Quantified in H) as the number of large protrusions per cell (yellow arrows, example). Anova on ranks, Dunn’s test (compared to variant without Kif5b); n > 50 cells (N = 3). \**P* < 0.05; \*\**P* < 0.01; \*\*\**P* < 0.001; \*\*\*\**P* < 0.001.

To compare our magnetogenetic results using an independent approach, we developed an additional strategy based on our NB^GFP^ module and GFP-tagged molecular motor Kif5b, which has previously been shown to associate with Rab11 vesicles (Matsuzaki *et al*., 2011). GFP-fused Kif5b [1-807] was truncated to prevent endogenous cargo binding (van Bergeijk *et al*., 2015) and used to semi-directly manipulate Rab25 variants to the cell periphery via NB^GFP^:GFP interaction (Fig. 2F). Stable expression of Kif5b[1-807]-GFP enriched NB^GFP^-mCherry control or fused variants of Rab25 in the cell periphery, and in combination with Rab25 wt and dC, but not control, induced an increase in cell length (fig. S6AB). Rab25 wt co-distributed with the filopodia and Rab25 dC with lamellipodia-like structures (fig. S6AB). These findings suggest that the major protrusions extending cell length increase are actin-dependent.

Interestingly, when the levels of GFP-KIF5b were closely matched in GFP-mCherry fusion-expressing cells (fig. S6C), a striking increase in the number of protrusions was observed in Rab25 wt-expressing cells, while no such increase was observed in the control or Rab25 dC cells (Fig. 2GH). This could suggest that Rab25 wt-induced protrusions are able to form and become stabilised more readily, perhaps due to delivery of adhesion receptor co-cargos. The increase in protrusion formation was accompanied by a significant reduction in the proliferation rate of Rab25 wt/Kif5b co-expressing cells (fig. S6D).

Taken together these data demonstrate that the positioning of Rab25 at the cell periphery is sufficient to drive the extension of actin-rich membrane protrusions from the cell periphery, but non-targeted delivery to the cell periphery via motor proteins could be detrimental to localized protrusion formation and other Rab25-mediated signalling processes such as control of proliferation. We further suggest that magnetogenetics is superior to methods that rely on recruiting engineered cytoskeletal motors, which suffer from overexpression side-effects and are limited by the cellular distribution of the engineered molecular motors and their polarised movement.

### Manipulation of Rab25 recycling endosomes in 3D matrix triggers robust F-actin protrusion

Next, we wanted to replicate our results under more challenging but physiological conditions using cell-derived matrices (CDM; (Kaukonen *et al*., 2017). In this 3D environment A2780 cells exhibit increased migration and front-rear polarization (Fig. 3A; (Hetmanski *et al*., 2019)), and form invasive protrusions (Caswell *et al*., 2007; Caswell and Zech, 2018; Paul *et al*., 2015). A gradient of mCherry protein was magnetically controlled through the interaction of NB^GFP^:GFP-MNPs without any noticeable impact on actin dynamics in control cells (fig. S7A). Upon expression of DExCon-modified NB^GFP^-mCherry-Rab25, the proximity of the magnetic tip to GFP-MNPs microinjected cells resulted in an unanticipated temporary cell contraction adaptation (Fig. 3AB; Movie S4). This resulted in a pulling force being exerted on surrounding matrix fibers (Fig. 3AB; fig. S7B). The cell adaptation was reproducibly noticeable when the magnet was moved into the field of view, but only for NB^GFP^-mCherry-Rab25 expressing cells. When cells were treated for 30 minutes with blebbistatin, no adaptation and matrix perturbation was observed (fig. S7B). This suggests that the actomyosin cytoskeleton is involved in the mechanosensing of magnetic force by MNP-bound Rab25 endosomes.

**Figure 3.**
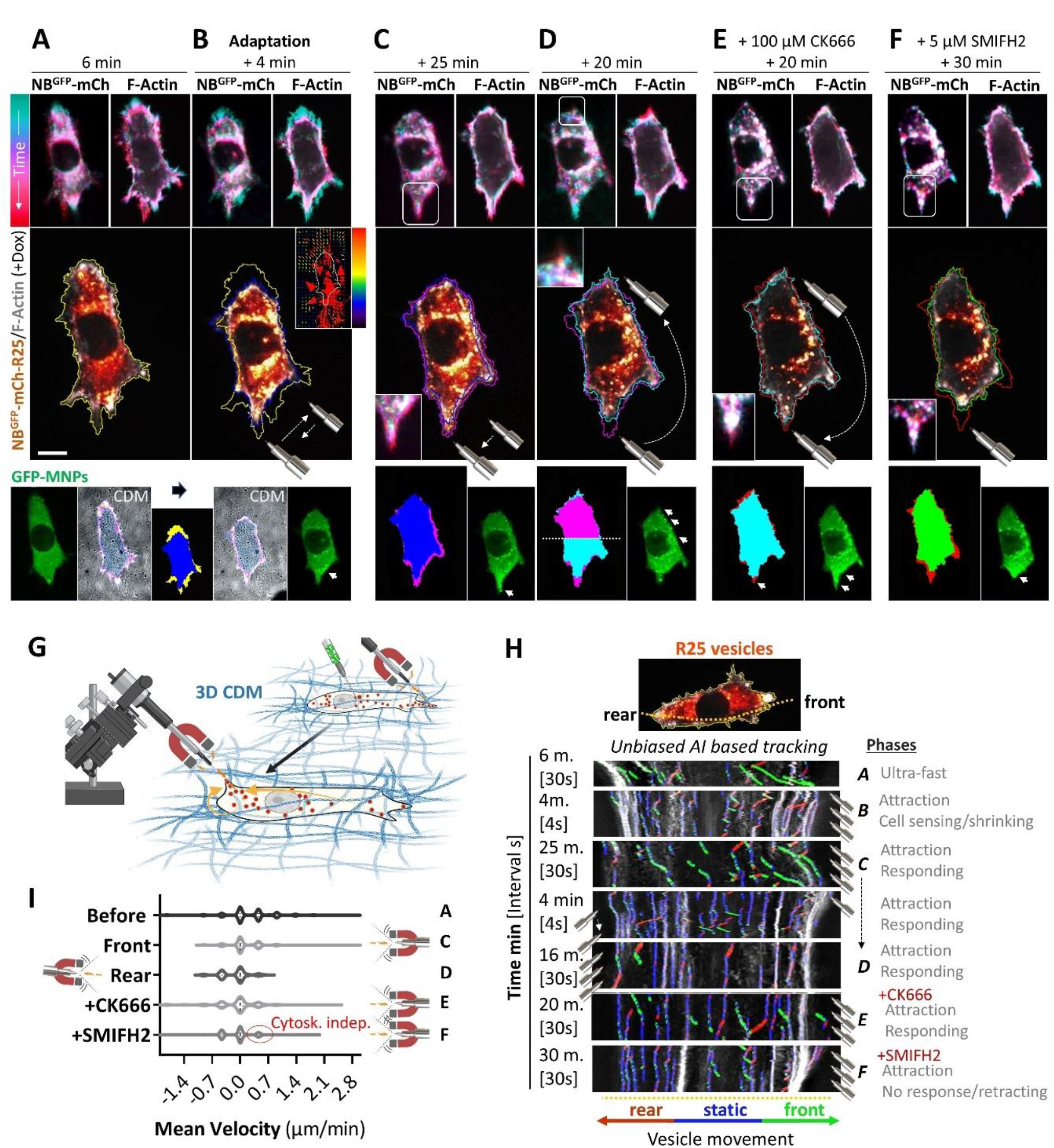
Manipulation of Rab25 recycling endosomes in living cells migrating in a 3D matrix triggers robust F-actin protrusion. **A-H)** Representative images from confocal spinning-disk live cell imaging. A2780 DExCon-modified NB^GFP^-mCherry-Rab25 cells (see fig. S1) dox pre-treated (72 h; 250 ng/ml) expressing Lifeact-iRFP670 (F-actin) migrating in 3D CDM. GFP-MNPs delivered by microinjection and repeatedly relocalized using a magnetic tip (movement specified) as depicted in **G)** in the presence or absence of different indicated inhibitors. See movie S4. N = 3. **A-F)** Masks created from F-actin images (solid colours overlaid from preceding and following event). Arrowheads, gradient of GFP-MNPs. Colour-grade (cyan-red) LUT images show changes in vesicles (NB^GFP^-mCh-R25) / GFP-MNPs distribution and cell shape (F-actin) in time. Relative stress map (inset B) is displayed using as a vectorial plot with red-blue LUT, size and direction of vectorial arrows illustrate force distribution exterted on CDM. Scale bar 20 μm. **H-I)** AI based tracking of vesicle (NB^GFP^-mCh-R25) movement (labelled blue, static; brown, towards rear; green, towards front) using kymographs (generated from yellow dashed line) and Kymobutler plugin during experiment shown in **A-F)**. Note: Ultra-fast phase kymograph (A) of vesicle movement was corrected by eye. The position of magnet relative to the cell front and rear is showed. Total quantification of H) is shown in **I**).

Cells adapted to the remote mechanical perturbation within approximately 5 minutes. Subsequently, MNP-bound Rab25 endosomes were attracted towards the magnet, and near-simultaneously, protrusion growth was observed within a region of the cell that had been protruding (dark blue to magenta outlines, Fig. 3C). Strikingly, repositioning of the magnet toward the cell rear led to Rab25 vesicle accumulation and actin protrusion formation in the posterior part of the cell which previously lacked protrusive structures (magenta to cyan outlines, Fig. 3D, G; Movie S4). The repositioning of the magnetic tip to its original position promoted vesicle movement accompanied by actin-based protrusion in a previously retracting region of the cell, even in the presence of the Arp2/3 inhibitor CK666 (cyan to red outlines, Fig. 3E). In contrast, addition of formin inhibitor (SMIFH2) caused retraction of the protrusion, suggesting dependence on the formin family of actin polymerising proteins (red to green outlines, Fig. 3F; Movie S4). The unbiased AI-based vesicular tracking confirmed that local actin polymerisation near the cell periphery (Fig. 3A-F; top right F-actin colour-grade timelapses) follows the relocalization of Rab25 endosomes that are controlled by magnetic tip movement (Fig. 3H-I). SMIFH2 treatment disrupted this connection between Rab25 and F-actin protrusion outgrowth (Fig. 3H-I). We then similarly tested the exogenously expressed Rab25 dC mutant with impaired membrane recruitment. However, despite rapid Rab25 dC attraction via magnetic repositioning of GFP-MNPs, we were unable to induce protrusions/PM growth in any of four biological repeats (fig. S7C).

We further confirmed our observation by an indirect approach, scoring the extension of pseudopodial protrusions at the cell front, previously associated with Rab25 expression in invasive cells (Caswell *et al*., 2007). The reactivation of endogenous Rab25 or expression of exogenous Rab25 wt correlated with an extension in pseudopodial protrusions in front of invasive cells and an increase in cell migration persistence. No change in average speed was observed (fig. S8A-E), which is consistent with previous findings (Caswell *et al*., 2007; Gemperle *et al*., 2022). Nevertheless, the expression of dominant negative Rab25 (DN) or Rab25 uncoupled from endosomes (Rab25 dC) did not result in an increase in pseudopod length or cell migration persistence (fig. S8A-E). To enhance the redistribution of Rab25 mutants to the cell periphery and to partially rescue the diffuse localisation of the Rab25 dC variant, we again stably co-expressed Rab25 variants fused to NB^GFP^-mCherry and Kif5b[1-807]-GFP. For Rab25 wt, Kif5b expression resulted in a striking increase in pseudopod extension at the cell front, to a significantly greater extent than Rab25 wt alone (fig. S8A-C). This effect was accompanied by high migration persistence (fig. S8D-E). Notably, this phenomenon was not observed in the control or in the Rab25 dC mutant, in contrast to the effects observed on cell elongation on 2D substrates. Formin inhibition (SMIFH2) blocked the ability of Kif5b-Rab25 wt to induce pseudopod extension in a dose-dependent manner (fig. S8F), consistent with previous results.

The inability of the Rab25 dC mutant to promote membrane/protrusion growth under 3D conditions, despite peripheral enrichment by kinesin motors or magnetogenetic targeting to the cell periphery, suggests that spatiotemporal control of Rab25 is not sufficient to control all aspects of protrusion formation. Rab25-driven tumor-cell invasion into fibronectin rich CDM is highly dependent on Rab25’s ability to interact with integrin β1-containing recycling vesicles (Caswell *et al*., 2007). Indeed, the elongation of pseudopodial protrusions promoted by Kif5b-dependent relocalisation of Rab25 endosomes could be inhibited by integrin β1 blocking antibodies, suggesting that in addition to formins, integrin β1 is also an important factor for protrusion formation (fig. S8F). Local targeting of GFP-MNPs:Rab25 endosomes to the cell periphery of cells in 3D CDM reproducibly induced protrusion growth, but this was completely abrogated by integrin β1 blockade (fig. S8G; Movie S5). These data suggest that delivery of integrin β1 containing vesicles to specific membrane sites is critical for Rab25-mediated protrusions in 3D CDM, but may be dispensable in 2D.

Taken together, these results demonstrate the critical role of Rab25 endosomes as a platform for the spatio-temporal control of Rab25 activity, coordinating modulation of actin polymerisation with integrin adhesion receptor delivery for protrusion formation in 3D matrix.

### Rab25 recycling endosomes recruit FMNL1

To identify Rab25 proximal interacting actin nucleators in living A2780 cells, we re-analyzed our previously published Rab25-BioID (proximity labelling) dataset (Wilson *et al*., 2023). In addition to known Rab25-associated membrane receptors (such as EGFR and integrin β1, motors (such as Kif5b), and motor coupling proteins (such as RAB11FIP2 which binds to MyoVb), we noted that one of the most highly enriched hits was the actin nucleator FMNL1, which belongs to the formin family (fig. S9A).

The use of confocal spinning-disk microscopy revealed the colocalisation of Rab25 and FMNL1 on perinuclear recycling endosomes, which were surrounded by abundant F-actin (Fig. 4A). Additionally, we found high colocalization of Rab25, FMNL1, and F-actin in actin-rich protrusions, including filopodia (fig. S9B). FMNL1:Rab25 remained highly colocalized upon inhibition of cytoskeletal elements/regulators, albeit with a slight decrease upon nocodazole treatment (fig. S9C-D). Interestingly, whilst pan-formin inhibition didn’t disrupt FMNL1:Rab25 association, it did decrease the association of both with F-actin. Notably, treatment with nocodazole or low-dose cytochalasin D disrupted the perinuclear Rab25 enrichment and resulted in a redistribution of FMNL1 and F-actin towards the cell periphery (fig. S9C-D).

**Figure 4.**
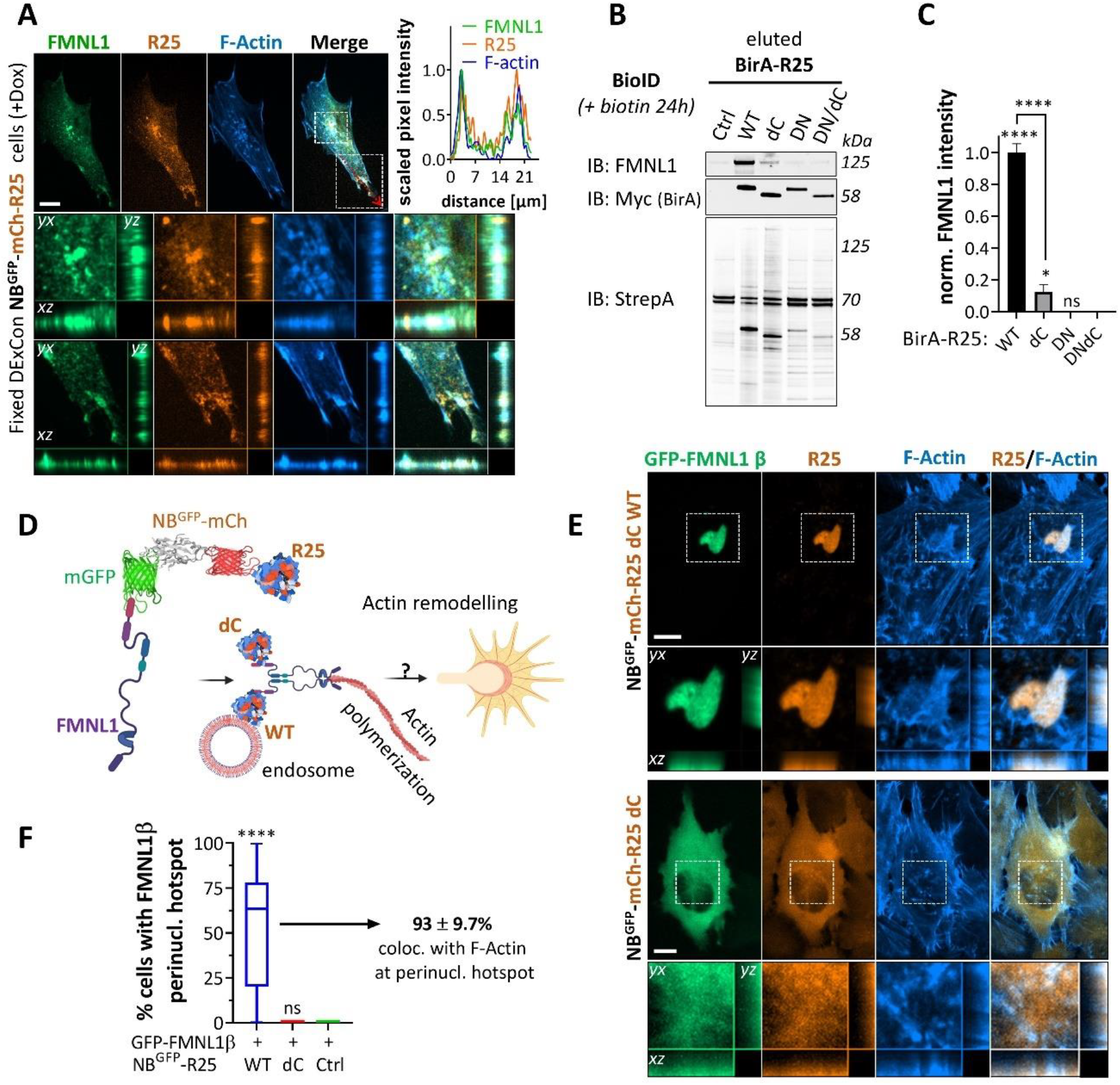
Rab25 recycling endosomes recruit FMNL1. **A)** Representative confocal spinning-disk images of A2780 DExCon-modified NB^GFP^-mCherry-Rab25 cell (mCherry, orange/amber) dox pre-treated (> 94 h; 250 ng/ml) immunolabeled for FMNL1 (green, rabbit anti-FMNL1, Alexa-488) and stained for F-actin (blue, Phalloidin-Alexa633). Red dotted line, line scan profile of normalized 0-1 scaled fluorescent intensities. Zoom inset from white box shown as MIP with cross-sections. FN-coated µ-Plate 96 plate (#1.5 IbiTreat). Scale bar 10 µm. **B, C)** BioID experiment. A2780 stably expressing BirA fused Rab25 WT or mutants (dC, DN, DN/dC) cultured with biotin (1 μM biotin, 16 h). Lyzates equalized to total protein amount and biotinylated proteins pulled down with Streptavidin beads (Ctrl, no BirA). **B)** Immunoblots of pulled down FMNL1, BirA (Myc epitope), Streptavidin (StrepA) as loading control. Fluorescent antibodies shown as black and white. **C)** Quantification of pulled down FMNL1 from immunoblots. The graph shows band intensities normalized to individual BirA levels and relative to BirA-R25 WT values (background from ctrl sample subtracted). Mean ± SD; N = 3. One-way ANOVA analysis Tukey post hoc test (compared to DNdC or as indicated); \**P* < 0.05; \*\*\*\**P* < 0.001. **D)** Schematic diagram of strategy to enforce the proximity of FMNL1β with Rab25 WT or dC mutant unable to bind endosomes by the interaction of GFP(-FMNL1) and NB^GFP^(-mCherry-Rab25). Created with BioRender.com. **E)** Representative confocal spinning-disk live cell images of A2780 stably co-expressing Lifeact-iRFP670 (F-actin), GFP-FMNL1β with NB^GFP^-mCherry-(Ctrl) or fused with different Rab25 mutants. Zoom inset from white box shown as MIP with cross-sections. FN-coated 96 well plate (cellvis, #1.5H cover glass). Scale bar 10 µm. Quantified in **F)** as % of cells with FMNL1 β perinuclear hotspot as shown in **E)**. Boxplot show the median, 25th, and 75th percentile with whiskers reaching the last data point; n > 55 cells/condition; N = 3. One-way ANOVA analysis Tukey post hoc test (compared to Ctrl); \*\*\*\**P* < 0.001.

To characterize FMNL1:Rab25 interaction in live cells we used a knock-sideways re-localization approach, whereby Rab25 fused to FKBP is re-localized (t_1/2_∼ 5.4 min ±0.1; (Wilson *et al*., 2023)) to the outer mitochondrial membrane upon rapamycin treatment via interaction of FKBP:FRBmito (fig. S9E). GFP-FKBP-Rab25 or GFP-FKBP were highly enriched to mitochondria upon rapamycin treatment, and there was a subtle redistribution of FMNL1 with GFP-FKBP-Rab25, suggesting that the Rab25:FMNL1 interaction is weak, transient or that re-distribution is limited by other factors (fig. S9F-G). To determine if Rab25’s association with FMNL1 is restricted to endosomes and/or to active Rab25, we conducted additional BioID experiments using Rab25 wt and Rab25 mutants (dC, DN, DN/dC). Interestingly, whilst Rab25 wt-induced extensive proximal biotinylation of FMNL1, inactive dominant negative forms of Rab25 did not (Fig. 4BC). However, Rab25 dC was also able to do so, albeit to a far lesser extent (Fig. 4BC). This suggests that endosomal localization is beneficial for Rab25 association with FMNL1, but not essential. The enrichment of Rab25 wt or Rab25 dC at the cell periphery by Kif5b[1-807]-GFP motor, via NB^GFP^:GFP interaction, showed strong co-localization with FMNL1 for both variants (fig. S9H). This suggests that Rab25 dC encounters FMLN1 at the plasma membrane rather than endosomes, hence lower association. Integrin β1 proximity labelling, however, was restricted solely to Rab25 wt, consistent with our previous findings that Rab25-GTP interacts with integrin β1 (fig. S9I). This indicates that endosomal localisation is required for Rab25 to interact with integrin β1.

Three isoforms of FMNL1 (α, β, γ) have been described, differing in the regulatory sequence at the C-terminus. Of these, FMNL1γ appears to be constitutively active (Han *et al*., 2009; Yayoshi-Yamamoto *et al*., 2000). We identified regulatable FMNL1β isoform as the major isoform in A2780 cells (fig. S10A). To provide evidence of FMNL1 activity and the functionality of the FMNL1β/Rab25 association in A2780 DExCon Rab25 cells, we enforced the proximity of FMNL1β and Rab25 by the interaction of GFP(-FMNL1) (Fig. 4D). This resulted in stable endosomal colocalization of Rab25, FMNL1 and F-actin (Fig. S10B). To support our hypothesis that endosomes locally regulate Rab25 activity to spatiotemporally control FMNL1-dependent actin nucleation, we generated stable cell lines co-expressing GFP-FMNL1β/NB^GFP^-mCherry-Rab25 wt or -Rab25 dC. The sustained association of FMNL1β with NB^GFP^-mCherry-Rab25, but not with diffuse Rab25 dC, resulted in striking co-enrichment of FMNL1β and Rab25 wt in a perinuclear area in approximately 50% of cells, with prominent actin nucleation at these sites in 93% of these cells (Fig. 4E-F). Overall, these results demonstrate that Rab25 recycling endosomes locally can change actin polymerisation via FMNL1, and that this association is likely to be transient in normal cells to maintain the F-actin network.

### Magnetic spatiotemporal control of Rab25 endosomes modulates F-actin polymerisation in protrusions via FMNL1

The rearrangement of the actin cytoskeleton to promote actin-based protrusions requires nucleation and elongation of actin filaments, a process that is catalyzed by actin assembly factors (Nürnberg *et al*., 2011). We inactivated the FMNL1 gene using CRISPR/Cas9-based gene editing in DExCon-modified NB^GFP^-mCherry-Rab25 cells. However, we were only able to generate heterozygote KOs of one allele (fig. S11A-D; for details see Methods). Therefore, we developed a combination strategy that efficiently depletes FMNL1 using a smart pool of siRNAs in the FMNL-/+ heterozygous background previously created by the CRISPR/Cas9 editing (Fig. 5A; fig. S11E-F). To demonstrate that Rab25 promotes protrusions via FMNL1 we then directly re-localized NB^GFP^-mCherry-Rab25 in cells using magnets. In 2D, we again observed formation of small but robust and clearly discernible actin-rich protrusions in the direction of the magnetic field in the majority of microinjected cells transfected with control siRNA, irrespective of their polarity (Fig. 5B top; 5C). By contrast, magnetic relocalization of NB^GFP^-mCherry-Rab25 towards the cell periphery did not induce protrusions in FMNL1 depleted cells (Fig. 5B bottom; 5C; Movie S6). In cells migrating in 3D, robust generation of protrusions was reproducibly induced by magnetic relocalisation of NB^GFP^-mCherry-Rab25 and FMNL1 expressing cells (Fig. 5D top; Movie S7). However, this phenomenon was not observed following FMNL1 depletion, demonstrating that Rab25 endosomes directly promote actin rich protrusions via FMNL1 (Fig. 5D bottom; Movie S8). Similarly, only in FMNL1 expressing cells, we were occasionally able to directly manipulate endosomes in a slowly moving cluster that induced the formation of a visible actin polymerisation hot-spot. This actin hot-spot moved and highly colocalized with Rab25 and GFP-MNPs over 60 min, co-attracted to the magnet, and returned after its removal (fig. S12) suggesting that FMNL1 regulate the formation of both actin tracks and filopodia to generate new protrusions.

**Figure 5.**
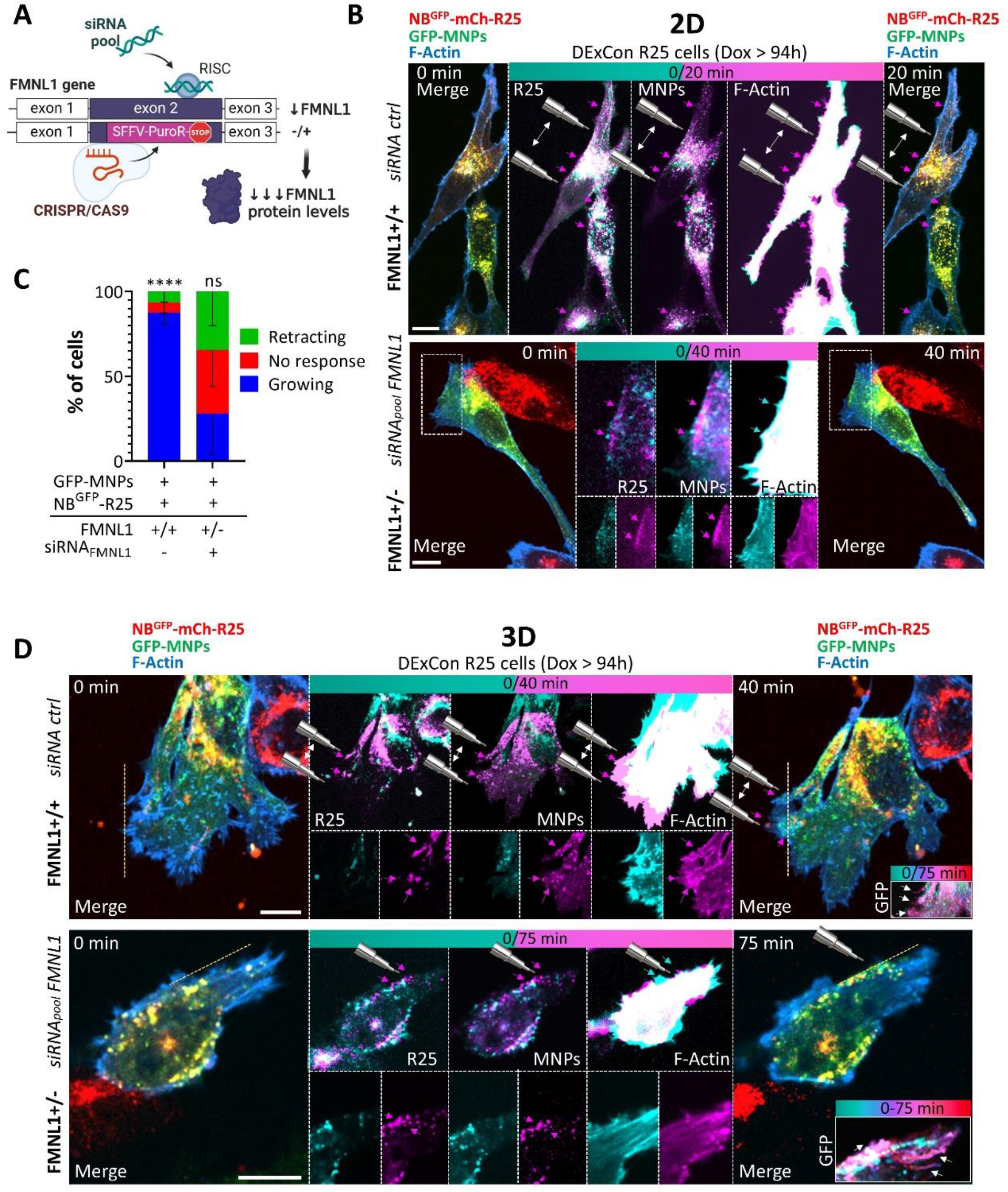
Magnetic spatiotemporal control of Rab25 endosomes modulates F-actin polymerisation in protrusions via FMNL1. **A)** Strategy to efficiently deplete FMNL1 using a smart pool of siRNAs in a FMNL+/-heterozygous background previously created by the CRISPR/Cas9 editing (see also fig. S11A-D; for details Methods). Created with BioRender.com. **B, D)** Representative frames from confocal spinning-disk live cell timelapse images of A2780 DExCon-modified NB^GFP^-mCherry-Rab25 FMNL1+/+ or FMNL1+/-cells (dox treated > 94h; 250 ng/ml) stably expressing Lifeact-iRFP670 (F-actin) on FN, microinjected with GFP-MNPs and nucleofected with chemically modified siRNA_pool_ anti-FMNL1 or non-targeting ctrl siRNA (see methods) as indicated. Experiments performed 5 days after (for immunoblot see fig. S11F). Rab25 endosomes attracted using magnetic tip (shown by cartoon, arrows indicate movement). Cyan arrows (time 0) or magenta (later timepoint) indicate the outcome. MIPs if not stated otherwise. Zoomed insets correspond to areas indicated by boxed areas. Scale bar 10 μm. Quantification in **C)** as in Fig. 1C, D as % of cells showing changes in protrusion growth towards magnetic tip; n = 12 (siRNA ctrl; N = 3), n = 14 (siRNA_pool_ anti-FMNL1; N = 3; see movie S6), mean ± SD. Ordinary one-way ANOVA (GraphPad); \*\*\*\**P* < 0.001. **D)** Representative images from confocal spinning-disk live imaging of cells in 3D CDM; N = 3. MIP (top) or 1 matched (0/40 min) Z plane shown (bottom). Dashed line, cell edge (F-actin) at the time 0. Colour-grade (cyan-red) LUT image show changes in GFP-MNPs distribution in time. See movies S7-8.

In summary, these results confirm our hypothesis that Rab25 endosomes promote F-actin polymerisation and protrusions by recruiting the formin FMNL1 in addition to integrin β1. Furthermore, this proof-of-principle data demonstrates the power of our approach using magnets in combination with genetic perturbation to demonstrate the causal effect of Rab25 endosome positioning on F-actin protrusions in invasive cancer cells.

### Rab25 recycling endosomes serve as signalling platform for RhoA/FMNL1 activity

Endosomes, including Rab11 recycling vesicles, are emerging platforms for integrin-mediated signalling that are linked with local RhoA activity to fine tune actomyosin-based contractility, thereby establishing front-rear cell polarity (Alanko and Ivaska, 2016; Eisler *et al*., 2018; Gaston *et al*., 2021; Jacquemet, Green, *et al*., 2013; Vassilev *et al*., 2017). FMNL1 has been recently identified as proximity interactor of RhoA (and of RhoB, RhoG, RhoD, RhoU, RhoF, RhoJ), binding to its active form (Bagci *et al*., 2020; Gomez *et al*., 2007). Since we identified a major FMNL1β isoform in A2780 cells, whose activity is likely regulated by Rho GTPases, we tested whether the activity of RhoA might be locally modulated by the magnetic relocalization of Rab25-positive endosomes. To monitor endogenous RhoA activity we adapted a localization-based activity probe iRFP670_3x_-RBD_4x_ that detects active RhoA with improved specificity and sensitivity (Mahlandt *et al*., 2021). We found that the active Rho probe iRFP670_3x_-RBD_4x_ is enriched with Rab25 at the perinuclear recycling compartment (PNRC) in a Rab25-dependent manner, both in 2D and 3D (Fig. 6AB; fig. S13AB), and in vesicles moving at the cell periphery (fig. S13C). This localisation was not perturbed with acute FMNL1 depletion (Fig. 6B). Furthermore, GFP-RhoA co-localised with mCherry-Rab25 in filopodia-based protrusions (fig. S13D), as well as with the Rho activity probe in the perinuclear region of cells (fig. S13E). This overlap was more pronounced upon expression of the constitutively active Q63L (CA) RhoA form, but not the dominant negative (DN) T19N RhoA mutant (fig. S13E).

**Figure 6.**
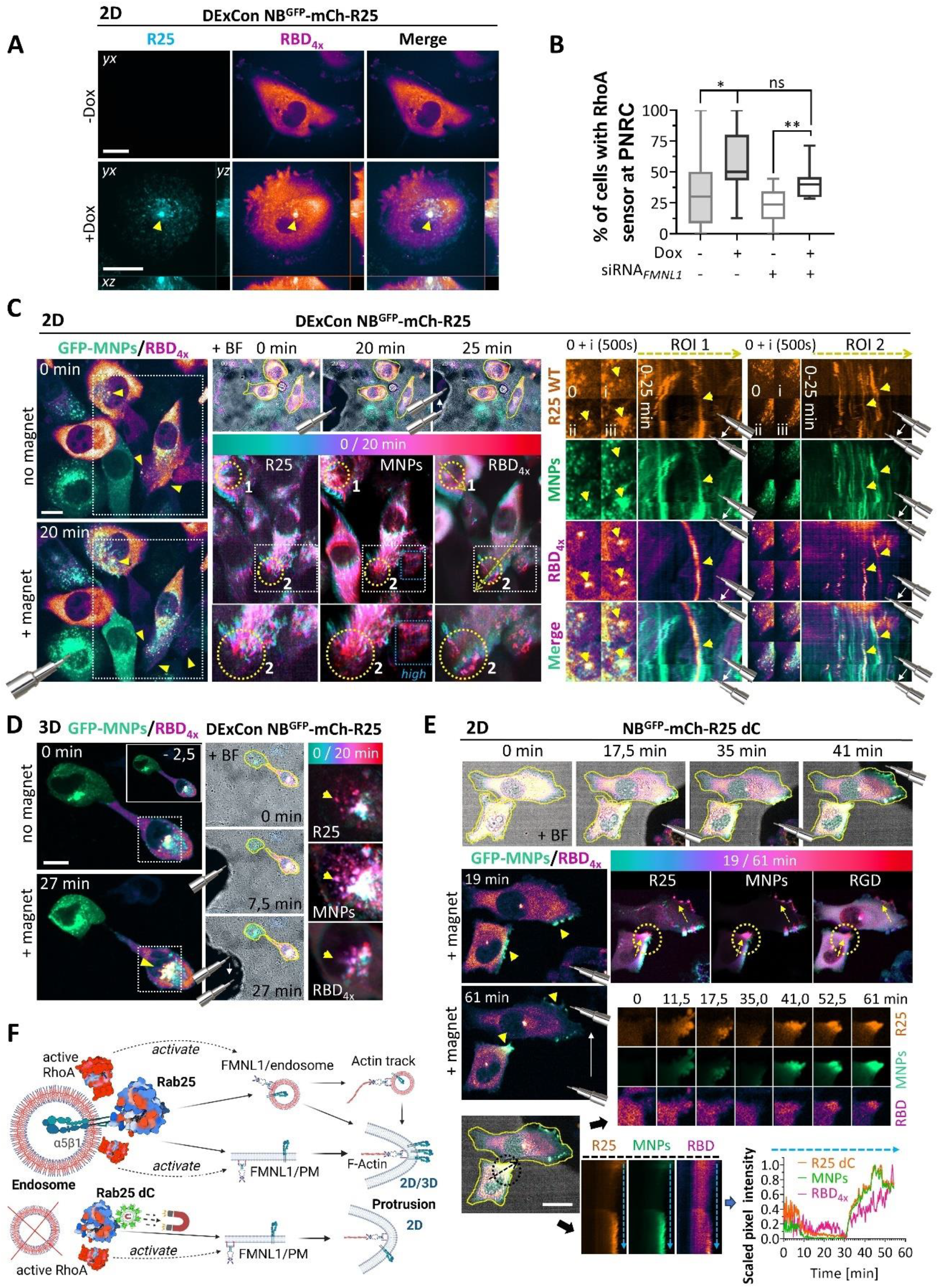
Rab25 recycling endosomes serve as signalling platforms for RhoA/FMNL1 activity. **A, C, D)** Representative images from confocal spinning-disk live cell microscopy of A2780 DExCon-modified NB^GFP^-mCherry-Rab25 (R25; dox treated if not stated otherwise > 72 h, 250 ng/ml) or E) A2780 NB^GFP^-mCherry-Rab25 dC expressing cells stably co-expressing active RhoA probe iRFP670_3x_-RBD_4x_ (RBD_4x_). GFP-MNPs microinjected (green). Colour-grade (cyan-red) LUT image show changes in mCherry/GFP-MNPs/RBD_4x_ in time (generated from one Z plane). Yellow arrowheads and arrows, mCherry/GFP-MNPs/RBD_4x_ signal enrichment/changes/movement. Position and movement of magnetic tip visible as shadow and depicted by cartoon (white arrows indicate movement). Cell shape outlined by yellow line in merged brightfield (BF) images. Scale bar 20 μm. A) MIPs with cross-sections, cells on FN. Arrow, PNRC. Quantified in **B)** as percentage of cells with fluorescent signal of RBD_4x_ visibly enriched at PNRC spot ± Rab25 (± dox 48h; n > 60 cells, N = 3) ± nucleofected with siRNA_pool_ anti-FMNL1 (n > 40 cells, N = 2). One-way ANOVA Tukey’s multiple comparison test; \**P* < 0.05; \*\**P* < 0.01. **C)** Cells on FN, one matched Z plane shown for all. See movie S9. White boxed areas, ×3 colour-grade LUT images for individual channels with indicated zoomed insets (bottom); blue box, higher constrast. Circle 1, zoomed inset ROI1 (time interval 500s = i). Circle 2 with yellow dotted line, zoomed inset ROI2 shown as kymograph. **D)** 3D CDM. GFP-MNPs/RBD_4x_ merge shown as MIPs. Top left Inset -2,5 min, no magnet. Movie S22 accessible via https://doi.org/10.6084/m9.figshare.22155083. **E)** FN-coated coverslip. One matched Z plane shown for all. See movie S10. Yellow circle, zoomed inset timelapse for individual channels. Black circle and black dashed line, zoomed inset shown as kymograph for individual channels. Blue dashed line across kymograph, line scan profile of normalized 0-1 scaled fluorescent intensities. **F)** Schematic model summarizing our results: Rab25 endosomes serves as a platform for spatio-temporal control of Rab25 activity which can be directly modulated using mangetogenetics. This control locally modulates RhoA activity to promote actin polymerisation in protrusions through an FMNL1-dependent actin nucleation mechanism, while simultaneously delivering integrin adhesion complexes for protrusion formation. Created with BioRender.com.

Giving the colocalization of Rab25 endosomes with RhoA and the active Rho probe (fig. S13C), we reasoned that magnetic relocalization of Rab25 towards PM could lead to dynamic co-redistribution of RhoA activity as well. Indeed, magnetic redistribution of NB^GFP^-mCherry-Rab25 correlated with local RhoA activity changes in cells moving in 2D and 3D (CDM) conditions (Fig. 6C, D; Movie S9). Sustained attraction of Rab25 endosomes resulted in an increased localisation of the active Rho probe in a punctate pattern in cell protrusions that were in close proximity to, but not completely superimposed with, Rab25 vesicles and GFP-MNPs (Fig. 6C, D; Movie S9).

Previous studies have suggested that α5β1 recycling can modulate RhoA activity through coordinated polarized signalling of receptor tyrosine kinases (RTKs), including EGFR1 and c-Met, as well as the activity of the RacGAP1-IQGAP1 complex (Caswell *et al*., 2008; Jacquemet, Green, *et al*., 2013; Muller *et al*., 2009). We therefore tested the ability of the Rab25 dC variant, which cannot associate with vesicles containing integrin β1 but can promote cell protrusion in 2D, to locally modulate RhoA activity. Magnetic spatio-temporal control of a NB^GFP^-mCherry-Rab25 dC gradient resulted in a rapid local increase in the intensity of the active Rho probe, which followed Rab25 dC enrichment towards the cell periphery or across the cell after redirection by the magnet (Fig. 6E; Movie S10). However, a punctate pattern of active Rho probe was not observed before or after relocalization in protrusions of Rab25 dC expressing cells suggesting that Rab25 can directly modulate RhoA activity, regardless of endosome association.

Taken together, our magnetogenetic approach demonstrates the importance of Rab25 endosomes as a platform for spatio-temporal control of Rab25-driven activities. This control locally modulates RhoA activity to promote actin polymerisation in protrusions through an FMNL1-dependent actin nucleation mechanism, while simultaneously delivering integrin adhesion receptors for protrusion formation and stabilisation in physiologically relevant 3D matrices (Fig. 6F).

## Discussion

The contribution of endocytic recycling to tumour progression and cancer cell invasion has been well documented. However, understanding of the precise mechanisms through which recycling vesicles can generate or maintain protrusions upon arrival at the plasma membrane has been hampered by an inability to directly manipulate vesicle positioning in time and space. Here, we developed a magenetogenetic approach for manipulation of vesicle positioning, by engineering the endogenous expression of the recycling regulator Rab25 such that it is expressed with an anti-GFP nanobody tag (NB^GFP^). This allows coupling of Rab25 vesicles to magnetic particles, and their relocalisation when placed within a magnetic field. We used this approach to demonstrate that positioning of Rab25 vesicles to the plasma membrane is sufficient to generate new F-actin protrusions in cancer cells. These protrusions are dependent on the actin polymerising protein FMNL1, and Rab25 coordinates the activity of RhoA with the arrival of FMNL1 and integrin β1 cargoes to form protrusions that promote invasive migration of cancer cell in 3D matrix.

Optogenetic approaches have been developed to manipulate endosome positioning, including via Rab11 family proteins (Nijenhuis *et al*., 2020; van Bergeijk *et al*., 2015). Light inducible dimerization was used to couple Rab11 to motor proteins kinesin or dynein which were able to promote or perturb the dynamics of neuronal growth cones respectively after illumination (van Bergeijk *et al*., 2015). However, these approaches rely on the coupling between the target protein and a motor attached to a cytoskeletal element, and hence relocalisation is therefore limited by the polarisation/orientation of the cytoskeleton. To overcome this limitation, we used a magenetogenetic technique, building on previous approaches (Bongaerts *et al*., 2020; Etoc *et al*., 2015; Kappen *et al*., 2024; Keizer *et al*., 2022; Liße *et al*., 2017; Steketee *et al*., 2011). Significantly, the non-specific manipulation of endolysosomes (where endosomes and lysosomes are loaded through fluid phase uptake of 991 ± 378 magnetic particles) was shown to bias cancer cell movement in 2D towards magnetic micropillars (reaching a force of 22 ± 16 pN per cell at 10-µm distance) (Bongaerts *et al*., 2020) and small shift in directional neurite outgrowth when cultured in a magnetic field gradient (Dhillon *et al*., 2022). Similarly, magnetic particles functionalised with TrkB agonist antibodies, delivered non-specifically into neurons by endocytosis, were able to stop neurite growth in response to a magnetic field (15 pN force; (Steketee *et al*., 2011)). For our purposes, microinjection of magnetic particles proved a suitable delivery method for loading of cells expressing NB^GFP^-mCherry proteins, and enabled specific manipulation of both cytosolic proteins and Rab25 coupled to endosomal membranes (Fig. 1, Movie S2; 2B-E, Movie S3), even in cells migrating in 3D matrix (Fig. 3A-G; Fig. 5D; fig. S7A-C; Movie S4 and S7-8). Interestingly, magnetic manipulation of endosomal proteins was slower than that of cytosolic proteins (Fig. 2C), suggesting that endosomal proteins are coupled to endosomes and to the cytoskeleton, and that magnetic redistribution may be enabled by transient detachment of endosomes from the cytoskeleton or associated motor proteins.

Our direct manipulation of Rab25 vesicle positioning demonstrates that upon reaching the cell periphery, Rab25 can initiate the formation of new F-actin based protrusions in a formin-dependent, Arp2/3-independent, manner. Interestingly the movement of Rab25 vesicles shows a modest but appreciable dependence on formins (fig. S4B), suggesting that the F-actin tracks involved in Rab25 vesicle positioning are also formin-dependent in this system. Rab25 associates with FMNL1 to a greater extent when able to stably interact with vesicles (Fig. 4BC), and Rab25 vesicles simultaneously coordinate the delivery of FMNL1 to the plasma membrane with the arrival of active Rho GTPases (Fig. 6C-E). On 2D substrates, whilst Rab25 wt re-positioning can drive modest changes in cell protrusions, the changes associated with repositioning of Rab25 dC are more striking. In 2D migration on a rigid flat surface, Arp2/3-driven lamellipodia are particularly important for driving cell motility, but in 3D matrices more plasticity in migration mode is observed and structures such a filopodia can promote significant invasiveness (Caswell and Zech, 2018). Interestingly, our previous work has shown that the Rab11 family promote invasiveness of cancer cells in 3D matrix, whilst effects on migration in 2D are more subtle (Caswell *et al*., 2007, 2008). The effectiveness of Rab25 dC in promoting new protrusions on 2D substrates may be explained by faster re-localisation kinetics (Fig. 2B, C), and its modest association with FMNL1 is likely enhanced upon repositioning to the cell periphery where a pool of FMNL1 resides and co-localises with Rab25 dC (fig. S9H). Strikingly, re-localisation of Rab25 to the cell periphery induced robust formation of F-actin protrusions made up predominantly of filopodia in 3D matrices, but uncoupling of Rab25 from endosomes abrogated this effect (fig. S6A, S8AB). We also found that integrin β1 is able to associate with vesicular Rab25, but is not detectable when Rab25 is truncated to inhibit vesicle recruitment (fig. S9I). Invasion and protrusion elongation driven by Rab25 is dependent upon integrin β1 ((Caswell *et al*., 2007), (fig. S8FG), and using this new magnetogenetic approach we now demonstrate that the ability of Rab25 vesicles to drive the formation of new F-actin rich protrusions in 3D matrices is a product of the ability to coordinate the activity of Rho GTPases together with delivery of FMNL1 and integrin β1.

Rab11 has previously been shown to recruit F-actin regulators which can generate tracks to permit vesicle trafficking over long distances within oocyte (Schuh, 2011). Similarly, trafficking regulator Rab27 has been demonstrated to cooperate with actin nucleators to drive long-range actin-dependent transport of melanosomes in melanocytes (Alzahofi *et al*., 2020). Our data confirm that Rab25 may similarly promote the formation of F-actin tracks (fig. S1H-I), which could be mediated through recruitment of FMNL1. The fact that Rab25 vesicles are closely associated to the F-actin cytoskeleton, is also reflected in the adaptation of cells to the introduction of magnetic force, which is only seen in a pliable 3D matrix and the adaptation observed is dependent upon the contractile F-actin cytoskeleton (Fig. 3B, fig. S7B). This observation could also indicate that endosomal vesicles are involved in mechanosensing and cellular mechanoprotection (Phuyal *et al*., 2023) and is consistent with studies describing roles for Rab11, Rab25, and the F-actin network in the positive regulation and propagation of actomyosin contractile forces in response to developmental morphological change (Dehapiot *et al*., 2020; Ossipova *et al*., 2014; Willoughby *et al*., 2021).

Interestingly, we found that a truncated form of Rab25, in which the C-terminal prenylation sites are deleted, is able to facilitate protrusion despite the lack of stable vesicle interaction (Fig. 2DE; Movie S3). Whilst it is possible that truncated Rab25 can interact with effectors and thus associate with vesicles to some extent, the limited association of Rab25 dC with FMNL1 (Fig. 4BC) and lack of detectable association with integrin β1 (fig. S9C) suggests that this is minimal. Instead, we suggest that Rab25 dC encounters FMNL1 at the plasma membrane to support the initiation of new protrusions, perhaps because Rab25 dC is still able to support the activation of Rho GTPases that are required for FMNL1 activity (Fig. 6E, Movie S10). This leads us to suggest that Rab25 plays a dual role in the regulation of F-actin by modulating Rho GTPases and associating with actin polymerising proteins. However, while these activities alone are sufficient to initiate cell protrusion, the formation of more stable protrusions requires the vesicular association of Rab25 to link to other cargoes, including integrins (Fig. 6F).

## Methods

Detailed list of antibodies, chemicals, sequences, plasmids and resources used are described in supplementary excel file S1 or accessible from https://doi.org/10.6084/m9.figshare.27084217.

### Constructs

A number of plasmid DNA constructs were used and generated in this study and these are listed with details in excel file S1. Plasmid maps are provided here https://doi.org/10.6084/m9.figshare.27084205. The online tool Benchling was used to design guideRNA sequences for specific CRISPR/Cas9 cleavage (excel file S1) and primers for cloning using Gibson assembly (www.benchling.com; provided upon request).

Plasmid constructs and donors for ssDNA synthesis (pJET-24_Rab25 DExCon, pJET-78-FMNL1-HR-SFFV-BlaR, pJET-77-FMNL1-HR-SFFV-PuroR) were prepared as described previously using Gibson assembly or restriction endonuclease based cloning ((Gemperle *et al*., 2022); see final plasmid maps). New sequences (gene specific exon/intron sequences for knock-in or cDNA for overexpression) were synthetized commercially by GENEWIZ (FragmentGENE) or IDT (gBlocks) with 25-50 bp long homologous overhangs for Gibson assembly depending on the synthesis limitations (GC-rich/repetitive sequences; sequences provided in excel file S1) and cloned into pJET1.2 (CloneJET PCR Cloning Kit. #K1231; Thermofisher) or different vector according to the requirements of the experiment (pCDH, pEGFP, pSBtet…). Universal linkers (including SpeI/SalI, XbaI/XhoI, AgeI, NheI, EcorI or other cleavage sites) were also included and used to add, switch or remove additional sequences (see final plasmid maps). Lentiviral pCDH-NB^GFP^-mCherry-Rab25 was cloned using pCDH-NB^GFP^-mCherry-Rab11a (as NB^GFP^-mCherry template; (Gemperle *et al*., 2022)) and XbaI/XhoI cleaved pCDH-tagBFP-T2A-mycBirA-Rab25 (Wilson *et al*., 2023). To create Rab25 dC mutant, Rab25 wt was cut with BspEI (=Kpn2I) and SalI to remove the C-terminus, filled with PfuX7 and religated (pCDH-tagBFP-T2A-mycBirA-Rab25 dC) or generated with specific Gibson assembly primers (pCDH-tagBFP-T2A-mycBirA-Rab25 dC), for Rab25 dN (T26N) including with the mutagenesis site. pLenti Lifeact-iRFP670 BlastR was purchased from Addgene ((Padilla-Rodriguez *et al*., 2018); #179888). Optimized Sleeping beauty transposon-based system pSBtet-puroR-BFP (Addgene Plasmid #60496; (Kowarz *et al*., 2015)) was open by SfiI cleavage and pB72_Kif5b-GFP-PDZ, a gift from Lukas Kapitein lab (van Bergeijk *et al*., 2015), used as a template to generate pSBtet Kif5b-GFP-pMagFast1_(3x)_ (last module synthesised and tandemly added via SpeI, NotI, AscI and KpnI restriction sites for possible optogenetic manipulation (Kawano *et al*., 2015)). Localization-based probe for active RhoA (tandem RBD, 4 in total separated by linker) together with iRFP670 (3 repeats to increase sensitivity) was synthetized as proposed (Mahlandt *et al*., 2021) and inserted into pCDH vector via XbaI and SalI restriction sites to create pCDH_iRFP670_3x_-RBD_4x_. pLVX-GFP-FMNL1β, a gift from Blystone lab (E. W. Miller and Blystone, 2019). All constructs were verified first by restriction analysis and then by sequencing. Further details on the cloning strategy, and plasmids used in this study upon reasonable request.

### Preparation of GFP-MNPs

The construct used to purify modified GFP (pET21_H6-mEGFP-Mms6[112-132]) prior to functionalising nanoparticles to prepare GFP-MNPs has been previously described (Kappen *et al*., 2024). The construct comprises a 6xHis-tag fused with the N-terminus of mEGFP (monomeric enhanced GFP), connected by a seven-amino acid linker with the iron-binding domain of Mms6 (22 C-terminal amino acids MKSRDIESAQSDEEVELRDALA). The pET21_H6-mEGFP-Mms6[112-132] construct was transformed into E. coli BL21-CodonPlus-RIL (Agilent) for preparative overexpression of H6-mEGFP-Mms6[112-132]. Cells were grown at 37 °C in LB medium supplemented with 100 μg/ml Ampicillin and 32 μg/ml Chloramphenicol. Subsequently, the expression of H6-mEGFP-Mms6 Mms6[112-132] was induced with 0.5 mM IPTG (isopropyl β-D-1-thiogalactopyranoside, Thermo Scientific) at OD600 = 0.6, followed by culturing at 16 °C overnight. The harvested cells (>4000 g for 10 minutes) were resuspended in lysis buffer (50 mM HEPES, 150 mM NaCl, 8 M urea, pH 7.5-8.0) followed by tip sonication (4x 3 min; 50% duty cycle, Sonifier 250, Branson) and centrifugation (20 000 g, 30 min) to pellet insoluble material. The protein was first purified using immobilized metal affinity chromatography (IMAC) by loading supernatant on a 5 ml hiTrap chelating HP column loaded with nickel (II) chloride and equilibrated in the 20 mM HEPES, 150 mM NaCl, pH 8.0 (HBS). The protein was eluted with HBS, 500 mM imidazole, pH of 8.0 using a linear gradient that spanned 10 times the bed volume. Subsequent purification based on size exclusion chromatography was performed, if needed (aggregates observed upon 1h incubation with 8 M urea which leads to a significant increase in total protein yield), using an FPLC system (Äkta Explorer, GE Healthcare) by loading protein solution onto a SEC column (HiLoad 16/60 Superdex 75, prep grade or HiLoad 26/60 Superdex 75, prep grade) that had been equilibrated in HBS. Protein integrity purity and integrity were confirmed by 12 % SDS-PAGE (expected size 31.2 kDa).

To prepare functional GFP coated MNPs (=GFP-MNPs), characterized and described previously (Kappen *et al*., 2024), purified H6-mEGFP-Mms6[112-132] (HBS buffer) was first transferred into water pH 7.0 using a buffer exchange column (PD10) and resulted concentration increased (> 100 uM; molar extinction coefficient at 488 nm = 56 000), if bellow, by ultrafiltration (Amicon Ultra-4, 10 kDa cut-off, 20 mM HEPES pH 7.5, room temperature (RT)). Magnetic core particles with average diameters of 12 nm (iron concentration of 900 μM, pH 2), synthesized as described before (Kappen *et al*., 2024), were diluted to final 90 µM concentration (water, pH 2) and sonicated (Ultrasound bath, 10 min, 10-15 °C). Immediately after these nanoparticles were added to a >75-fold molar excess of purified H6-mEGFP-Mms6[112-132] (> 100 uM; Water pH 7.0) followed by additional sonication (Ultrasound bath, 5 min, 15-20 °C) and incubated for > 1 h (RT). The protein excess was removed by ultrafiltration (Amicon Ultra-4, 100 kDa cutoff, 20 mM HEPES pH 7.5, RT) until the filtrate was devoid of unbound coating protein. Finally, GFP-MNPs were incubated in 1h paraformaldehyde (4% PFA in 20 mM HEPES, RT) followed by extensive washing (>3x) using ultrafiltration (Amicon Ultra-4, 100 kDa cutoff, 20 mM HEPES pH 7.5, RT). The final GFP-MNPs (20 mM HEPES pH 7.5, 10 µg/ml ciprofloxacin to prevent bacterial growth) were stored for up to 14 days at RT prior to live cell experiments, and no significant deterioration in magnetic attraction was observed. Directly before use inside living cells, GFP-MNPs were centrifuged to remove aggregates (2500-4000 g, 5 min, RT).

### Cell culture

Ovarian cancer cell line A2780-DNA3 (Caswell *et al*., 2007) was cultured in RPMI-1640 medium (R8758, Sigma). TIFs (Telomerase immortalised fibroblasts (Caswell *et al*., 2007)) and HEK293T cells were cultured in Dulbecco’s Modified Eagles Medium (DMEM; D5796, Sigma). All cell culture media were supplemented with 10% v/v foetal bovine serum (FBS) and ciprofloxacin (0.01 mg/ml; Sigma), and cells were maintained at 37°C in a humidified atmosphere with 5% (v/v) CO_2_. The cells were confirmed to be free of mycoplasma contamination by PCR analysis. A2780 were selected for antibiotic resistance as follows: puromycin 0.5-1 µg/ml; blasticidin 5 µg/ml.

### CRISPR/Cas9 RNP based transfection and homologous recombination

CRISPR/Cas9-based modifications (FMNL1), including the DexCon approach (Rab25), were performed as previously described in detail (Gemperle *et al*., 2022). Single-stranded DNA (ssDNA) coding sequences, flanked by specific homologous sequences (FMNL1 exon 2 and Rab25 exon 1), were employed to generate a specific CRISPR-based knock-in outcome, eliminating the necessity for clonal selection (Gemperle *et al*., 2022; Li *et al*., 2017). Purified ssDNA were prepared as previously described (Bennett *et al*., 2021; Gemperle *et al*., 2022) using dsDNA (pJET-24_Rab25 DExCon, pJET-78-FMNL1-HR-SFFV-BlaR and pJET-77-FMNL1-HR-SFFV-PuroR) and PCR with reverse and biotinylated forward primers (primer sequences in excel file S1). Biotinylated PCR-product was purified using Dynabeads™ MyOne™ Streptavidin C1 (Cat# 65001, Thermo), anti-sense ssDNA strand eluted by 20mM NaOH (later neutralized by HCl) and final ssDNA purified using SPRI beads (AMPure XP, Agencourt). Anti-sense ssDNA strand was verified by agarose-gel electrophoresis (denaturated) and/or sequencing and used for knock-in (detail protocol provided in https://doi.org/10.48420/16878859 (Gemperle *et al*., 2022)). High fidelity Cas9 (Alt-R S.p. HiFi Cas9 Nuclease V3, IDT) and guideRNA (crRNA:tracrRNA, synthetized by IDT) were delivered together as RNP (final 10 nM) with ssDNA donor (300-530b homologous arms; 15-50 ng) via CrisperMAX (Thermo) combined with nanoparticles for magnetofection (for sequences see excel file S1). The Combimag nanoparticles (OZBiosciences) were used in accordance with the manufacturer’s instructions, utilising a magnetic plate.

A2780 stably expressing tagBFP-T2A-TetOn3G (sorted for tagBFP) were used to integrate TRE3GS-NB^GFP^-mCherry into Rab25 exon1 locus, FACS sorted and specific knock-in validated by fluorescent microscopy, western blot and by sequencing of PCR-amplified genomic DNA as previously described (primers are listed in excel file S1; (Gemperle *et al*., 2022)).

To inactivate the human FMNL1 gene in A2780 cells, we employed our previously published strategy for gene inactivation, which obviates the necessity for clonal selection through the precise integration of the puromycin (PuroR) and/or blasticidine (BlaR) resistance cassette (exon-SFFV-PuroR/BlaR-polyA-exon) (fig. S11A; (Gemperle *et al*., 2022)). While we were able to readily prepare FMNL-/+ heterozygotes, the use of any BlaR/PuroR combination proved unsuccessful in obtaining FMNL1-/-homozygotes (fig. S11B). This finding suggests that FMNL1 is essential for A2780 survival, as has been demonstrated in mice (M. R. Miller *et al*., 2017), or that the efficiency of the homozygous knock-in is too low. Therefore, we generated clones of A2780 (33 in total) with one allele targeted by ssDNA carrying puroR followed by puromycin treatment in the hope that the second allele would be inactivated by the more common indel-rich out-of-frame non-homologous end-joining pathway (fig. S11C). Surprisingly, all puromycin-surviving clones were once again heterozygous, with the FMNL1 protein oscillating around the expected size on the Western blot, indicating the presence of in-frame indels. However, these indels were always in frame, suggesting that FMNL1 plays an essential role in A2780 (fig. S11D).

### DNA transfection and siRNA-mediated protein depletion

Unless otherwise stated, Lipofectamine 2000 (Invitrogen) was employed to transiently transfect plasmid DNA into A2780 cells in accordance with the manufacturer’s instructions. A2780 transfected with pSB-pSBtet-Kif5b-GFP-pMagFast1were-codelivered with Sleeping Beauty transposase SB100X in the ratio as suggested (Kowarz *et al*., 2015) followed by puromycin selection (1 µg/ml) and FACS-sorting (BFP/GFP; 500 ng/ml dox treated, 48h).

All FMNL1 targeting siRNAs were nucleofected into A2780 cells or A2780 FMNL1-/+ (PuroR/+) harvested from sub-confluent 150 mm cell culture dish using a nucleofector (Amaxa, Lonza) with solution T (VCA-1002, Lonza), program A-23, 5μl 20 µM siRNA (= total 100 pmol; 1 µM) as per the manufacturer’s instructions. Silencer select siRNA reagents, with locked nucleic acid chemical modifications to increase specificity and stability, were purchased from Invitrogen: negative control (NS) (4390843); FMNL1 siRNA #s26 (4392420, ID: s2226) and #s27 (4392420, ID: s2227) and #s28 (4392420, ID: s2228). FMNL1 siRNA #s26, #s27 and #s28 were combined together to create a smart pool.

### Lentivirus packaging generation

Lentiviral constructs (pCDH, pLenti, pLVX) coding different FMNL1 and Rab25 fusion constructs (see excel file S1) were used to generate stable cell lines as follows: Lentiviral particles were produced in HEK293T cells via a polyethylenimine (PEI)-mediated transfection with packaging plasmids pM2G and psPAX2 with lentiviral vector (pCDH, pLenti, pLVX) in a ratio of 1.5:1:2. Supernatants were collected at 94 h post-transfection, filtered through a 0.45 μm filter, and added to target cells. Transduced cells were selected by appropriate antibiotics (GFP-FMNL1 β; puromycin) or sorted by FACS (Aria II, BD; iRFP670-Lifeact, NB^GFP^-mCherry fusions, tagBFP for T2A-mycBirA* fusion protein constructs and for tagBFP-T2A-TetOn3G prior Rab25 DExCon modification). NB^GFP^-mCherry fused –Ctrl or Rab25 dC or Rab25 wt were sorted for matched expression levels.

### Cell-Derived Matrix preparation and cell migration analysis

Cell derived matrices were generated as previously described (Caswell et al., 2007; Cukierman et al., 2001) on either 40 mm coverslips (#1.5H; for magnetic live imaging experiments) or tissue-culture plastic 12-well plates (for long-term time-lapse microscopy). Briefly, plates were coated with 0.2% gelatin (v/v, Sigma Aldrich), crosslinked with 1% glutaraldehyde (v/v, Sigma Aldrich) and quenched with 1M glycine (Thermo Fisher) in PBS before washing and equilibration in DMEM. TIFs were plated onto the prepared plates at a density that would ensure confluency within the following day or two. TIFs were cultivated for a period of 8–10 days, with the medium being replaced with DMEM supplemented with 50 μg/ml ascorbic acid (Sigma-Aldrich) 24 h after seeding and subsequently every 48 h. Cells were denuded with extraction buffer (0.5% (v/v) Triton X-100; 20 mM ammonium hydroxide (NH4OH)) to leave only matrix) and washed twice with PBS+ (Dulbecco’s phosphate buffered saline with CaCl_2_ and MgCl_2_, Sigma-Aldrich). Finally, the matrices were incubated with 10 μg/ml DNase I (Roche) and washed three times with PBS+, before A2780 cells were plated at a low density into 12-well plate or 40 mm coverslip inserted into 60 mm dish. The cells were then allowed to spread and start migrating overnight (40 mm coverslip; see spinning disc confocal) or at least 4 h before being imaged (12-well plate). The images were obtained using the confocal spinning disc microscopes described below (magnetic experiments, see confocal live cell imaging section) or on an DMi8 inverted widefield microscope (Leica) using either a 20×/0.40 L Plan Fluor dry-corrected or a 10×/0.25 N Plan Fluor objective, and LED light source for transmitted light. Images were collected using a DFC 9000 – monochromatic sCMOS camera, and cells were maintained at 37°C and 5% CO_2_ for the duration of imaging. LAS X software (Leica) was used to acquire images of multiple positions per well, every 5 min for 16 h. For cell migration analysis, at least 70 cells (in total) were individually manually tracked per condition from three independent replicates using CellTracker software (Piccinini *et al*., 2015), and this was used to calculate the average cell speed and persistence, where persistence is equal to the path length divided by Euclidean distance.

### Magnetic tweezers

Magnetic tweezers were home-built by combining sharp paramagnetic tip and neodymium magnets as follows: To create a sharp paramagnetic tip, an iron wire (0.4 mm diameter) was pulled in the flame of a Bunsen burner. The string was pulled at a slow rate, resulting in the formation of two sharp extremities with a diameter of 20 µm. These were used as paramagnetic tips on a permanent magnet in two distinct configurations, but generated the same steady-state gradient profiles of GFP-MNPs at a distance of 0-150 µm (measured in Fig. S2): Sharp tip (of length 1,5 mm) was placed on top of a tower of small resizing magnets made of neodymium iron boron N-45 (dimensions diameter×height: 1×2, 3×6, 5×10, 8×30, 10×20; Supermagnete, Gottmadingen, Germany) or similarly as described (Kappen *et al*., 2024; Liße *et al*., 2017) by attaching a paramagnetic tip (of length 3-5 mm) on top of block neodymium magnet (5×1,5×1 mm, N-45, Supermagnete) and glueing it into a plastic pipette tip, ensuring that the tip extend 0.5 mm beyond the end of the magnet. Subsequently, magnetic tweezers were attached to the micromanipulation apparatus (mikromanipulator 5171 or InjectMan NI2), which was situated in close proximity to the Zeiss microscope with a CSU-X1 spinning disc confocal (Yokagowa) or mounted on a Leica microscope equipped with a Dragonfly 503 spinning disc confocal (Andor), respectively. Both the apparatus and the microscope were surrounded by heating chambers.

### Microinjection

Prior to microinjection, cells were seeded on 40 mm coverslips (#1.5H) with FN coating or with prepared CDM and inserted into a non-magnetic FCS2 chamber (Bioptechs, United States) or into newly designed custom-made lidded imaging chamber insert (Vivanto, Czechia; available for purchase). Subsequently, these were inserted into Zeiss microscope with a CSU-X1 spinning disc confocal (Yokagowa) or Leica microscope equipped with a Dragonfly 503 spinning disc confocal (Andor Technologies), respectively. GFP-MNPs (0,8-2μM solution in 20 mM Hepes) were injected into cells using commercial injection capillaries (Femtotip, 0.5 μm inner and 1.0 μm outer diameter; Eppendorf) and a micro manipulation system (mikromanipulator 5171 or InjectMan NI2 and 10 FemtoJet Express, Eppendorf) in semi-automatic mode. The capillary pressure was maintained at a constant level of 18-20 psi, while the injection pressure (80 – 180 psi) and time (0,3 – 2 s) were subject to inverse variation. To microinject cells migrating in CDM, higher injection pressure was usually needed (180 psi). Following microinjection, the cells were allowed to recover (5-10 minutes) before being imaged. The microinjected cells were then subjected to a quality control assessment to ascertain their unharmed capacity for migration and to check the quantity and quality of the microinjected GFP-MNPs. Subsequently, injection capillary was replaced with magnetic tweezers followed by live cell imaging.

### Spinning disc confocal cell imaging and analysis

Magnetic spatiotemporal control of GFP-MNPs in living cells was performed with A2780 cells (± expressing NB^GFP^-mCherry fusion protein variant) situated on FN-coated or CDM-covered 40 mm coverslips (#1.5H), inserted into the aforementioned chamber insert (see microinjection), and the aforementioned micromanipulator with magnetic tweezers (see magnetic tweezers) at a distance of 30-120 μm from cells. GFP-MNPs were delivered by microinjection (see microinjection). Imaging was recorded using CSU-X1 spinning disc confocal (Yokagowa) on a Zeiss Axio-Observer Z1 microscope or Dragonfly (Andor Technologies) spinning disc confocal Leica microscope with an APO 40×/1.25 NA oil-immersion or APO 63x/1.20 water-immersion objectives. Microscopes were equipped with an Evolve EMCCD camera (Photometrics) or a Zyla 4.2 PLUS sCMOS camera, respectively, both with motorized XYZ stage and 405, 488, 561, 637 lasers. Fluorescent Iimages were captured with the appropriate excitation/emission spectrum and white LED source was used for transmitted light to monitor the position of the magnetic tip. Images were captured using SlideBook 6.0 (3i) or Fusion software, respectively. The objective drift was corrected automatically using the HWada coordinate shift plugin in Fiji-ImageJ (NIH, USA) wherever necessary. During the imaging and manipulation processes, the cells were maintained in a 1x Opti-Klear medium (6 ml; Marker Gene Technologies Inc.) supplemented with 10% (v/v) FCS in a heating chamber. The time interval acquisition was always set according to the specific requirements of the experiment, with a duration of either 2 seconds for rapid kinetics and magnetic tip positioning or 30 seconds for long-term imaging. The laser powers were set to an optimal level that would not induce stress on the cells. Laser-based (point visiting) Definite Focus was employed to maintain the correct plane in focus, with manual re-adjustment if re-focus occurred due to magnetic tip movement. On each day of the experiment, the functionality of the magnetic tweezers and GFP-MNPs was initially evaluated on control A2780 cells (which express NBGFP-mCherry). A Z stack was acquired before and after 10–60 minutes of manipulation and used for the analysis of protrusion growth (LifeAct reporter), as described in (Fig. 1A-D), using Fiji-ImageJ (NIH, USA). Only cells with visible GFP-MNPs relocalization/enrichment were included in the analysis, and protrusions were induced by Rab25 before various inhibitor treatments to observe their effects. Magnetic attraction and release kinetics of NB^GFP^-mCherry-(X) and GFP-MNPs in living cells quantified as described in Fig. 2B-C legend. Note: NB^GFP^-mCherry-Rab25 wt overexpressing cells exhibiting a higher proportion of diffusively localized Rab25 with fast kinetics (approximately 25%; not observed with A2780 DExCon-modified NB^GFP^-mCherry-Rab25 cells) were not included in this analysis. The relative stress maps were analysed using brightfield images of cells migrating in CDM, taken before and after the magnetic attraction of GFP-MNPs via particle image velocimetry (PIV) analysis plugin (Tseng *et al*., 2012) in Fiji-ImageJ. Kymographs were created from time-lapse images using KymoToolBox in Fiji-ImageJ. The AI-based Kymobutler tool (Jakobs *et al*., 2019) was used to analyse particle traces in kymographs via the cloud application or as a plugin in Fiji-ImageJ.

Additional fluorescent imaging was conducted using the aforementioned spinning disc confocal microscopes with 63x/1.40 Plan-Apochromat objective on randomly chosen representative fixed (see Immunostaining) or live cells situated on FN-coated µ-Plate 96 plate (#1.5 IbiTreat) or FN-coated 96 well plate (cellvis, #1.5H cover glass) and analysed using Fiji-ImageJ (NIH, USA), as specified in the figure legends. For colocalization analysis, Colocalization finder plugin in Fiji-ImageJ was used to generate Pearson’s correlation coefficient (no thresholding, noise subtraction using the ScatterPlot function).

### Magnetic force calculation

Quantification of forces exerted by magnetic manipulation were quantified from images taken by spinning disc confocal microscopes (see spinning disc confocal). The rosette plot was generated in Prism 8.0 (GraphPad Prism software); the angles between cells with a GFP-MNPs gradient and a magnet-aligned line extending from the magnet tip were divided into twelve 30° segments; the relative sizes of the triangular segments are directly proportional to the number of cells with angles contained in each bin. The attraction and relaxation kinetics and the applied force on GFP-MNPs were determined in the cell cytoplasm at the steady state of attraction. The distance-dependent decrease in particle intensity I(r) was fitted exponentially. The theoretical profile was assumed to follow the Boltzmann law:

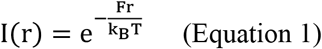

The applied force F was determined from the fitted exponential decay (k_B_: Boltzmann constant; T: temperature).

### Immunostaining

For all antibody staining procedures, cells were fixed in 4 %(w/v) paraformaldehyde (PFA, Sigma-Aldrich) for 10-15 min RT (or 2% PFA 10 min on ice to preserve endogenous fluorescent signal, e.g. mCherry) followed by mild permeabilization with 0.1% Triton-X100 (Sigma-Aldrich, 2-10 min) in PBS (Dulbecco’s phosphate-buffered saline) or 0.1 % Saponin (Sigma-Aldrich) added to all antibody incubation steps (to better preserve endosomes). Fixed cells were blocked in 3% (w/v) bovine serum albumin (BSA) (Sigma-Aldrich) in PBS and incubated with corresponding primary and secondary antibodies diluted in 3% BSA in PBS for 1-2 h RT. Cells were washed 3-times with PBS between each incubation step and once with ddH_2_O before mounting with either ProLong Diamond Antifade Mountant (Molecular Probes, Invitrogen) or ProLong Gold Antifade Mountant (Molecular Probes, Invitrogen).

### Antibodies and reagents

The following antibodies and reagents were used (details in excel file S1): α-tubulin (mouse mAb DM1A, #ab7291), Rab25 (rabbit mAb DH46P, #13048S), RFP (rat mAb 5F8), GFP (mouse mAB 1E10H7; #66002-1-Ig), FMNL1 (rabbit pAB, #27834-1-AP), GAPDH (rabbit pAb, #G9545), integrin β1 (mouse mAB 18/CD29, #610467; immunoblot), integrin β1 (mouse mAb, 5PD2; blocking), integrin β1 (rat mAb, AIIB2; blocking). Streptavidin DyLight®-800 (1:5000) (Thermo Fisher Scientific). Fluorescent secondary antibodies (Jackson ImmunoResearch Laboratories or Li-cor, Invitrogen) were used as recommended by the manufacturer. The reagents used were puromycin dihydrochloride (Thermo) and blasticidin S HCl (Gibco), fibronectin (sigma), Dynabeads™ MyOne™ Streptavidin C1 (Thermo), Agencourt AMPure XP (Beckman Coulter). MagReSyn Streptavidin microspheres (ReSyn Biosciences). Doxycycline hydrochloride (Sigma) was dissolved as 250 μg/ml in water (stored -20 °C in aliquiots) and used within 14 days if kept in 4 °C.

### Knock-sideways imaging and analysis

A2780 cells were nucleofected (VCA-1002, Lonza) with 2 μg pMito-iRFP670-FRB and 1 μg of the required GFP–FKBP (Ctrl or -Rab25) using programme A-23 (Lonza). The following day, rapamycin was added to the cell growth medium at a final concentration of 200 nM for 4 h. Cells were then fixed and stained (see Immunostaining) for the FMNL1 (rabbit pAB, #27834-1-AP; 1:400) followed by Cy3 secondary antibody detection (Molecular Devices). Z-stacks were captured using CSU-X1 spinning disc (Yokogawa) on a Zeiss Axio-Observer Z1 microscope with a 63×/1.46 α Plan-Apochromat objective, a Prime 95B Scientific CMOS camera (Photometrics) and Slidebook software (3i). All images were processed using ImageJ, relocalisation of GFP–FKBP or GFP–FKBP–Rab25 proteins to mitochondria was analysed using the Colocalization finder plugin and scored based on Pearson’s correlation coefficient (no thresholding, noise subtraction using the ScatterPlot function). For FMNL1 redistribution, a custom-made script was used to create mitochondrial mask (available at https://doi.org/10.48420/19391597) and the mean pixel intensity of the FMNL1 protein was compared between that in mitochondria and that in the cell body.

### Proliferation

Cell proliferation rate was analysed via live, real-time, label-free, image-based measurements using xCELLigence RTCA eSight system (Agilent) situated in temperature and CO2 controlled incubator. A total of 5,000 cells were seeded into each well of the E-Plate VIEW 96 (quadruplicates per condition) and automatically imaged at 2 h intervals over a 70-h period using a 10x objective. Subsequently, the RTCA eSight software was trained to create object masks in order to determine cell numbers in the field of view.

### Scanning transmission electron microscopy (STEM) tomography

GFP-MNPs (1,1 uM, Opti-MEM) were electroporated (0,2 cm aluminum cuvettes (100 µL), Lonza) into A2780 DExCon-modified NB^GFP^-mCherry-Rab25 cells (pre-treated with dox >94 h; 250 ng/ml) by using an Amaxa Nucleofector II Electroporation machine (Lonza) on program A-023. This approach results in a high efficiency of GFP-MNPs delivery (Kappen *et al*., 2024) and the ratio of GFP-MNPs:Rab25 was close to 1:1 or less, ensuring that the majority of GFP-MNPs were Rab25-bound. Cells were then let to spread (3 h) on FN-coated 12 mm round coverslip before rinsing with a Sörensen’s phosphate buffer (SB, 0.1 M, pH 7.2-7.4) prewarmed to 37 °C and fixing in a mixture of 1% glutaraldehyde and 2.5% paraformaldehyde in SB for 30 min RT and then overnight at 4 °C. Next day, fixatives were washed out with SB and cells were post-fixed with 1% OsO_4_ solution in SB for 1 h in the dark. Samples were washed again with SB and MilliQ water and dehydrated with a series of water/acetone mixtures finished with dried acetone. Finally, embedding was performed into Epon-Durcupan resin followed by polymerisation at 60 °C for 3 days. Embedded samples were cut into semithin sections of 500nm thickness, placed on gilded copper slot (2 × 0.5 mm) without any supporting film, so that the section exceeds the whole slot and is attached to its rim. To enhance the contrast, the section on a slot was immersed into a droplet of a 2% uranyl acetate/water solution for 30 min. However, such sample is still too fragile for the scanning transmission electron microscopy (STEM) tomography. To make it more durable, it was coated with 4nm carbon layer from both sides (Leica EM ACE600 Sputter coater). Finally, fiducial markers (6nm gold nanoparticles) were introduced on both sides of the section to facilitate the tilt series reconstruction. Cells positive for GFP-MNPs and high-contrast recycling endosomes [50-200 nm], were firstly found by standard TEM imaging in the Jeol JEM-1400 Flash transmission electron microscope equipped with tungsten cathode, bottom-mounted FLASH 2kx2k CMOS camera and operated at 80 kV. Sample was mapped with Limitless Panorama (LLP) software at high magnifications (up to x30 000) and the stitched map was used to re-find the regions of interest in a high-end TEM Jeol JEM-F200. Following the acquired LLP map, the regions of interest were located in the Jeol JEM-F200 - high-end S/TEM electron microscope equipped with cold field emission gun, TVIPS TemCam – XF416 cooled 4kx4k CMOS camera and operated at 200 kV. The series of tilted images (tilt series, TS) of individual regions within the section was acquired in the scanning mode (STEM), in which was employed dark field STEM detector. The TS acquisition was controlled by a SerialEM software (program developed by Dr. David Mastronarde, University of Colorado) including STEM tomography module. The TS was collected automatically in one direction from -60 to +60 degrees with a step of 1°, recorded image time frame was 40 s and used magnification was x100 000. Reconstruction of the obtained TS was performed in a module of IMOD software called “Etomo”, where the fiducial markers were used for fine alignment of TS frames. TS frames were binned to final size of 1024×1024 pixels, the reconstruction itself was performed with a SIRT-like filter of 15 and the LUT of finished tomogram was inverted to look like a bright field image – this sequence generated the best image quality to distinguish the studied objects for a consequent segmentation. GFP-MNPs and endosomes were manually segmented using MATLAB-based Microscopy Image Browser software (Belevich *et al*., 2016) and visualized using Imaris (Bitplane).

### Proximity labelling

Cells stably expressing mycBirA* fusion protein constructs (see Lentivirus packaging generation) were plated onto tissue culture plates (1×100 mm for FMNL1 detection or 4×150 mm for integrin β1, respectively) at a density to ensure sub-confluency the following day and BioID performed as described previously (Wilson *et al*., 2023). Briefly, more than 4 h after cells were plated, 1 μM biotin (Sigma-Aldrich) was added to the cell culture medium and cells were incubated for 16 h. Subsequently, cells were washed in PBS before addition of the BioID lysis buffer [50 mM Tris pH 7.4, 500 mM NaCl, 5 mM EDTA, 0.4% SDS, 1 mM dithiothreitol (DTT), supplemented with the protease inhibitors: 100 μg/ml aprotinin (Sigma-Aldrich), 100 μg/ml leupeptin (Sigma-Aldrich), 0.5 mM AEBSF (Calbiochem), 50 μM PD150606 (Calbiochem) and 500 μM ALLN (Calbiochem); RT]. Cell lysates were collected using a cell scraper, mixed with an equal volume of 50 mM Tris pH 7.4 and Triton X-100 (final concentration 2%, Sigma-Aldrich), and then lysed with a needle. Lysates, clarified by centrifugation (16,000 *g* for 10 min at 4°C), were equalized to total protein amount and incubated with 15 μl of MagReSyn Streptavidin microspheres (per 10 cm plate; ReSyn Biosciences) overnight at 4°C with rotation. Beads were extensively washed: twice in wash buffer 1 (BioID lysis buffer with 2% SDS), once in wash buffer 2 (0.1% deoxycholate, 1% Triton X-100, 500 mM NaCl, 1 mM EDTA, 50 mM HEPES pH 7.4) and once in wash buffer 3 (0.5% NP-40, 0.5% deoxycholate, 1 mM EDTA, 10 mM Tris pH 8.1). Bound proteins were eluted by the addition of 40 μl of 2× sample buffer saturated with biotin (250 mM Tris-HCl pH 6.8, 2% SDS, 10% glycerol, 0.2% Bromophenol Blue, 20 mM DTT, with >1 mM biotin) at 70°C for 5 min. Finally, eluted samples were analysed by western blot analysis.

### SDS-PAGE and Western blotting

Cells were lysed in a denaturing lysis buffer (2% SDS, 20% glycerol, 120 mM Tris-HCl (pH 6.8), and 0.1% bromophenol blue) and heated for 10 minutes at 98 °C followed by SDS-PAGE under denaturing conditions (4–12% Bis-Tris gels; Invitrogen). Subsequently, resolved proteins were transferred to nitrocellulose membrane using the Trans-Blot Turbo Transfer System (Bio-Rad), then blocked with 4% BSA-TBS (Tris-buffered saline; 1 h), incubated overnight at 4°C with the appropriate primary antibody in 2-3% BSA TBST (2-3% BSA in TBS with 0.05% Tween 20; overnight 4°C) and fluorophore-conjugated secondary antibody (2.5% milk in TBS with 0.05% Tween 20; 1h RT). The membranes were imaged using an infrared imaging system (Odyssey; LI-COR Biosciences).

### RNA extraction & reverse transcription, PCR

RNA extraction and reverse-transcription were performed as previously described (Gemperle *et al*., 2022). PCR was conducted using PfuX7 home-made polymerase (Nørholm, 2010), in accordance with the standard Pfu polymerase protocol. Reactions were performed with 500nM primers designed for this study (listed in Excel file S1). PCR products were run on a 2% agarose gel with SYBR green to check primer specificity and for quantification of band intensities.

### Statistical analysis and bioinformatics

The data were tested for normality and analysed using one-way or two-way ANOVA with corresponding post hoc tests as indicated in legends. If the data were not normally distributed, ANOVA on ranks was used instead. All statistical analysis was done with GraphPad Prism software. P values are described in relevant figure legends, where ∗∗∗ denotes p < 0.001, ∗∗ denotes p < 0.01, and ∗ denotes p < 0.05. The number of independent experiments (*N*) and data points (*n*) is indicated in each of the figure legends.

Bioinformatic re-analysis of Rab25-BioID (proximity labelling) was done on dataset available via the PRIDE partner repository with the dataset identifier PXD033693 (Wilson *et al*., 2023). Proteins exhibiting a minimum of 1.8-fold MS1 intensity-based (mass spectrometry-based label-free quantification) increase in the Rab25-BioID sample relative to the BioID control sample (BirA alone), observed at least in 3 out of four biological replicates, were considered to be statistically significant. Selected Rab25proximity interactome was visualized by STRING protein-protein interaction analysis (https://string-db.org/).

## Supporting information

Excell file S1

movie S6_Figure 5B bottom

Movie S7_Fig. 5D top

movie S8_Figure 5D bottom

## Data and materials availability

List of primers, antibodies, chemicals, materials and all necessary sequences for replication of this study are provided (excel file S1 or accessible from https://doi.org/10.6084/m9.figshare.27084217). Annotated plasmid maps used and generated in this study can be download from https://doi.org/10.6084/m9.figshare.27084205. Plasmids can be provided upon reasonable request from corresponding author. Supplementary movies can be downloaded from https://doi.org/10.6084/m9.figshare.22155083.

## Acknowledgement and funding

We would like to thank all the members of the Caswell (Manchester, UK) and Gregor (Prague, CZ) labs for their support, Lukas Kapitein for providing pB72_Kif5b-GFP-PDZ plasmid; Scott Blystone for pLVX-GFP-FMNL1β plasmid; Davide Normanno for his constructive input on the technical aspects of microinjection; Helena Raabová, Erik Vlcek and Vlada Filamenko for their invaluable technical assistance with the challenging electron microscopy; Martin Kopecky for production of custom-made lidded imaging chamber; Peter March (WTCCMR, Manchester) and Ivan Novotný (IMG CAS, Prague) for their outstanding technical assistance in setting up an imaging system for magnetogenetic manipulation of GFP-MNPs, consisting of confocal spinning disc microscope (CSU-X1, Manchester; Dragonfly, Prague), remote-controlled micromanipulator surrounded by a heating chamber and FemtoJet pump for microinjection; WTCCMR and IMG CAS core facilities for the assistance with flow cytometry and sorting.

This project received funding from the European Union’s Horizon 2020 research and innovation programme under grant agreement No [836212], CRUK (DCRPGF\100002, and the Wellcome Trust (203128/A/16/Z and 226804/Z/22/Z). The Flow Cytometry Core Facility is supported in part by the University of Manchester with assistance from MRC Grant ref MR/L011840/1.

This work was supported by the Grant Agency of the Czech Republic (GA21-21736S); the Institutional Research Project of the Czech Academy of Sciences (RVO 68378050); National Institute for Cancer Research (Programme EXCELES, LX22NPO5102) - Funded by the European Union - Next Generation EU;. We acknowledge the Light Microscopy Core Facility, IMG, Prague, Czech Republic, supported by *MEYS CR (LM 2023050 Czech-BioImaging, project* CzBI-IMG-LMCF-155; LM2018129, CZ.02.1.01/0.0/0.0/18_046/0016045) and RVO – 68378050-KAV-NPUI, for their support with the confocal imaging presented herein.

## Author Contributions

Author contributions: Conception and design of the work, J.G., P.C., M.C., J.P.; acquisition, analysis and interpretation of data, J.G., M.K., P.C; writing – original draft: J.G., P.C; writing – review and editing: all authors. Funding, J.G., M.G., P.C. Technical and material support: D.L., J.P., E.S., C.M.

Competing interests

The authors declare that they have no competing interests.

## Supplementary Figures

**Figure S1.**
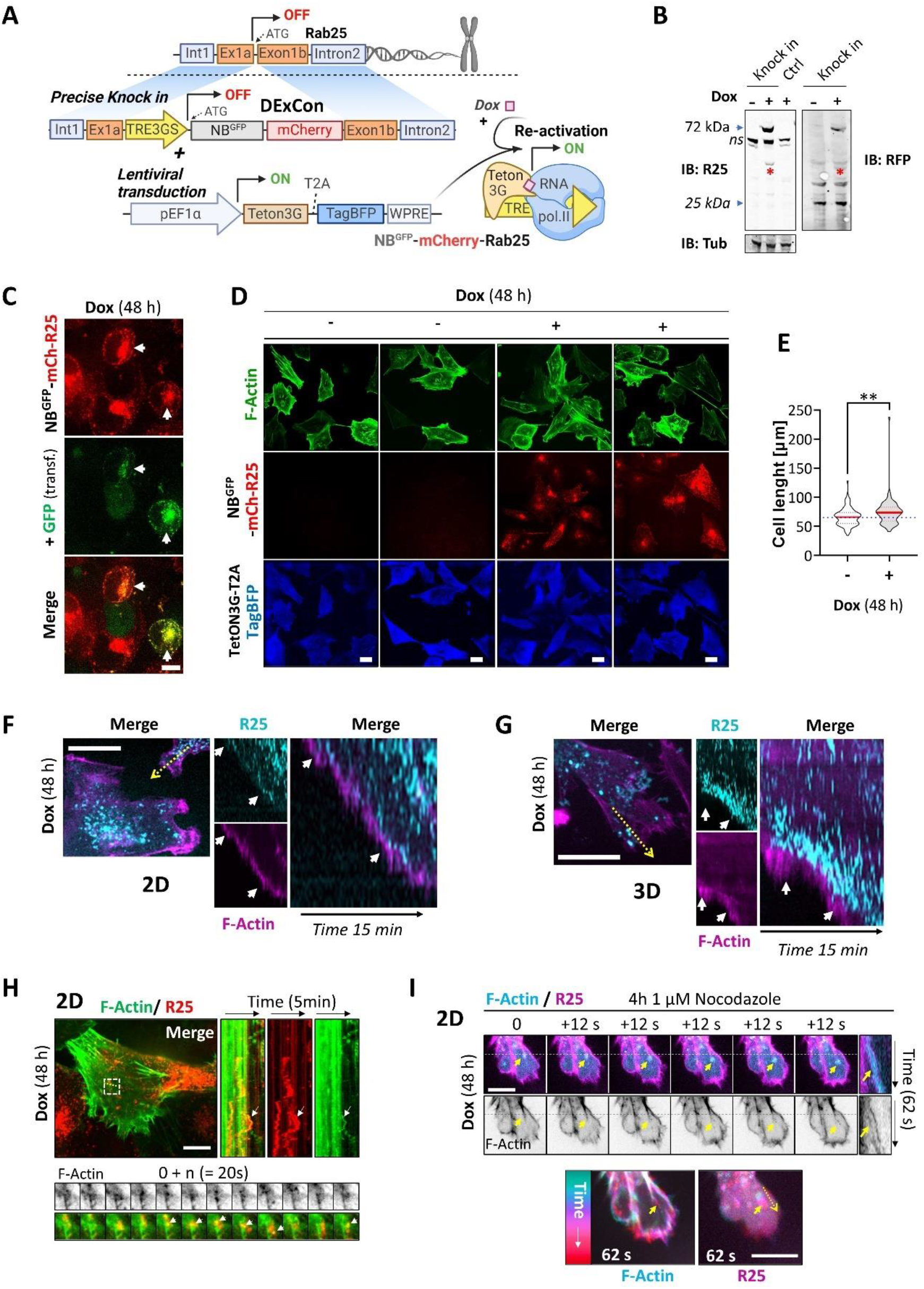
Positive correlation between the trafficking of Rab25 endosomes towards the plasma membrane (PM) and actin polymerisation. **A)** Schematic representation of DExCon approach based on CRISPR/Cas9 editing used to generate A2780 DExCon-modified NB^GFP^-mCherry-Rab25 cells. Created with BioRender.com. **B)** Immunoblots A2780 DExCon-modified NB^GFP^-mCherry-Rab25 cells (± dox 48h; 250 ng/ml) stably expressing Lifeact-iRFP670. Fluorescent antibodies: anti-Rab25 (R25), anti-mCherry (RFP) shown as black and white. Tubulin (Tub), loading control. Ctrl = un-modified wt A2780; arrow, size of predicted endogenous Rab25 (bottom) or dox-reactivated modified NB^GFP^-mCherry-Rab25 (top); ns, non-specific band; star, mCherry/iRFP670 fusion band caused by hydrolyzed C⩵N acylimine bond due to boiling (Gross et al., 2000). **C, D, F, G, H, I)** Representative confocal spinning-disk live cell images of A2780 DExCon-modified NB^GFP^-mCherry-Rab25 (R25; dox 48 h, 250 ng/ml) stably expressing Lifeact-iRFP670 (F-actin). Scale bar 20 μm. **C)** Cells also transfected with GFP. Arrows, colocalization of mCherry/GFP at perinuclear recycling compartment. **E)** Violin plot shows the quantification of cell length (shown in D) measured as maximum ferret diameter based on Lifeact-iRFP670 (F-actin). n > 74 cells ± dox from, N = 3. Two-tailed Mann–Whitney U test (**P < 0.01). **F)** Live imaging of cells on FN, dotted line shows orientation of kymograph. Arrow, positive correlation between trafficking of Rab25 endosomes towards the plasma membrane and actin polymerisation/protrusion (lamellipodia-based). **G)** Live imaging of cells in 3D CDM. Dotted line shows orientation of kymograph. Arrow, positive correlation between trafficking of Rab25 endosomes towards the plasma membrane and actin polymerisation/protrusion (filopodia-based). **H, I)** Correlation between the movement of Rab25 endosomes and the tracks of actin polymerisation events. **H)** Live imaging of cells on FN, dotted line shows orientation of kymograph. Boxed area, zoom inset showing timelapse frames. Arrow, colocalizing fluorescent signal of NB^GFP^-mCherry-Rab25 endosomes (R25) and actin tracks visualized by LifeAct-iRFP670 (F-actin). **I)** Live imaging of cells on FN-treated 4 h with 1 µM Nocodazole. Dashed line, position of Rab25 endosome (R25) at the time 0. Colour-grade (cyan-red) LUT image show Rab25 endosome movement (R25) and F-actin fiber polymerisation in time not blocked by depolymerisation of microtubules. Arrow, colocalizing fluorescent signal of NB^GFP^-mCherry-Rab25 endosomes (R25) and actin tracks.

**Figure S2.**
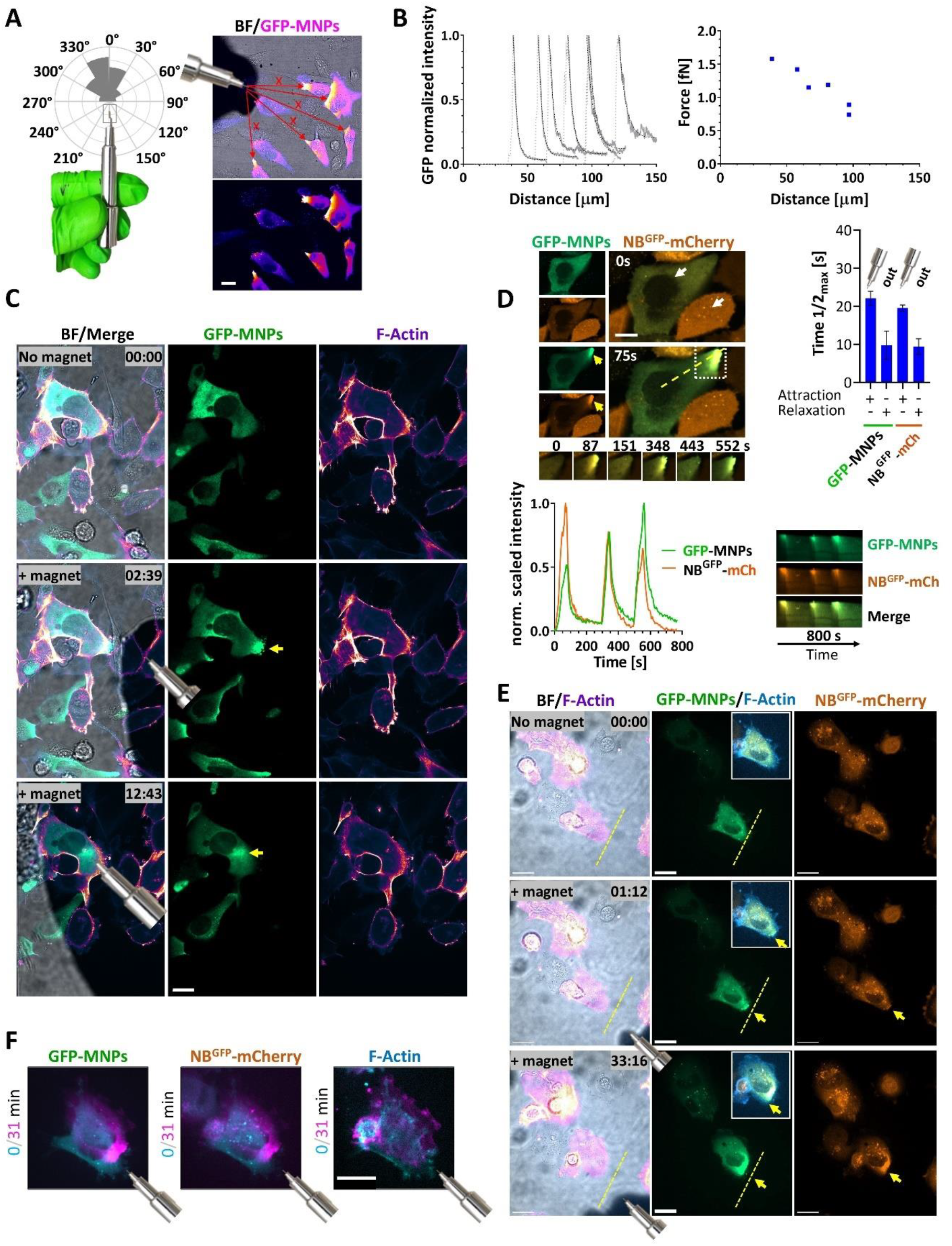
Magnetic manipulation of GFP-MNPs and NB^GFP^-mCherry inside living cells. **A-F)** Images generated by confocal spinning-disk microscopy (live cells). GFP-MNPs delivered by microinjection to cells on FN-coated coverslips. Brightfield (BF). **A-B)** Steady-state GFP-MNPs gradient profiles inside the cytoplasm of living A2780 cells generated by a magnetic gradient. **A)**Confocal spinning-disk live cell images of A2780 and angle between micro-magnet and centre of the microinjected cells with observed attraction gradient, N = 3 (n = 40). Scale bar 20 μm. **B)** The decay into the cytoplasm was fitted by an exponential decay to estimate the force exerted on the GFP-MNPs (see Methods, Equation 1) and plotted as function of magnetic tip distance from cells. **C)** Key time-lapse image frames showing magnetic manipulation of GFP-MNPs inside living A2780 cells stably expressing Lifeact-iRFP670 (F-actin). Shadow in brightfield indicates magnetic tip (exact position indicated by cartoon). Arrow, GFP-MNPs enrichment (see movie S1). Scale bar 20 μm. **D-E)** A2780 cells stably expressing NB^GFP^-mCherry (amber), Lifeact-iRFP670 (F-actin, blue or gem LUT) microinjected with GFP-MNPS (green). Movies S11-S13 (S12 shows magnetic manipulation across whole cells) accessible via https://doi.org/10.6084/m9.figshare.22155083. **D)** Magnetic attraction and release kinetics of NB^GFP^-mCherry-(X) and GFP-MNPs in living cells; individual representative frames (scale bar 10 μm) from highlighted white box, kymographs from dashed line and profiles (bottom left) determined from the changes in fluorescence intensity across kymograph (black arrow; normalized 0-1 scaled fluorescent intensities shown), exponentially fitted and attraction and relaxation kinetics quantified (right) as time needed to reach 50 % of maximal gradient of the steady-state (max; top right); all conditions n = 3 (N = 3). Out: no magnet. Arrows, accumulation of NB^GFP^-mCherry in the cytoplasm (out from nucleus) in GFP-MNPs positive cells. **E-F)** Representative example of ctrl migrating cell with magnetically attracted GFP-MNPs/NB^GFP^-mCherry, arrow. Magnetic tip indicated by shadow and/or cartoon. Dashed line, cell edge (F-actin) at the time 0. Inset, merge GFP-MNPs/NB^GFP^-mCherry/F-actin. Scale bar 20 μm. **F)** MIPs used for scoring protrusion growth (F-actin) shown in Fig. 1D.

**Figure S3.**
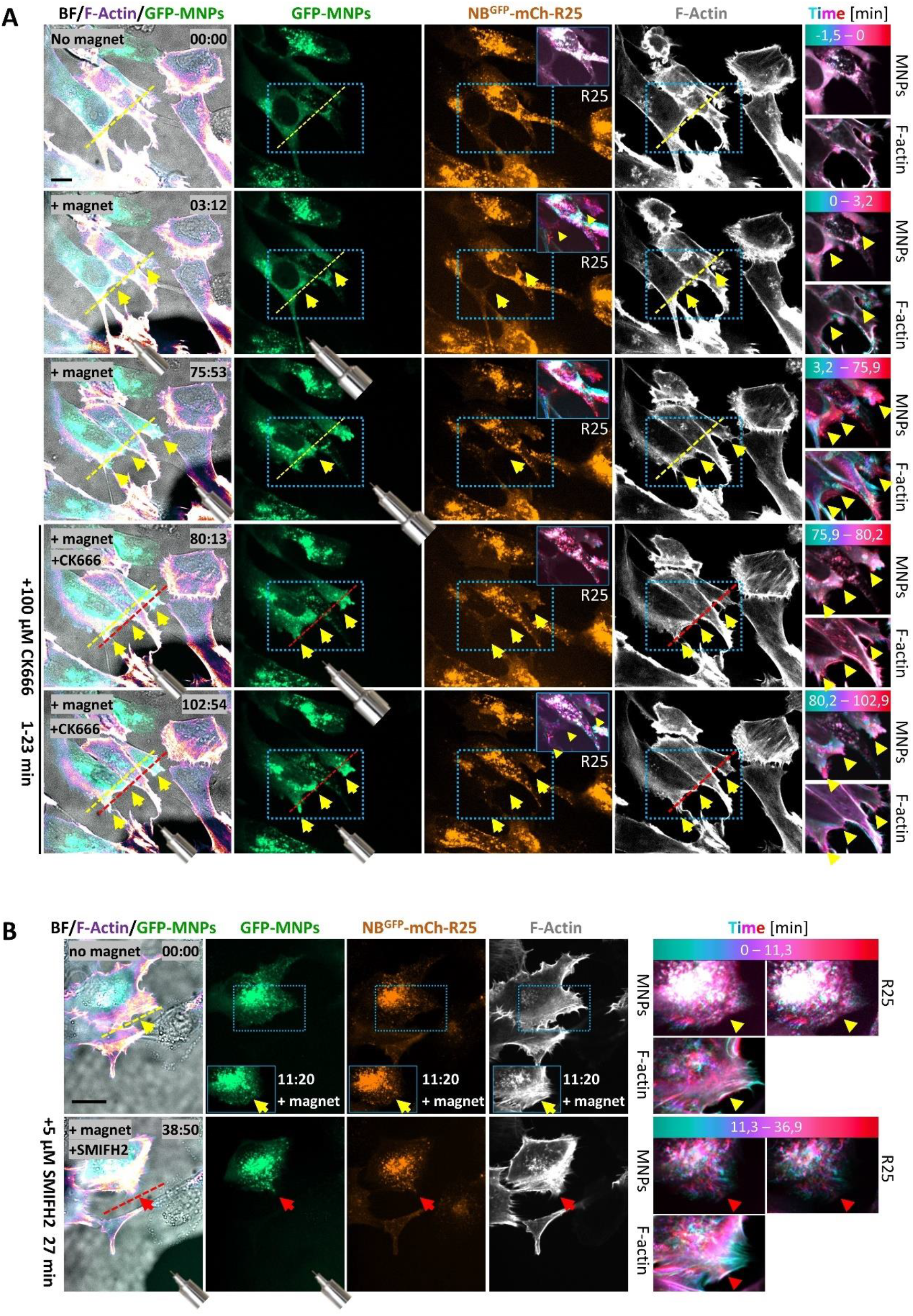
Remote manipulation of Rab25 endosomes controls protrusion outgrowth in an Arp2/3 independent and formin-dependent manner. **A, B)** Representative spinning-disk confocal live cell imaging, A2780 DExCon-modified NB^GFP^-mCherry-Rab25 cells dox pre-treated (>94 h; 250 ng/ml) stably expressing Lifeact-iRFP670 (F-actin), microinjected with GFP-MNPs. The effect of Rab25 endosomes remotely redistributed to promote protrusions is not inhibited by **A)** CK666 treatment (100 μM, see movie S14 accessible via https://doi.org/10.6084/m9.figshare.22155083, but blocked with **B)** SMIFH2 (5 uM; see movie S15 accessible via https://doi.org/10.6084/m9.figshare.22155083). Shadow in brightfield (BF) and cartoon indicates position of the magnetic tip. Yellow arrow, changes in GFP-MNPS/NB^GFP^-mCherry fluorescent signal or protrusion growth (F-actin). Red arrow, protrusion retraction/no response (F-actin). Boxed area, cyan-red LUT illustrates changes in GFP-MNPs and vesicle distribution (R25) and protrusion changes (F-actin) over time through colour grading. Yellow dashed line, cell edge (F-actin) at the time 0. Red dashed line, cell edge (F-actin) when added inhibitor. FN-coated coverslips, Scale bar 20 μm.

**Figure S4.**
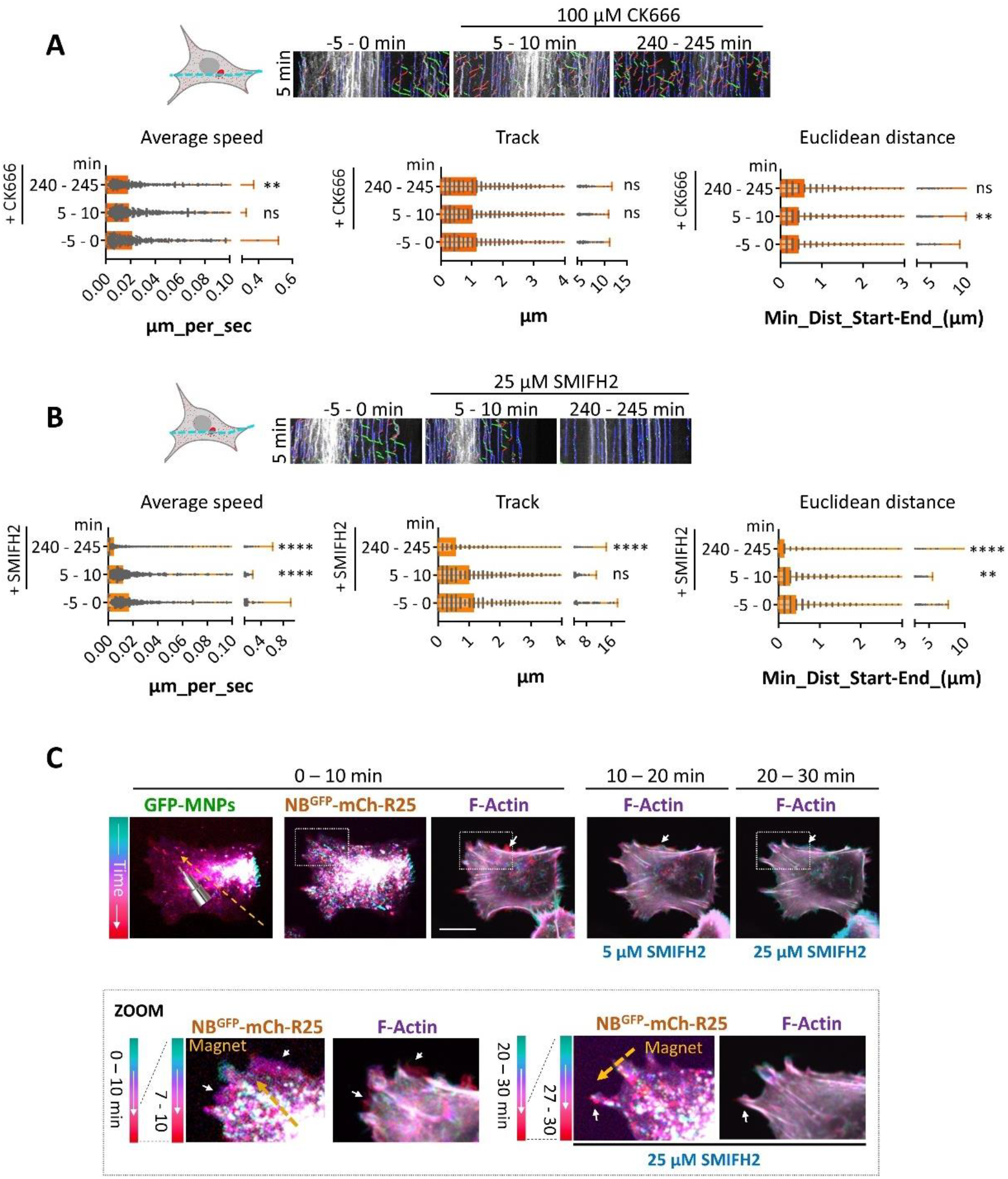
Active movement of Rab25 endosomes in A2780 cells requires formins, but not Arp2/3. **A, B, C)**Confocal spinning-disk live cell imaging of A2780 DExCon-modified NB^GFP^-mCherry-Rab25 (R25; dox treated > 48 h, 250 ng/ml) stably expressing Lifeact-iRFP670 (F-actin) on FN. **A, B)** AI based tracking of vesicles (NB^GFP^-mCh-R25) movement (labelled blue, static; brown, towards rear; green, towards front) using kymographs (generated from dashed line, schematic cartoon) and Kymobutler plugin before and after treatments: **A)** CK666 (100 µM) or **B)** SMIFH2 (25 µM) as indicated. Representative kymographs are shown, -5 – 0 min/5-10 min same cell; 240-245 min different cell. Average speed, vesicle total track or euclidean distance quantified; *n* > 9 cells/condition (N = 3); Anova on ranks, Dunn’s test (compared to no treatment). \*\**P* < 0.01; \*\*\**P* < 0.001; \*\*\*\**P* < 0.001. **C)** Magnetic relocalization of GFP-MNPs and Rab25 endosomes (NB^GFP^-mCh-R25) towards upper part of the cell, cells treated by SMIFH2 as indicated and position/movement of magnet (cartoon) indicated by orange arrows. Boxed area, cyan-red LUT illustrates changes in GFP-MNPs and vesicle distribution (R25) and protrusion changes (F-actin) over time through colour grading. Representative cyan-red LUT images of F-actin show actin-dependent protrusion dynamics before and after SMIFH2 treatment (N = 3). Boxed area, zoom inset bellow. White arrow, protrusion and vesicle movement highlighted. FN-coated cover-slip. Scale bar 20 μm.

**Figure S5.**
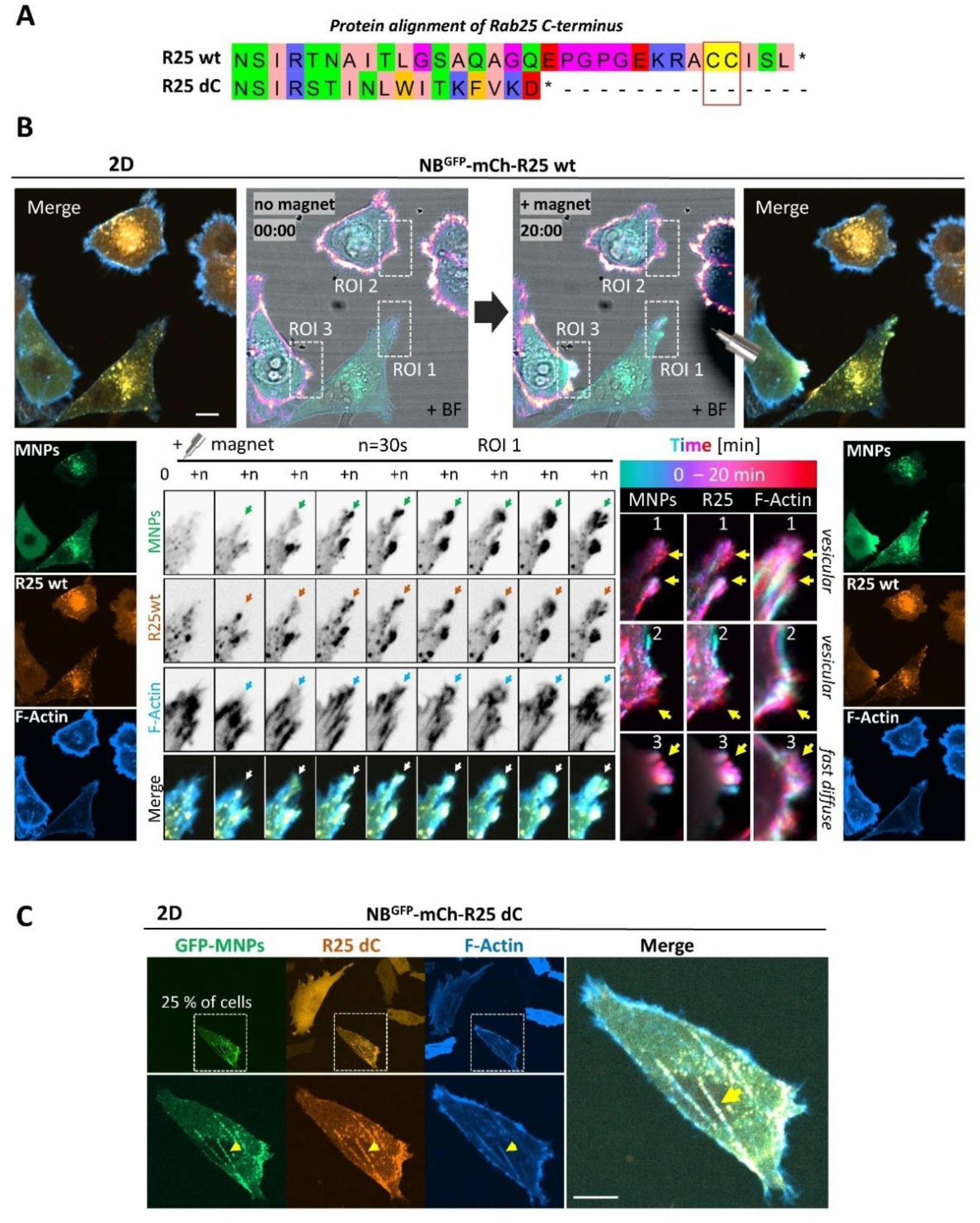
Rab25 promotes actin-dependent protrusion growth. **A)** Protein alignment of C-terminus of Rab25 wt and dC mutant. Zappo color coding in Jalview. Red rectangle, membrane attachment site (CC). **B)** Magnetic attraction of Rab25 (R25) endosomes promotes protrusion growth independently of initial cell polarity. Confocal spinning-disk live cell timelapse imaging of A2780 cells stably expressing NB^GFP^-mCherry-Rab25 wt (biop LUT amber) with Lifeact-iRFP670 (F-actin, biop LUT azure) on FN, microinjected with GFP-MNPs (biop LUT spring green). Magnetic tip visible as shadow in brightfield (BF) and indicated by cartoon. Boxed area, Region of interest (ROI) 1-3 shown as cyan-red LUT illustrating changes in GFP-MNPs and vesicle distribution (R25) and protrusion changes (F-actin) over time through colour grading. ROI 1 also shown as individual time frames. Arrows, correlation of changes in individual channels. See movie S17 accessible via https://doi.org/10.6084/m9.figshare.22155083. Scale bar =10 μm. **C)** Confocal spinning-disk live cell imaging of A2780 cells stably expressing NB^GFP^-mCherry-Rab25 dC (biop LUT amber) with Lifeact-iRFP670 (F-actin, biop LUT azure) on FN, microinjected with GFP-MNPs (biop LUT spring green). Representative example of ∼25 % cells with Rab25 dC signal co-aligned with F-actin stress fibers induced by GFP-MNPs microinjection, arrow (not visible in Ctrl NB^GFP^-mCherry expressing cells). Scale bar =10 μm.

**Figure S6.**
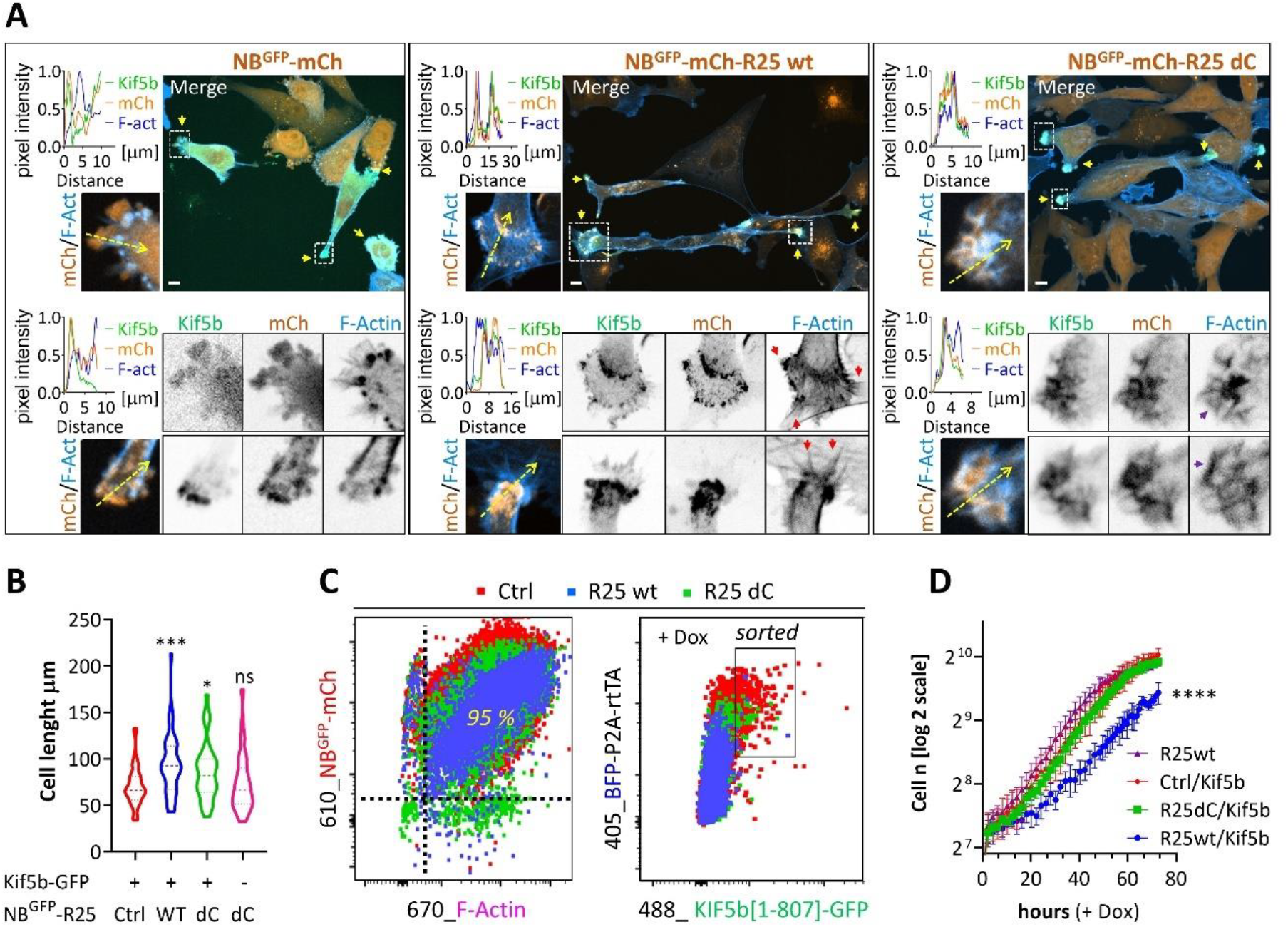
Rab25 targeted to cell periphery via Kif5b locally stimulates actin polymerisation. **A)** Confocal spinning-disk live cell imaging of A2780 stably expressing NB^GFP^-mCherry (Ctrl, left) or fused with Rab25 wt (middle) or Rab dC (right) with Lifeact-iRFP670 (F-actin) prepared by lentiviral transduction (sorted for matched expression levels) and truncated GFP-fused Kif5b [1-807] generated by sleeping Beauty system constitutively expressing BFP, rtTA and puromycin resistance gene ((Kowarz *et al*., 2015); selected by puromycin). All dox treated (48 h; 500 ng/ml). Merge, MIPs. Boxed area, zoom inset shown as one Z plane, two examples for each variant shown. Dotted line, line scan profile of normalized 0-1 scaled fluorescence intensities. Yellow arrows, Kif5b positive protrusions. Red arrows, filopodia. Purple arrows, lamellipodia-based structures. FN-coated µ-Plate 96 plate (#1.5 IbiTreat). Scale bar 10 μm. Cell length quantified in **B)** based on F-actin as maximum ferret diameter. Only cells co-expressing BFP/GFP (readout of Kif5b positivity) were analysed (except Kif5b negative Rab25 dC cells). One-way ANOVA analysis Dunnet post hoc test (compared to Ctrl). N > 42 cells all conditions; N = 3. \**P* < 0.05; \*\*\**P* < 0.001. C) Cells described in **A)** sorted for matched GFP-KIF5b expression levels as indicated. D) Representative example of proliferation rate of cells described in **C)** analysed by eSight real-time proliferation assay; N = 4 (all dox treated; 500 ng/ml). Cell numbers (n) derived from brightfield trained object masks, normalized for matched numbers at the time 0. The linear part of the cell proliferation rate (10-50 h) plotted by log2 was fitted by simple linear regression and compared using an Analysis of Covariance (ANCOVA). \*\*\*\**P* < 0.001 (between Rab25 wt ± Kif5b).

**Figure S7.**
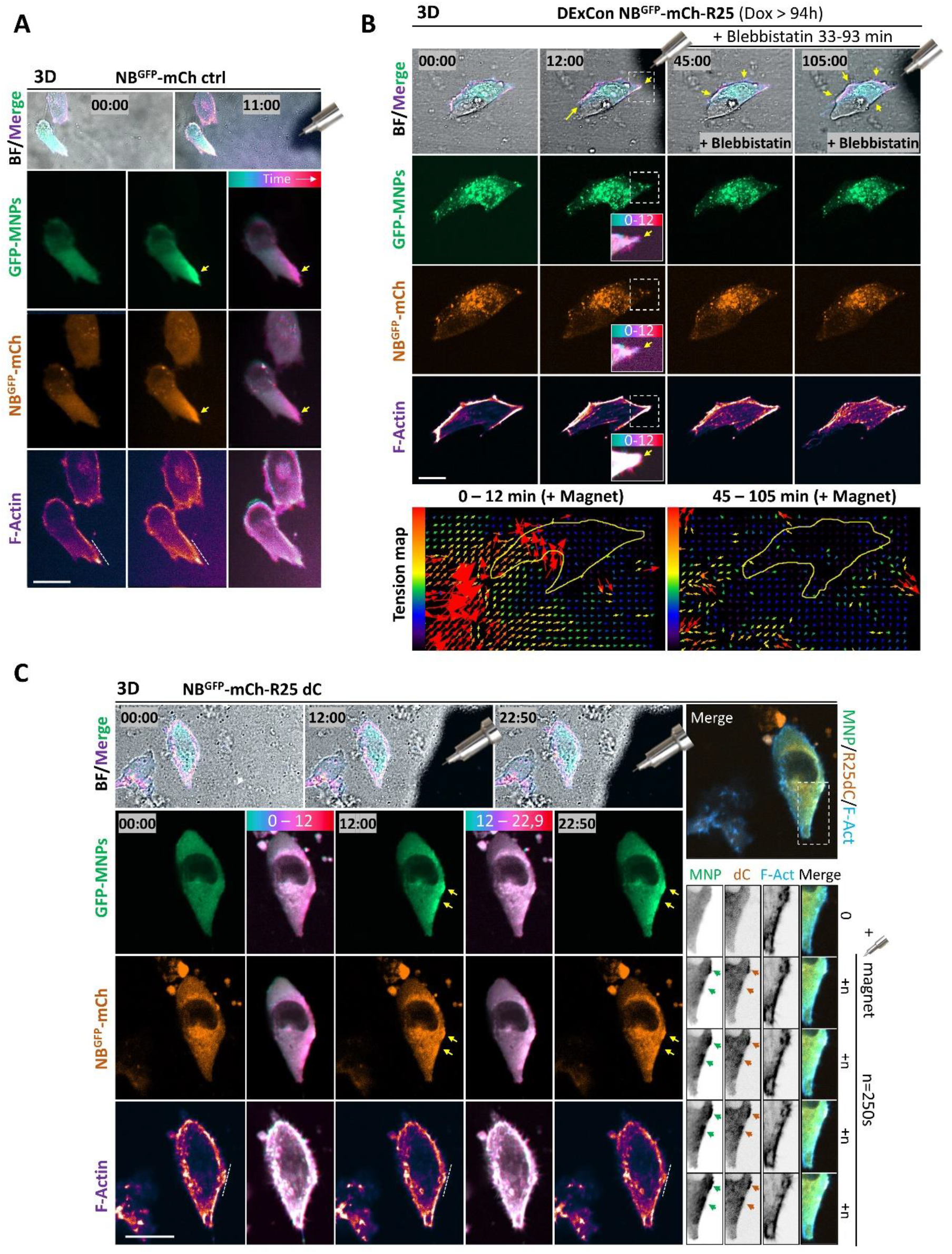
Rab25’s ability to interact with vesicles enables the mechanosensing of magnetic forces by MNP-bound Rab25 endosomes and is essential to promote protrusion growth in CDM. **A-C)** Representative images from confocal spinning-disk live cell imaging of cells in 3D CDM; Scale bar 20 μm; GFP-MNPs delivered by microinjection and position of magnetic tip visible as shadow in brightfield (BF) or indicated by cartoon. The cyan-red LUT illustrates changes in NB^GFP^-mCherry-X/GFP-MNPs distribution and cell shape (F-actin) over time through colour grading. Representative example; N = 3 (4 for Rab25 dC shown in **C)**. A) A2780 stably expressing NB^GFP^-mCherry A2780 with Lifeact-iRFP670 (F-actin). Dashed line, cell edge (F-actin) at the time 0. Arrow, gradient of fluorescent signal. B) Confocal spinning-disk live cell imaging of A2780 DExCon-modified NB^GFP^-mCherry-Rab25 (R25; dox treated > 94 h, 250 ng/ml)stably expressing Lifeact-iRFP670 (F-actin). Cell adaptation to mechanosensing of magnetic force by MNP-bound Rab25 endosomes (time 0-12 min) is perturbed by Blebbistatin (5 µM) treatment (18-105 min), no magnet in between (30 min Blebbistatin pre-incubation). Arrows, cell shape changes (shrinking/growth 0-12 min or swelling 45-105 min). Relative stress maps are displayed using as a vectorial plot with red-blue LUT, size and direction of vectorial arrows illustrate force distribution exterted on CDM. C) A2780 cells stably co-expressing NB^GFP^-mCherry-Rab25 dC and Lifeact-iRFP670 (F-actin). Dashed line, cell edge (F-actin) at the time 0. Arrow, gradient of fluorescent signal. Movies S18-20 accessible via https://doi.org/10.6084/m9.figshare.22155083.

**Figure S8.**
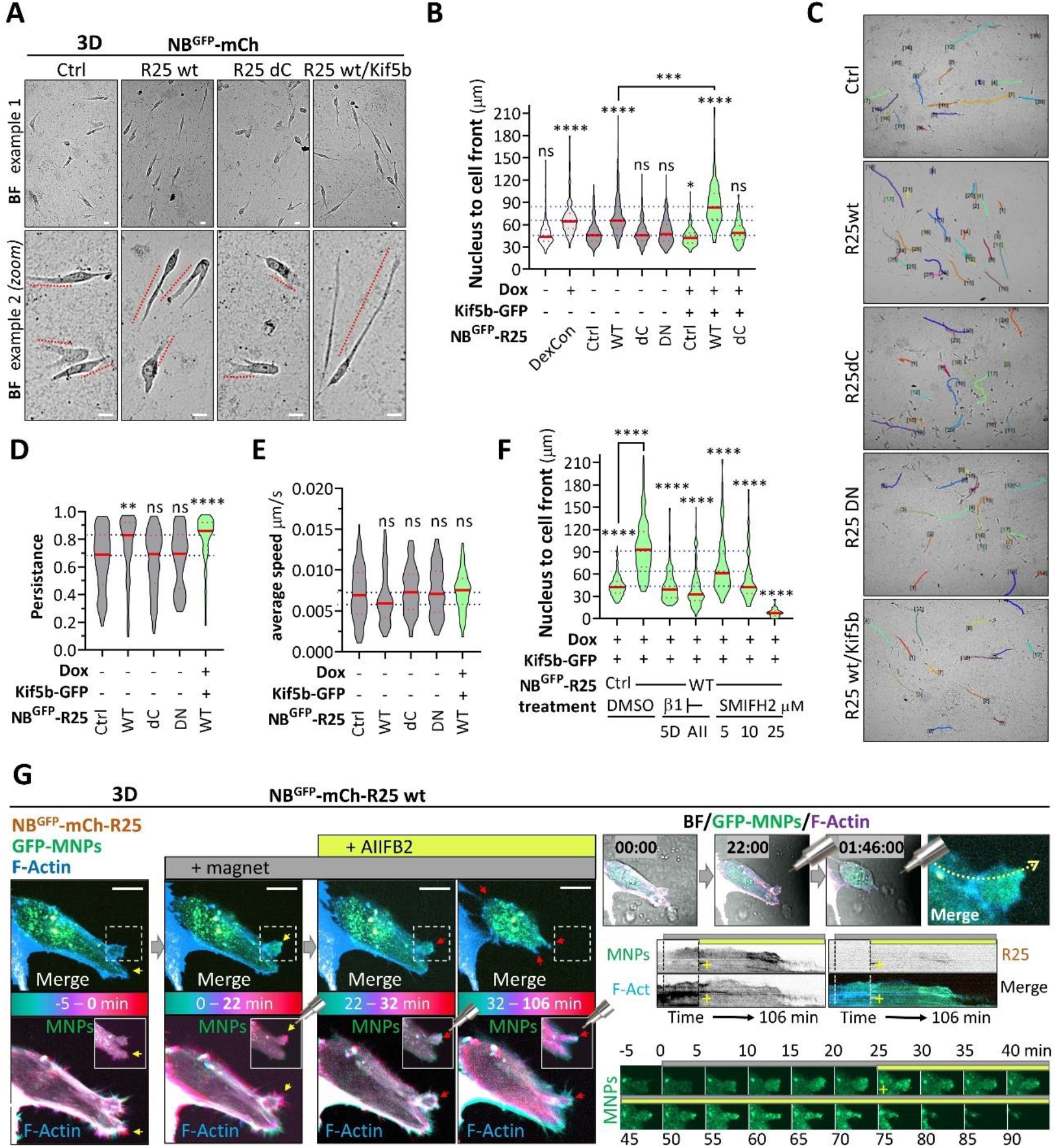
Rab25’s ability to interact with integrin β1-containing recycling vesicles is essential to promote protrusion growth in CDM. **A-F)** Cell migration in 3D CDM; A2780 DExCon-modified NB^GFP^-mCherry-Rab25 (± dox treated 48 h, 250 ng/ml) or A2780 cells stably co-expressing Lifeact-iRFP670 (F-actin) with NB^GFP^-mCherry (Ctrl) or fused with Rab25 wt or Rab dC and truncated GFP-fused Kif5b [1-807] sorted for matched GFP-KIF5b expression levels (dox treated 24-48 h, 500 ng/ml) shown in fig. S6C. **A)** Widefield live cell imaging (brightfield (BF)). CDM (in for 8-24h). Zoom inset from diff. area. Scale bar 10 μm. Dotted line, length of pseudopodial protrusion (nucleus to the cell front of migrating cells) quantified in **B)** and shown as violin plot; n = 150-300 cells/condition, N = 3-4. **C)** Representative example of cell tracks, quantification of migration persistence **D)** and average speed **E)**; cells in CDM for 8-24h; n > 70, N = 3. **F)** Quantification of length of pseudopodial protrusion (nucleus to the cell front of migrating cells) of cells described in A) ± treatment shown as violin plot. Cells in CDM for 8-24h; n > 150 cells/condition (50 for 25uM SMIFH2 as majority of cells are immobile); N = 3 if not stated otherwise (vehicle or no treatment N = 4); SMIFH2 added 4h after spreading (μg/ml as indicated); Integrin β1 blocking antibodies added before seeding cells into CDM: AIIB2 (AII) 10 μg/ml (N = 3); 5PD2 (5D) 10-15 ug/ml (N = 2). All graphs Anova on ranks, Dunn’s test (compared to ctrl or Rab wt/Kif5b as indicated); \**P* < 0.05; \*\**P* < 0.01; \*\*\**P* < 0.001; \*\*\*\**P* < 0.001. **G)**. Representative images from confocal spinning-disk timelapse images of A2780 NB^GFP^-mCherry-Rab25 wt cell expressing Lifeact-iRFP670 (F-actin), migrating in CDM (3D), microinjected with GFP-MNPs before (-5-0 min) or after GFP-MNPs magnetically attracted towards magnetic tip (0-106 min) visible as shadow in brightfield (BF); see cartoon).Rab25 promoted protrusion growth (0-22 min) is blocked by integrin β1 blocking antibody (+; yellow rectangle) treatment (AIIFB2, 10 μg/ml; 22-32 min) followed by protrusion retraction/cell rounding (32-106 min) despite sustained GFP-MNPs magnetic attraction (grey rectangle, see timelapse frames and kymographs). The cyan-red LUT illustrates changes in protrusion growth (F-actin) over time through colour grading (GFP-MNPs inset; cropped cell front). Yellow arrow, protrusion growth. Red arrow, protrusion no change/retraction. Boxed area, individual representative time frames. Dotted line, kymograph. N = 3. Scale bar 10 μm. See movie S5.

**Figure S9.**
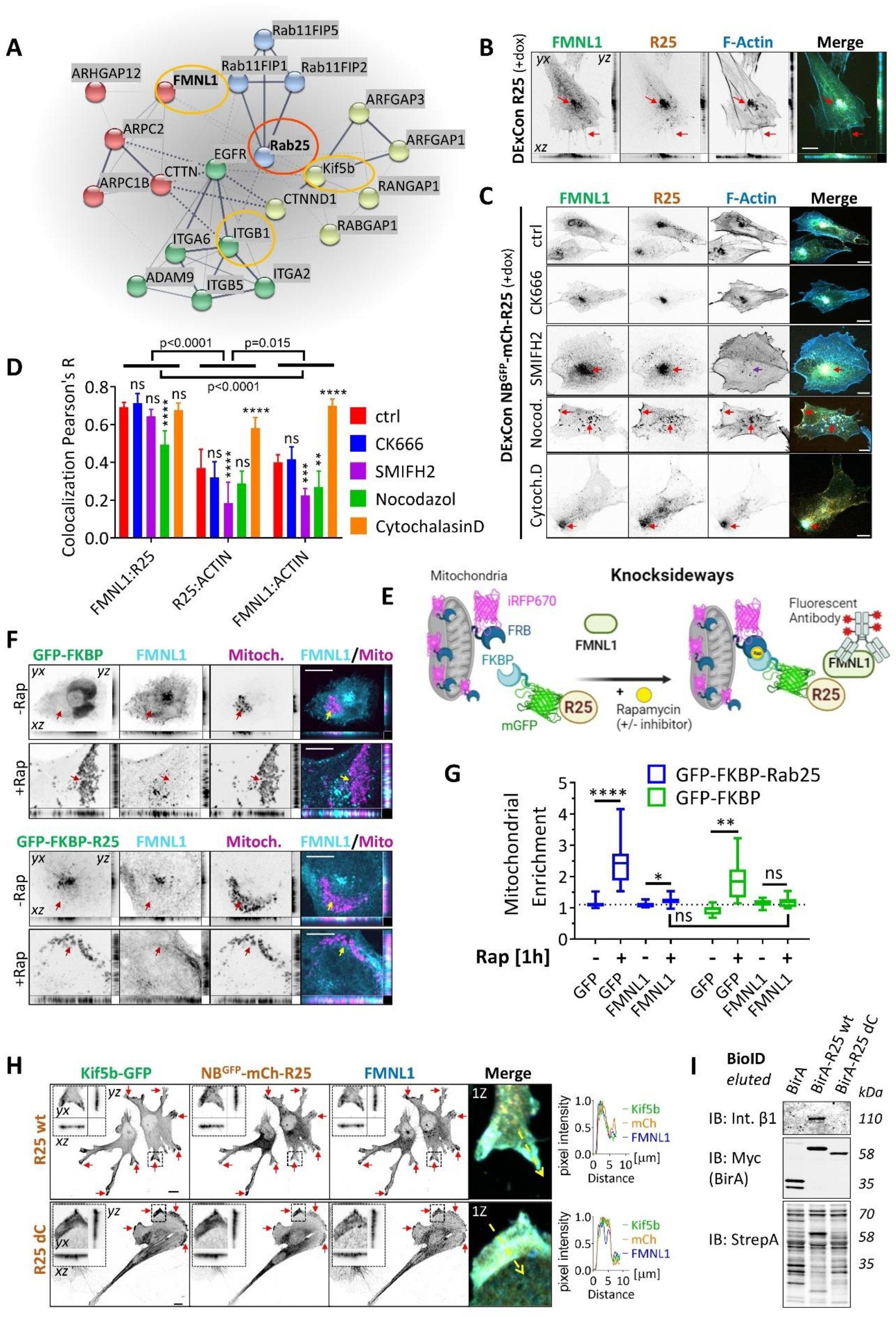
The interaction between Rab25 and FMNL1 is of a transient nature, but FMNL1 and integrin β1 are cargoes of Rab25 vesicles. **A)** Rab25-BioID (proximity labelling) dataset (Wilson *et al*., 2023) re-analysis (see methods). Rab25-BirA (red circle) enriched cell matrix interactors (green); Actin/GTPase regulators (red and white), Rab effectors (blue) are represented using STRING protein-protein interaction analysis (https://string-db.org/). Interesting targets are in circled in yellow (ITGB1/integrin β1; FMNL1; KIF5b). **B-C)** Representative confocal spinning-disk images of A2780 DExCon-modified NB^GFP^-mCherry-Rab25 cells (mCherry) on FN-pre-treated with dox (> 94 h; 250 ng/ml) immunolabeled for FMNL1 (rabbit anti-FMNL1; Alexa-488) and stained for F-actin (Phalloidin-Alexa633). Scale bar=10 μm. B) MIP with cross-sections. Arrows, filopodia or perinuclear recycling compartment. **C)** MIPs. Ctrl (no treatment); CK666 (100 μM; 4h); SMIFH2 (25 μM; 4h); Nocodazol (1 μM; 4h); Cytochalasin D (100 ng/ml (200 nM); 4h). Red arrows, Enriched colocalized FMNL1/Rab25/F-actin signal (purple arrow; missing F-actin signal upon SMIFH2 treatment). FMNL1/Rab25/F-actin signal colocalization in D) using Pearson’s R. Two-way ANOVA analysis Tukey post hoc test (compared with ctrl or cross-correlation of the similarity of treatment-induced changes between FMNL1:R25, R25:ACTIN, and FMNL1:ACTIN, as indicated); \*\**P <* 0.01; \*\*\**P <* 0.001; \*\*\*\**P <* 0.001. **E)** Schematic diagram (created with BioRender.com) of knock-sideways experiment whose representative images are shown in **F)** and quantified in G) as colocalization of GFP signal or proportional co-enrichment of FMNL1 in mitochondria (mask from mitochondrial targeting sequence (Mito) fused with iRFP670-FRB). A2780 co-transfected with Mito-iRFP670-FRB (iRFP670) and FKBP-GFP-Rab25 wt (GFP), ± Rapamycin (Rap; 1h, 200 nM), fixed and immunolabeled for FMNL1 (rabbit anti-FMNL1; cy3). MIP with cross-section is shown. Arrows, mitochondria based on Mito-iRFP670-FRB (iRFP670) channel. Scale bar 10 μm. **G)** One-way ANOVA analysis Tukey post hoc test (compared between GFP; GFP-Rab25 ± rap or between FMNL1 ± rap); \**P <* 0.05; \*\**P <* 0.01; \*\*\*\**P <* 0.001. H) Representative confocal spinning-disk images of A2780 cells stably co-expressing NB^GFP^-mCherry-Rab25 wt or Rab dC and truncated GFP-fused Kif5b [1-807] sorted for matched GFP-KIF5b expression levels (dox treated 24-48 h, 500 ng/ml) immunolabeled for mCherry (rat anti-RFP; Alexa-555); GFP (mouse anti-GFP; Alexa-488), and FMNL1 (rabbit anti-FMNL1; Alexa-633). MIPs. Boxed area, zoom inset (MIP) shown with cross-sections or shown as merge (1Z). Dashed line, line scan profile of normalized 0-1 scaled fluorescent intensities from 1Z plane. Arrows, protrusions (large filopodia based with vesicular FMNL1 (Rab25 wt) or lamellipodia based (Rab25 dC)). Scale bar=10 μm. I) A2780 stably expressing BirA or BirA fused with Rab25 WT or Rab25 dC mutant cultured with biotin (1 μM biotin, 16 h). Lysates equalized to total protein amount and biotinylated proteins pulled down with Streptavidin beads. Immunoblots of pulled down integrin β1, BirA (Myc epitope), Streptavidin (StrepA) as loading control. Fluorescent antibodies shown as black and white.

**Figure S10.**
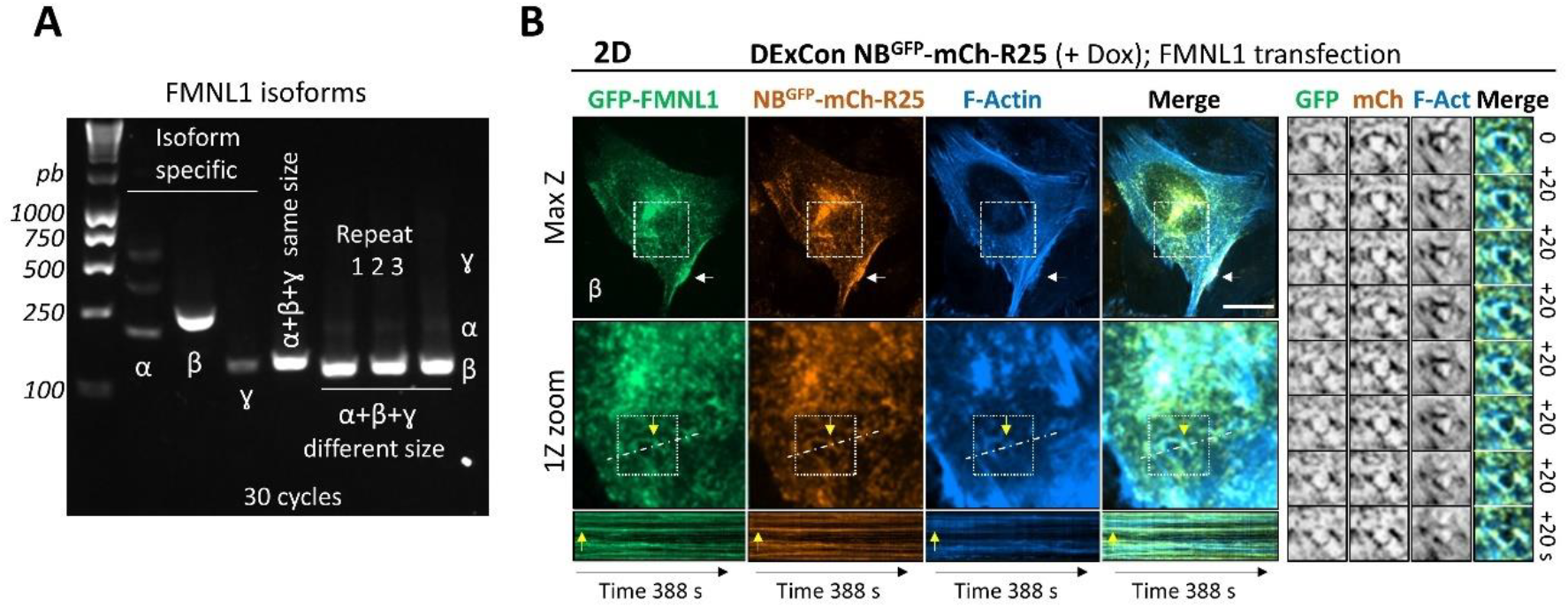
FMNL1 β is major isoform in A2780 cells and its interaction with Rab25 associated endosomes is functional. **A)** Semiquantitative PCR (30 cycles). Agarose gel, PCR products of FMNL1 isoforms (α, β, γ) amplified using combination of non-distinguishing, isoform specific or semi-specific primers leading to different sizes of PCR products of FMNL1 isoforms (3 independent biological repeats shown). **B)** Representative confocal spinning-disk live cell imaging of A2780 DExCon-modified NB^GFP^-mCherry-Rab25 (R25; dox treated 48 h, 250 ng/ml) stably expressing Lifeact-iRFP670 (F-actin) transfected by GFP fused FMNL1 β isoform. MIP or 1Z plane shown as indicated. Boxed dashed area, zoomed inset. Solid boxed area, individual timelapse frames (n + 20s). Dashed line, kymograph. White arrow, FMNL1/Rab25/F-actin colocalizing signal in protrusion. Yellow arrow, FMNL1/Rab25/F-actin stable endosomal colocalization. Scale bar 20 µm.

**Figure S11.**
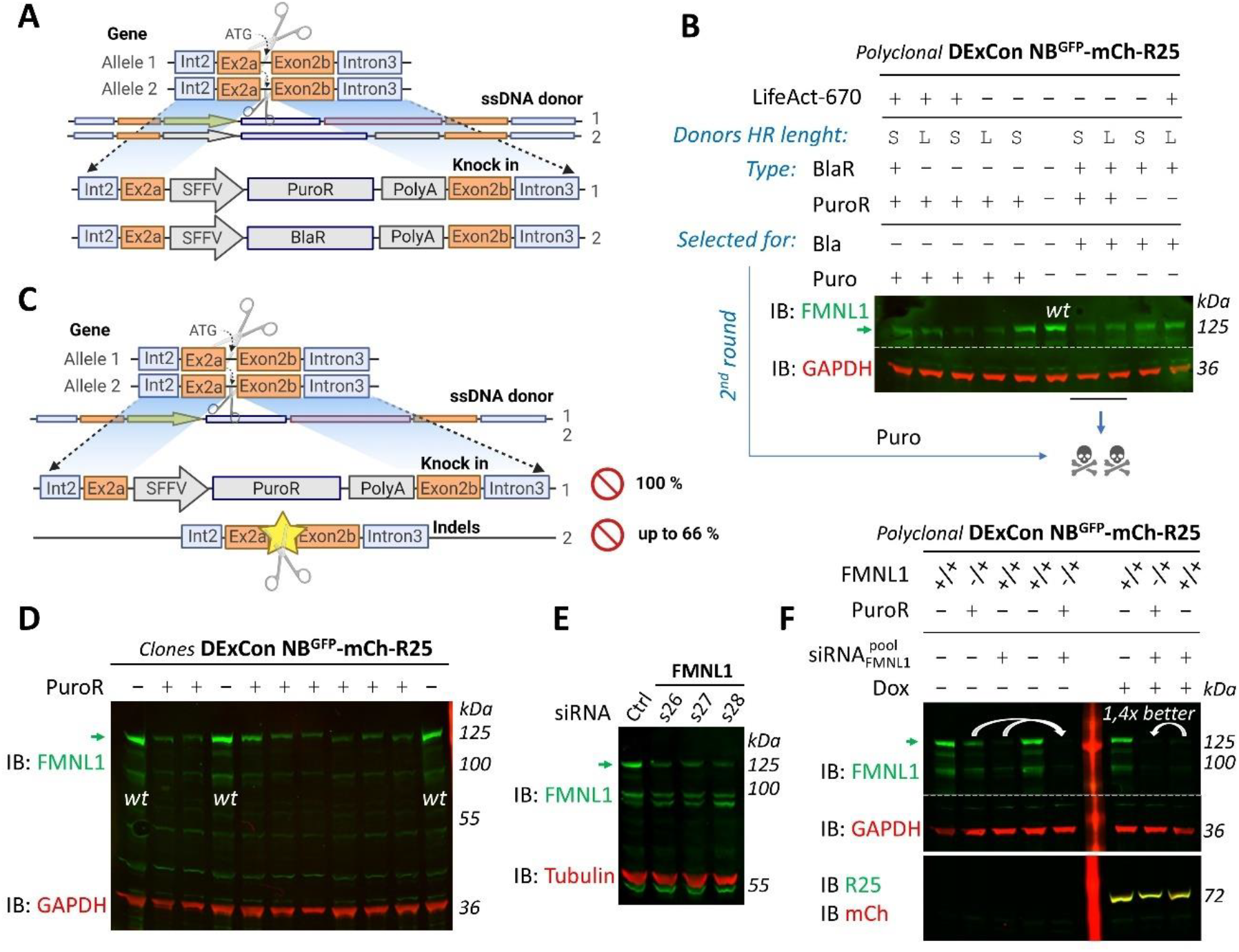
Efficient depletion of FMNL1 in A2780 cells. **A, C)** Strategy to inactivate FMNL1 gene using CRISPR/Cas9 and **A)** combination of single strand donor (ssDNA) cassettes carrying SFFV promoter followed by gene for puromycin (PuroR) or blasticidine (BlaR) resistance, flanked by homologous FMNL1 exon2 sequences (HR, long (L; 530 b) or short (S; 300 b) used) or **C)** combination of ssDNA donor for homologous recombination of puromycin resistance into exon2 of FMNL1 with formation of non-specific indels by non-homologous end joining as indicated (see also Fig. 5A; for details Methods). Created with BioRender.com. **B)** Immunoblot of DExCon-modified NB^GFP^-mCherry-Rab25 cells ± Lifeact-iRFP670 modified by CRISPR/Cas9 RNP and different combination of FMNL1 specific donor ssDNA cassettes, selected for the resistance as indicated. Only FMNL1 heterozygotes (FMNL1-/+) observed, secondary complementary editing (FMNL1 blaR/+) of remaining 2^nd^ allele by ssDNA carrying puroR followed by puromycin treatment led to cell death. Overlay of FMNL1 and GAPDH (loading control), fluorescent antibodies. **D)** Representative immunoblot of clonally derived DExCon-modified heterozygous FMNL1 PuroR/+ NB^GFP^-mCherry-Rab25 cells (in total 33 clones) with residual FMNL1 expression oscillating around the FMNL1 wt size (arrow), indicating the presence of in-frame indels. Overlay of FMNL1 and GAPDH (loading control), fluorescent antibodies. E) Immunoblot of A2780 nucleofected (48 h after) with different variants of siRNA anti-FMNL1. Non-targeting siRNA used as control (see methods). Arrow, FMNL1. Overlay of FMNL1 and Tubulin (loading control), fluorescent antibodies. F) Comparison of gene editing, siRNA knockdown and siRNA/CRISPR combined strategy (arrows). Immunoblot of DExCon-modified NB^GFP^-mCherry-Rab25 cells FMNL1+/+ or FMNL1 PuroR/+ (-/+) nucleofected (5 days after) with siRNA pool (s26, s27, s28) anti-FMNL1, ± dox (48 h, 250 ng/ml) as indicated. Overlay of FMNL1 and GAPDH (loading control) or Rab25 (R25) and mCherry (mCH), respectively. fluorescent antibodies.

**Figure S12.**
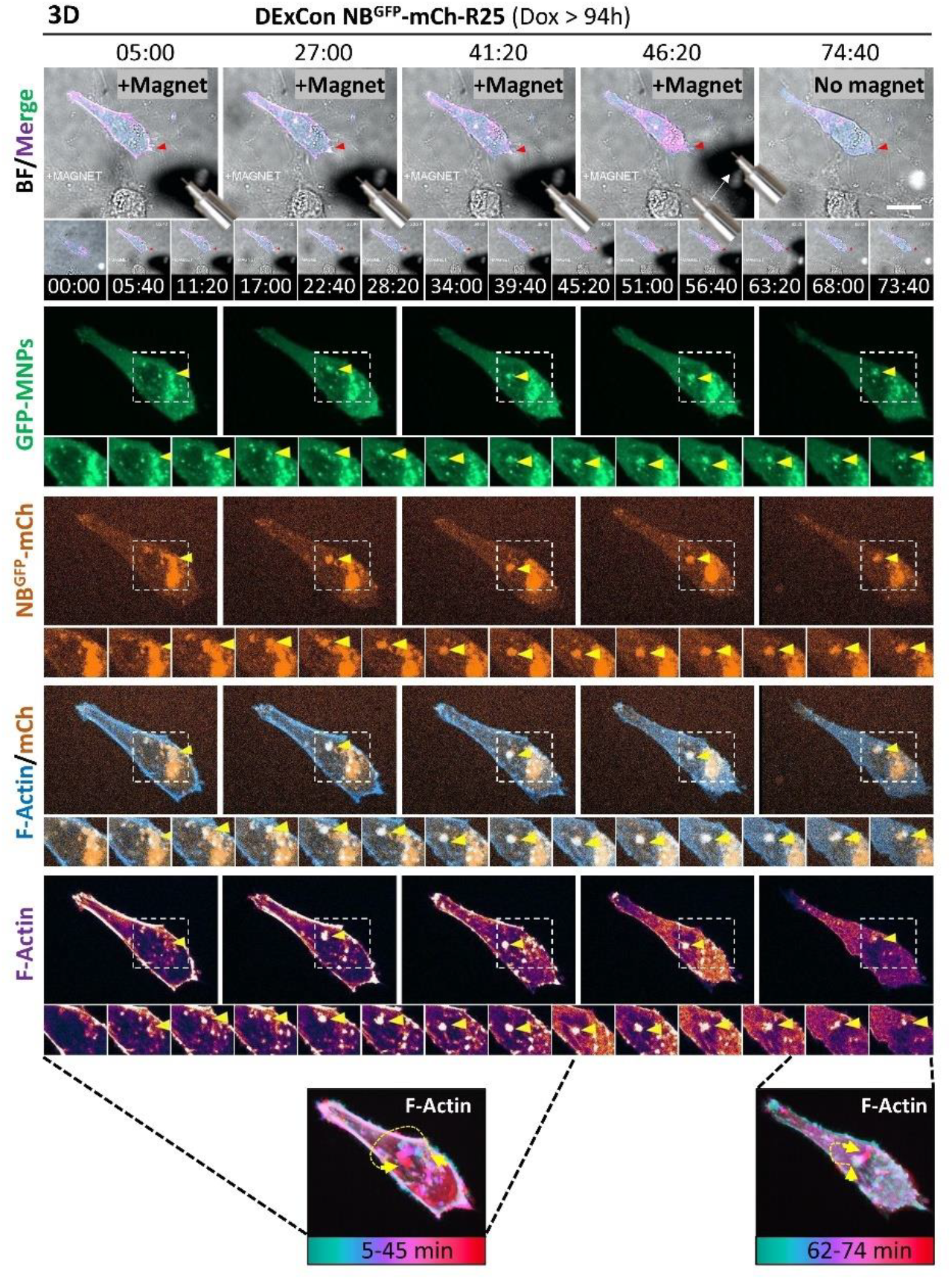
Magnetic relocalization of endosomal cluster via Rab25 locally induce actin polymerisation hot-spot. Representative images from confocal spinning-disk live cell imaging. A2780 DExCon-modified NB^GFP^-mCherry-Rab25 cells dox pre-treated (72 h; 250 ng/ml) expressing Lifeact-iRFP670 (F-actin) migrating in CDM (3D). GFP-MNPs delivered by microinjection. Magnetic attraction of GFP-MNPs/NB^GFP^-mCherry-Rab25 endosomal cluster towards magnetic tip (0-46 min) visible as shadow in brightfield (BF); see cartoon) promotes formation of co-moving actin polymerisation hotspot (arrow; F-actin shown as cyan-red LUT 5-45 min) followed by reverse movement and intensity decrease upon magnet removal (arrow; F-actin shown as cyan-red LUT 62-74 min). Boxed area, individual timelapse frames (n + 5 min 40s). Scale bar 20 μm. Movie S21 accessible via https://doi.org/10.6084/m9.figshare.22155083.

**Figure S13.**
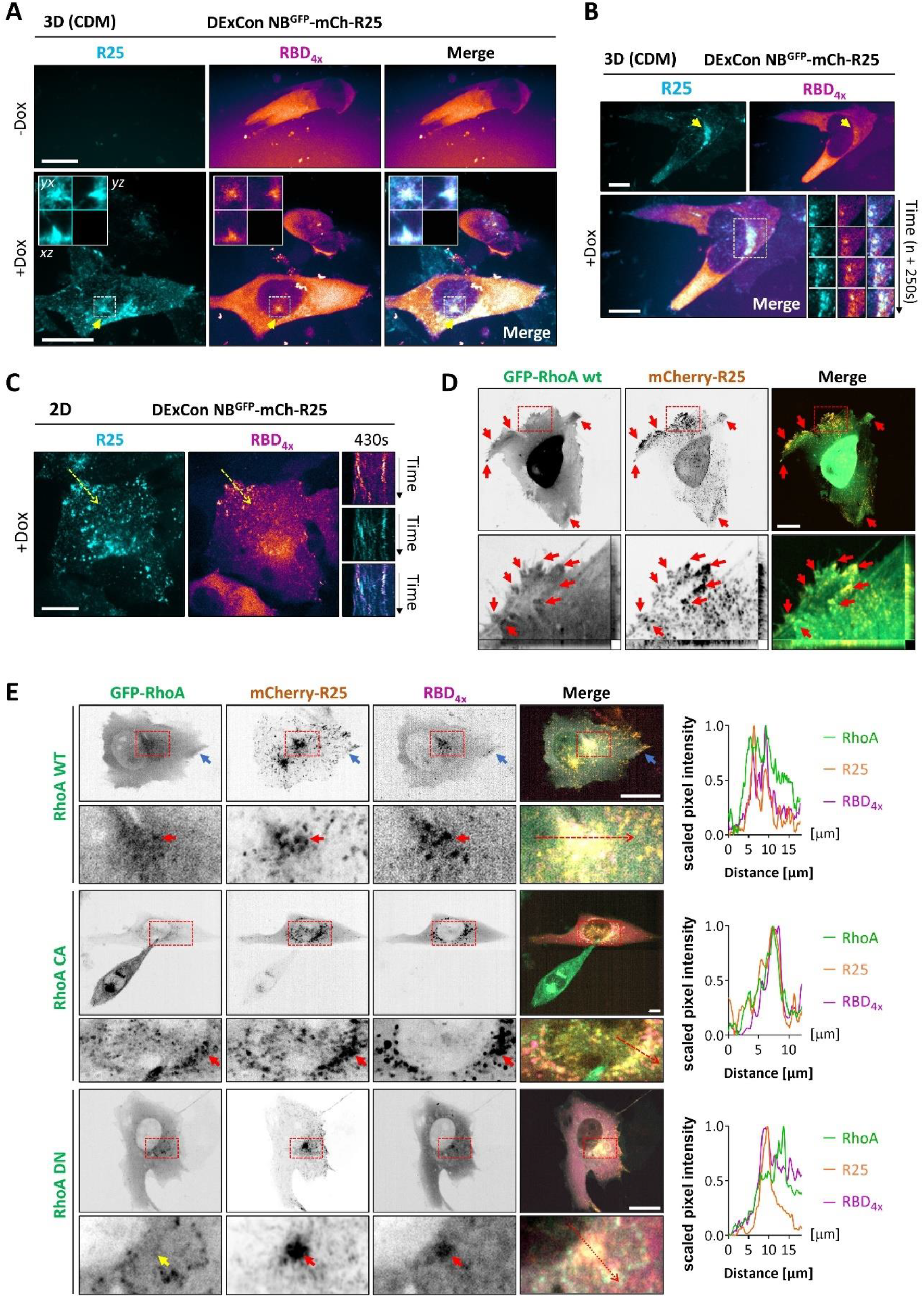
Specificity of iRFP670(3x)-RBD(4x) sensor to report RhoA activity and its colocalization with Rab25. **A-E)** Representative images from confocal spinning-disk live cell imaging. Scale bar 20 μm. A-C) A2780 DExCon-modified NB^GFP^-mCherry-Rab25 (R25; dox treated ≥ 48 h, 250 ng/ml) stably expressing active RhoA probe iRFP670_3x_-RBD_4x_ (RBD_4x_). A, B) 3D CDM. MIPs. Boxed area, zoom inset with cross-sections **A)** or timelapse frames (from 1Z plane; **B)**. Arrow, PNRC. C-E) FN-coated µ-Plate 96 plate (#1.5 IbiTreat). **C)** Line, kymograph area, zoom inset with cross-sections. Arrow, PNRC. **D)** A2780 cells co-transfected by GFP-RhoA wt and mCherry-Rab25 wt. MIPs. Boxed area, zoomed inset with cross-sections. Arrows, filopodia based protrusions. E) A2780 cells co-transfected by mCherry-Rab25 wt (R25), iRFP670_3x_-RBD_4x_ (RBD_4x_) and GFP fused RhoA wt or mutants (DN = dominant negative; T19N, CA = constitutively active; Q63L). MIPs. Boxed area, zoom inset. Blue arrow, RhoA/R25/RBD_4x_ signal at protrusion. Red arrow, RhoA/R25/BD_4x_ signal enriched at PNRC/endosomes (RhoA WT or CA, but missing from DN, yellow arrow). Red dashed line, line scan profile of normalized 0-1 scaled fluorescence intensities.

## Supplementary information

Plasmid maps with sequences generated in this study can be download from https://doi.org/10.6084/m9.figshare.27084205. List of material and other sequences are provided in excel file S1 or accessible from https://doi.org/10.6084/m9.figshare.27084217. Plasmids can be provided upon reasonable request from corresponding authors or will be available from Addgene. All representative full-resolution S1-S10 movies (Full HD) have been deposited in the Figshare repository and can be accessed via from https://doi.org/10.6084/m9.figshare.22155083, as can the additional movies S11-S22 which could not be incorporated directly with this study. Full description of included movies is in the accompanying movie legend.

Excell file S1 with list of primers used for ssDNA preparation, screening of knock in outcomes and semi-quantitative PCR, synthetized DNA sequences, crRNA sequences and list of all plasmids, regencies and antibodies used in this study.

## Supplementary movie legends

Representative live-cell imaging movies (spinning disk confocal) are separate files relevant to the study. Movies S1-S10 show spatiotemporal magnetic control of GFP-MNPs inside living A2780 cells expressing NB^GFP^-mCherry (ctrl) or NB^GFP^-mCherry fused with different variants of Rab25 stably co-expressing LifeAct-iRFP670 (F-actin) or active RhoA probe iRFP670_3x_-RBD_4x_ (RBD_4x_), as detailed in the accompanying movie legends. For full-resolution and additional movies see https://doi.org/10.6084/m9.figshare.22155083.

### Supplementary movie S1_fig. S2C

Timelapse video (3i Marianas spinning disk) of remote magnetic manipulation of GFP-MNPs (biop-SpringGreen LUT) inside living A2780 cells stably expressing LifeAct-iRFP670 (F-actin, gem LUT) on FN-coated coverslips. Merge: GFP-MNPs/F-actin/Brightfield, left. Magnetic tweezers/tip visible as shadow in brightfield. GFP-MNPs relocalization to various subcellular locations, red arrow. Timelapse covers total 18 min 33 s with frame taken every 3.18 s (approximately 67 s elapsed time per second of the movie). Scale bar, 10 µm. GFP-MNPs delivered by microinjection. Selected frames from this movie are shown in fig. S2C. See also additional relevant extra movies S11-S13 accessible via https://doi.org/10.6084/m9.figshare.22155083.

### Supplementary movie S2_Fig. 1E

Timelapse video (3i Marianas spinning disk) showing remote manipulation of endogenous Rab25 endosomes demonstrating a direct role in cell protrusion. A2780 DExCon-modified NB^GFP^-mCherry-Rab25 (biop-Amber LUT) cells dox pre-treated (>94 h; 250 ng/ml) stably expressing LifeAct-iRFP670 (F-actin, gem or biob-Azure LUT) on FN-coated coverslips. GFP-MNPs delivered by microinjection (biop-SpringGreen LUT). Individual or merged channels as indicated in the movie. Magnetic tweezers/tip visible as shadow in brightfield (left). GFP-MNPs/NB^GFP^-mCherry relocalization followed by local F-actin-dependent protrusion changes, yellow arrows (with cytochalasin D treatment, red arrows). The timelapse video comprises a total duration of 76 minutes, with a frame shown every 30 s (00:00 – 67:10; approximately 450 s elapsed time per second of the movie) or 12 s (71:08 – 76:32; approximately 180 s elapsed time per second of the movie; Cytochalasin treatment from time 68:00), as indicated by the depicted time interval. Note: fast 2s or 5s time interval was captured for the adjustment and re-adjustment of the magnet tip position, but for the movie clarity resliced to 30s (or 12s, respectively) interval. Scale bar, 20 µm. Selected key frames from this movie are shown in Fig. 1E. See also additional relevant movies S14-S15 accessible via https://doi.org/10.6084/m9.figshare.22155083.

### Supplementary movie S3_Fig. 2E

Timelapse video (Andor Dragonfly spinning disk) showing magnetic manipulation of membrane-free Rab25 in A2780 ovarian cancer cells to control protrusion outgrowth. A2780 stably co-expressing LifeAct-iRFP670 (F-actin, gem LUT) with NB^GFP^-mCherry-Rab25 dC mutant (biop-Amber LUT) on FN-coated coverslips. GFP-MNPs delivered by microinjection (biop-SpringGreen LUT). Individual or merged (GFP-MNPs/F-actin/brightfield) channels as indicated in the movie. Magnetic tweezers/tip visible as shadow in brightfield (right). Gradient of NB^GFP^-mCherry-Rab25 dC and GFP-MNPs induced by the magnet, blue arrows; protrusion growth reflected with F-actin. Timelapse covers total 45 min with frame taken every 5 s (approximately 135 s elapsed time per second of the movie). Scale bar, 20 µm. Selected key frames from this movie (time stamper here starts from t=5, just before approaching with the magnet) are shown in Fig. 2E.

### Supplementary movie S4_Fig. 3A-F

Timelapse video (3i Marianas spinning disk) showing magnetic manipulation of endogenous Rab25 recycling endosomes in living cells migrating in 3D-Cell-Derived Matrix (CDM) to control F-actin protrusion. A2780 DExCon-modified NB^GFP^-mCherry-Rab25 cells (Red or Red Hot LUT) dox pre-treated (72 h; 250 ng/ml) expressing Lifeact-iRFP670 (F-actin, Grey or gem LUT). GFP-MNPs (Green LUT) delivered by microinjection and repeatedly relocalized using home-made magnetic tip, shadow in merge brightfield/F-actin (middle). Enrichment of endosome-bound NB^GFP^-mCherry-Rab25 and protrusion growth reflected with F-actin, yellow arrow. 100 μM CK666 and 5 μM SMIFH2 treatment as depicted in the movie. The timelapse video comprises a total duration of 128 minutes, with a frame shown every 30 s, as indicated by the depicted time interval (approximately 450 s elapsed time per second of the movie). Note: 4s interval was captured for the adjustment and re-adjustment of the magnet tip position as depicted in Fig. 3H, but for the movie clarity resliced to 30s interval. Scale bar, 20 µm. Selected key frames from this movie are shown in Fig. 3A-F.

### Supplementary movie S5_ fig. S8G

Timelapse video, captured using the Andor Dragonfly spinning disk, illustrates the magnetic manipulation of Rab25 recycling endosomes in living cells migrating in 3D-Cell-Derived Matrix (CDM) to control F-actin protrusion which is abrogated by integrin β1 blockade. A2780 NB^GFP^-mCherry-Rab25 wt (biop-Amber LUT) cell stably expressing LifeAct-iRFP670 (F-actin, biob-Azure LUT) on FN-coated coverslips. GFP-MNPs delivered by microinjection (biop-SpringGreen LUT, middle). Individual or merged channels as indicated in the movie. Magnetic tweezers/tip visible as shadow in merge brightfield/F-actin (right). Original main protrusion without magnetic tip, yellow arrowheads. GFP-MNPs/NB^GFP^-mCherry-Rab25 relocalization followed by local F-actin-dependent protrusion changes, orange arrowheads. Integrin β1 blocking antibody treatment indicated (AIIFB2, 10 μg/ml; 27:00) followed by protrusion retraction/cell rounding, red arrowheads. The timelapse video comprises a total duration of 111 minutes, with a frame shown every 10 s, as indicated by the depicted time interval (approximately 150 s elapsed time per second of the movie). Scale bar 10 μm. Selected key frames from this movie are shown in fig. S8G.

### Supplementary movie S6_Fig. 5B bottom

Timelapse video (3i Marianas spinning disk) showing magnetic re-localization of endogenous membrane-bound Rab25 in FMNL1 depleted cells migrating on FN-coated coverslip. A2780 DExCon-modified NB^GFP^-mCherry-Rab25 FMNL1+/-cells (biop-Amber LUT; dox treated > 94h; 250 ng/ml) stably expressing Lifeact-iRFP670 (F-actin, gem or biob-Azure LUT), microinjected with GFP-MNPs (biop-SpringGreen LUT) and nucleofected with chemically modified siRNA_pool_ anti-FMNL1 (FMNL1 depleted cells). Magnetic tweezers/tip visible as shadow in brightfield/F-actin merge (left). Zoomed inset, merge F-actin/GFP-MNPs/NB^GFP^-mCherry-Rab25; magnetic enrichment, red arrow (no F-actin protrusion outgrowth). Timelapse covers total 37 min with frame taken every 2s (00:00-01:10; approximately 42 s elapsed time per second of the movie) before magnetic approach, 5s (03:00-05:15; approximately 105 s elapsed time per second of the movie) for initial adjustment of the magnet tip position or 30 s (approximately 630 s elapsed time per second of the movie) for long term imaging. Scale bar 10 µm. Movie relevant to Fig. 5B bottom.

### Supplementary movie S7_Fig. 5D top

Timelapse video (3i Marianas spinning disk) showing magnetic manipulation of endogenous Rab25 recycling endosomes in living cells migrating in 3D-Cell-Derived Matrix (CDM) to control F-actin filopodia based protrusions. A2780 DExCon-modified NB^GFP^-mCherry-Rab25 FMNL1+/+ cells (biop-Amber LUT; dox treated > 94h; 250 ng/ml) stably expressing Lifeact-iRFP670 (F-actin, gem or grey or biob-Azure LUT), microinjected with GFP-MNPs (biop-SpringGreen LUT) and nucleofected with non-targeting ctrl siRNA. Individual or merged channels as indicated in the movie. The magnetic tweezers/tip is discernible as a left shadow in brightfield/F-actin merge (top middle). The relocalisation of GFP-MNPs and endosomal NB^GFP^-mCherry-Rab25 towards a magnetic tip correlates with the polymerisation of F-actin and the formation of filopodia-based protrusions, as indicated by the yellow and red arrowheads. Timelapse covers total 75 min 25 s with frame taken every 5 s, as indicated by the depicted time interval (approximately 155 s elapsed time per second of the movie). Scale bar 10 μm. Selected key frames from this movie are shown in Fig. 5D top.

### Supplementary movie S8_Fig. 5D bottom

Timelapse video (3i Marianas spinning disk) showing magnetic re-localization of endogenous membrane-bound Rab25 in FMNL1 depleted cells migrating in 3D CDM. A2780 DExCon-modified NB^GFP^-mCherry-Rab25 FMNL1+/-cells (biop-Amber LUT; dox treated > 94h; 250 ng/ml) stably expressing Lifeact-iRFP670 (F-actin, gem or biob-Azure LUT), microinjected with GFP-MNPs (biop-SpringGreen LUT) and nucleofected with chemically modified siRNA_pool_ anti-FMNL1 (FMNL1 depleted cells). Magnetic tweezers/tip visible as shadow in brightfield/GFP-MNPs or brightfield/F-actin merge (left). Merge F-actin/GFP-MNPs/NB^GFP^-mCherry-Rab25; magnetic enrichment, red arrow (no F-actin protrusion outgrowth). Timelapse covers total 79 min with frame taken every 10 s (00:00-07:10; approximately 300 s elapsed time per second of the movie) for initial adjustment of the magnet tip position or 30 s (07:10-78:40; approximately 900 s elapsed time per second of the movie) for long term imaging. Scale bar, 10 µm. Movie relevant to Fig. 5D bottom.

### Supplementary movie S9_ Fig. 6C

Timelapse video (Andor Dragonfly spinning disk) showing local modulation of RhoA activity by magnetic re-localization of endogenous membrane-bound Rab25 in A2780 ovarian cancer cells migrating on FN-coated coverslip. A2780 DExCon-modified NB^GFP^-mCherry-Rab25 (biop-Amber LUT) cells dox pre-treated (> 72 h; 250 ng/ml) stably co-expressing active RhoA probe iRFP670_3x_-RBD_4x_ (RBD_4x_; gem LUT). GFP-MNPs delivered by microinjection (biop-SpringGreen LUT). Magnetic tweezers/tip visible as shadow in brightfield/RBD_4x_ (top middle). Individual or merged channels as indicated in the movie. Sustained attraction of Rab25 endosomes via bound GFP-MNPs and increased localisation of the active Rho probe in a punctate pattern (top right), yellow arrowheads. Timelapse covers total 25 min with frame taken every 5 s (approximately 150 s elapsed time per second of the movie, respectively). Scale bar, 20 µm. Movie relevant to Fig. 6C.

### Supplementary movie S10_Fig. 6E

Timelapse video (Andor Dragonfly spinning disk) showing local modulation of RhoA activity by magnetic re-localization of membrane-free Rab25 in A2780 ovarian cancer cells migrating on FN-coated coverslip. A2780 stably co-expressing NB^GFP^-mCherry-Rab25 dC mutant (biop-Amber LUT) with active RhoA probe iRFP670_3x_-RBD_4x_ (RBD_4x_; gem LUT). GFP-MNPs delivered by microinjection (biop-SpringGreen LUT). Magnetic tweezers/tip visible as shadow in brightfield/RBD_4x_ (top right). Magnetic control of GFP-MNPs and NB^GFP^-mCherry-Rab25 dC gradient is followed by local increase in the intensity of the active Rho probe, yellow arrowheads. Timelapse covers total 61 min with frame taken every 5 s (approximately 500 s elapsed time per second of the movie, respectively). Scale bar, 10 µm. Movie relevant to Fig. 6E.

## References

Alanko J, and Ivaska J (2016) Endosomes: Emerging Platforms for Integrin-Mediated FAK Signalling Trends Cell Biol 26, 391–398.

Alzahofi N, Welz T, Robinson CL, Page EL, Briggs DA, Stainthorp AK, … Hume AN (2020) Rab27a co-ordinates actin-dependent transport by controlling organelle-associated motors and Gemperle et al. track assembly proteins Nat Commun 11.

Bagci H, Sriskandarajah N, Robert A, Boulais J, Elkholi IE, Tran V, … Côté JF (2020) Mapping the proximity interaction network of the Rhofamily GTPases reveals signalling pathways and regulatory mechanisms Nat Cell Biol 22, 120–134.

Belevich I, Joensuu M, Kumar D, Vihinen H, and Jokitalo E (2016) Microscopy Image Browser: A Platform for Segmentation and Analysis of Multidimensional Datasets PLOS Biol 14, e1002340.

Bennett H, Aguilar-Martinez E, and Adamson AD (2021) CRISPR-mediated knock-in in the mouse embryo using long single stranded DNA donors synthesised by biotinylated PCR Methods 191, 3–14.

Bongaerts M, Aizel K, Secret E, Jan A, Nahar T, Raudzus F, … Coppey M (2020) Parallelized Manipulation of Adherent Living Cells by Magnetic Nanoparticles-Mediated Forces Int J Mol Sci 21, 1–20.

Caswell PT, Chan M, Lindsay AJ, McCaffrey MW, Boettiger D, and Norman JC (2008) Rab-coupling protein coordinates recycling of alpha5beta1 integrin and EGFR1 to promote cell migration in 3D microenvironments J Cell Biol 183, 143–155.

Caswell PT, Spence HJ, Parsons M, White DP, Clark K, Cheng KW, … Norman JC (2007) Rab25 Associates with α5β1 Integrin to Promote Invasive Migration in 3D Microenvironments Dev Cell 13, 496–510.

Caswell PT, and Zech T (2018) Actin-Based Cell Protrusion in a 3D Matrix 28, 823–834.

Cheng KW, Lahad JP, Kuo W, Lapuk A, Yamada K, Auersperg N, … Mills GB (2004) The RAB25 small GTPase determines aggressiveness of ovarian and breast cancers Nat Med 10, 1251–1256.

Cho KH, and Lee HY (2019) Rab25 and RCP in cancer progression Arch Pharm Res.

Cross RA (2006) Myosin’s mechanical ratchet Proc Natl Acad Sci U S A 103, 8911.

Dehapiot B, Clément R, Alégot H, Gazsó-Gerhát G, Philippe JM, and Lecuit T (2020) Assembly of a persistent apical actin network by the formin Frl/Fmnl tunes epithelial cell deformability Nat Cell Biol 2020 227 22, 791–802.

Dhillon K, Aizel K, Broomhall TJ, Secret E, Goodman T, Rotherham M, … Gates MA (2022) Directional control of neurite outgrowth: emerging technologies for Parkinson’s disease using magnetic nanoparticles and magnetic field gradients J R Soc Interface 19.

Dozynkiewicz MAA, Jamieson NBB, Macpherson I, Grindlay J, van den Berghe PVE, von Thun A, … Norman JCC (2012) Rab25 and CLIC3 collaborate to promote integrin recycling from late endosomes/lysosomes and drive cancer progression. Dev Cell 22, 131–45.

Eisler SA, Curado F, Link G, Schulz S, Noack M, Steinke M, … Hausser A (2018) A rho signaling network links microtubules to PKD controlled carrier transport to focal adhesions Elife 7.

Etoc F, Vicario C, Lisse D, Siaugue J-MM, Piehler J, Coppey M, and Dahan M (2015) Magnetogenetic Control of Protein Gradients Inside Living Cells with High Spatial and Temporal Resolution Nano Lett 15, 3487–3494.

Eva R, Crisp S, Marland JRK, Norman JC, Kanamarlapudi V, Ffrench-Constant C, and Fawcett JW (2012) ARF6 directs axon transport and traffic of integrins and regulates axon growth in adult DRG neurons J Neurosci 32, 10352–10364.

Gaston C, De Beco S, Doss B, Pan M, Gauquelin E, D’Alessandro J, … Delacour D (2021) EpCAM promotes endosomal modulation of the cortical RhoA zone for epithelial organization Nat Commun 12.

Gebhardt C, Breitenbach U, Richter KH, Fürstenberger G, Mauch C, Angel P, and Hess J (2005) c-Fos-Dependent Induction of the Small Ras-Related GTPase Rab11a in Skin Carcinogenesis Am J Pathol 167, 243–253.

Gemperle J, Harrison TS, Flett C, Adamson AD, and Caswell PT (2022) On demand expression control of endogenous genes with DExCon, DExogron and LUXon reveals differential dynamics of Rab11 family members Elife 11, 1–39.

Golachowska MR, Hoekstra D, and van IJ zendoorn SCD (2010) Recycling endosomes in apical plasma membrane domain formation and epithelial cell polarity Trends Cell Biol.

Goldenring JR, Shen KR, Vaughan HD, and Modlin IM (1993) Identification of a small GTP-binding protein, Rab25, expressed in the gastrointestinal mucosa, kidney, and lung J Biol Chem.

Gomez TS, Kumar K, Medeiros RB, Shimizu Y, Leibson PJ, and Billadeau DDD (2007) Formins regulate the actin-related protein 2/3 complex-independent polarization of the centrosome to the immunological synapse Immunity 26, 177–190.

Han Y, Eppinger E, Schuster IG, Weigand LU, Liang X, Kremmer E, … Krackhardt AM (2009) Formin-like 1 (FMNL1) is regulated by N-terminal myristoylation and induces polarized membrane blebbing J Biol Chem 284, 33409–33417.

Hetmanski JHR, de Belly H, Busnelli I, Waring T, Nair R V., Sokleva V, … Caswell PT (2019) Membrane Tension Orchestrates Rear Retraction in Matrix-Directed Cell Migration Dev Cell 51, 460.

Hetmanski JHR, Jones MC, Chunara F, Schwartz JM, and Caswell PT (2021) Combinatorial mathematical modelling approaches to interrogate rear retraction dynamics in 3D cell migration PLoS Comput Biol 17.

Higuchi Y, Ashwin P, Roger Y, and Steinberg G (2014) Early endosome motility spatially organizes polysome distribution J Cell Biol 204, 343–357.

Hoogenraad CC, Akhmanova A, Howell SA, Dortland BR, De Zeeuw CI, Willemsen R, … Galjart N (2001) Mammalian Golgi-associated Bicaudal-D2 functions in the dynein-dynactin pathway by interacting with these complexes EMBO J 20, 4041–4054.

Jacquemet G, Green DM, Bridgewater RE, von Kriegsheim A, Humphries MJ, Norman JC, and Caswell PT (2013) RCP-driven α5β1 recycling suppresses Rac and promotes RhoA activity via the RacGAP1–IQGAP1 complex J Cell Biol 202, 917–935.

Jacquemet G, Humphries MJ, and Caswell PT (2013) Role of adhesion receptor trafficking in 3D cell migration Curr Opin Cell Biol 25, 627–632.

Jakobs MA, Dimitracopoulos A, and Franze K (2019) Kymobutler, a deep learning software for automated kymograph analysis Elife 8.

Jeong BY, Cho KH, Jeong KJ, Park YY, Kim JM, Rha SY, … Lee HY (2018) Rab25 augments cancer cell invasiveness through a β1 integrin/EGFR/VEGF-A/Snail signaling axis and expression of fascin Exp Mol Med 2018 501 50, e435–e435.

Jin H, Tang Y, Yang L, Peng X, Li B, Fan Q, … Li H (2021) Rab GTPases: Central Coordinators of Membrane Trafficking in Cancer Front Cell Dev Biol 9, 1398.

Kappen M, Gemperle J, Secret E, Flesch J, Caswell PT, Coppey M, … Piehler J (2024) Biofunctional coating of synthetic magnetic nanoparticles enables magnetogenetic control of protein functions inside cells BioRxiv 2024.10.31.621314.

Kaukonen R, Jacquemet G, Hamidi H, and Ivaska J (2017) Cell-derived matrices for studying cell proliferation and directional migration in a complex 3D microenvironment Nat Protoc 12, 2376–2390.

Kawano F, Suzuki H, Furuya A, and Sato M (2015) Engineered pairs of distinct photoswitches for optogenetic control of cellular proteins Nat Commun 6, 6256.

Keizer VIP, Grosse-Holz S, Woringer M, Zambon L, Aizel K, Bongaerts M, … Coulon A (2022) Live-cell micromanipulation of a genomic locus reveals interphase chromatin mechanics Science (80-) 377, 489–495.

Kelly EE, Horgan CP, and McCaffrey MW (2012) Rab11 proteins in health and disease Biochem Soc Trans 40, 1360–1367.

Kowarz E, Löscher D, and Marschalek R (2015) Optimized Sleeping Beauty transposons rapidly generate stable transgenic cell lines Biotechnol J 10, 647–653.

Li H, Beckman KA, Pessino V, Huang B, Weissman JS, and Leonetti MD (2017) Design and specificity of long ssDNA donors for CRISPR-based knock-in BioRxiv 178905.

Liße D, Monzel C, Vicario C, Manzi J, Maurin I, Coppey M, … Dahan M (2017) Engineered Ferritin for Magnetogenetic Manipulation of Proteins and Organelles Inside Living Cells Adv Mater 29, 1–7.

Mahlandt EK, Arts JJG, van der Meer WJ, van der Linden FH, Tol S, van Buul JD, … Goedhart J (2021) Visualizing endogenous Rho activity with an improved localization-based, genetically encoded biosensor J Cell Sci 134.

Matsuzaki F, Shirane M, Matsumoto M, and Nakayama KI (2011) Protrudin serves as an adaptor molecule that connects KIF5 and its Gemperle et al. cargoes in vesicular transport during process formation 22.

Miller EW, and Blystone SD (2019) The carboxy-terminus of the formin FMNL1? bundles actin to potentiate adenocarcinoma migration J Cell Biochem 120, 14383–14404.

Miller MR, Miller EW, and Blystone SD (2017) Noncanonical activity of the podosomal formin FMNL1γ supports immune cell migration J Cell Sci 130, 1730.

Mitra S, Cheng KWWW, and Mills GBBB (2012) Rab25 in Cancer: A brief update Biochem Soc Trans 40.

Muller PAJ, Caswell PT, Doyle B, Iwanicki MP, Tan EH, Karim S, … Vousden KH (2009) Mutant p53 drives invasion by promoting integrin recycling Cell 139, 1327–1341.

Nijenhuis W, van Grinsven MMP, and Kapitein LC (2020) An optimized toolbox for the optogenetic control of intracellular transport J Cell Biol 219.

Nørholm MHH (2010) A mutant Pfu DNA polymerase designed for advanced uracil-excision DNA engineering BMC Biotechnol 10.

Nürnberg A, Kitzing T, and Grosse R (2011) Nucleating actin for invasion Nat Rev Cancer 11, 117–187.

Ossipova O, Kim K, Lake BB, Itoh K, Ioannou A, and Sokol SY (2014) Role of Rab11 in planar cell polarity and apical constriction during vertebrate neural tube closure Nat Commun 2014 51 5, 1–8.

Padilla-Rodriguez M, Parker S, Adams D, Westerling T, Puleo J, Watson A, … Mouneimne G (2018) The actin cytoskeletal architecture of estrogen receptor positive breast cancer cells suppresses invasion Nat Commun 9.

Paul NR, Allen JL, Chapman A, Morlan-Mairal M, Zindy E, Jacquemet G, … Caswell PT (2015) α5β1 integrin recycling promotes Arp2/3-independent cancer cell invasion via the formin FHOD3. J Cell Biol 210, 1013–31.

Phuyal S, Romani P, Dupont S, and Farhan H (2023) Mechanobiology of organelles: illuminating their roles in mechanosensing and mechanotransduction Trends Cell Biol.

Piccinini F, Kiss A, and Horvath P (2015) CellTracker (not only) for dummies. Bioinformatics btv686-.

Pylypenko O, Welz T, Tittel J, Kollmar M, Chardon F, Malherbe G, … Kerkhoff E (2016) Coordinated recruitment of spir actin nucleators and myosin V motors to rab11 vesicle membranes Elife 5.

Ray GS, Lee JR, Nwokeji K, Mills LR, and Goldenring JR (1997) Increased immunoreactivity for Rab11, a small GTP-binding protein, in low-grade dysplastic Barrett’s epithelia. Lab Invest 77, 503–511.

Sadowski L, Pilecka I, and Miaczynska M (2009) Signaling from endosomes: location makes a difference Exp Cell Res 315, 1601–1609.

Schlierf B, Fey GH, Hauber J, Hocke GM, and Rosorius O (2000) Rab11b is essential for recycling of transferrin to the plasma membrane Exp Cell Res 259, 257–265.

Schnitzer MJ, Visscher K, and Block SM (2000) Force production by single kinesin motors Nat Cell Biol 2, 718–723.

Schuh M (2011) An actin-dependent mechanism for long-range vesicle transport Nat Cell Biol 13, 1431–1436.

Scita G, and Di Fiore PP (2010) The endocytic matrix Nature 463, 464–473.

Shukla S, Troitskaia A, Swarna N, Maity BK, Tjioe M, Bookwalter CS, … Selvin PR (2022) High-Throughput Force Measurement of Individual Kinesin-1 Motors during Multi-Motor Transport Nanoscale 14, 12463.

Sittewelle M, and Royle SJ (2023) Passive diffusion accounts for the majority of intracellular nanovesicle transport Life Sci Alliance 7.

Steketee MB, Moysidis SN, Jin X, Weinstein JE, Pita-thomas W, Raju HB, … Goldberg JL (2011) Nanoparticle-mediated signaling endosome localization regulates growth cone motility and neurite growth PNAS 108.

Tseng Q, Duchemin-Pelletier E, Deshiere A, Balland M, Guilloud H, Filhol O, and Thery M (2012) Spatial organization of the extracellular matrix regulates cell-cell junction positioning Proc Natl Acad Sci U S A 109, 1506–1511.

Vaidžiulyte K, Macé AS, Battistella A, Beng W, Schauer K, and Coppey M (2022) Persistent cell migration emerges from a coupling between protrusion dynamics and polarized trafficking Elife 11.

van Bergeijk P, Adrian M, Hoogenraad CC, and Kapitein LC (2015) Optogenetic control of organelle transport and positioning. Nature 518, 111–114.

Vassilev V, Platek A, Hiver S, Enomoto H, and Takeichi M (2017) Catenins Steer Cell Migration via Stabilization of Front-Rear Polarity Dev Cell 43, 463-479.e5.

Wang S, Hu C, Wu F, and He S (2017) Rab25 GTPase: Functional roles in cancer Oncotarget 8, 64591–64599.

Willoughby PM, Allen M, Yu J, Korytnikov R, Chen T, Liu Y, … Bruce AEE (2021) The recycling endosome protein Rab25 coordinates collective cell movements in the zebrafish surface epithelium Elife 10.

Wilson B, Flett C, Gemperle J, Lawless C, Hartshorn M, Hinde E, … Caswell PT (2023) Proximity labelling identifies pro-migratory endocytic recycling cargo and machinery of the Rab4 and Rab11 families J Cell Sci 136.

Yayoshi-Yamamoto S, Taniuchi I, and Watanabe T (2000) FRL, a novel formin-related protein, binds to Rac and regulates cell motility and survival of macrophages Mol Cell Biol 20, 6872–6881.

Zerial M, and McBride H (2001) Rab proteins as membrane organizers Nat Rev Mol Cell Biol.

