## Additional supplementary movie legends for "Live-cell magnetic micromanipulation of recycling endosomes reveals their direct effect on actin-based protrusions to promote invasive migration"

Included full-resolution live-cell imaging movies (Spinning Disc) are separate files relevant to the study "Live-cell magnetic micromanipulation of recycling endosomes reveals their direct effect on actin-based protrusion to promote invasive migration" and can be accessed via https://doi.org/10.6084/m9.figshare.22155083. Movies included in S1-S10 are of particular significance for the purposes of this study. Movies S11-S22 serve to provide supplementary corroborative evidence which could not be incorporated directly with this study.

### Supplementary movie S11_ fig. S2D

Timelapse video (Andor Dragonfly spinning disk) showing reversible magnetic manipulation, attraction and release kinetics of $NB^{GFP}$-mCherry (biop-Amber LUT) and GFP-MNPs (biop-SpringGreen LUT) in A2780 ovarian cancer cells stably co-expressing LifeAct-iRFP670 (F-actin, gem LUT) on FN-coated coverslips. F-actin shown as merge with brightfield (top), with GFP-MNPs (middle) or with $NB^{GFP}$-mCherry (bottom). Magnetic tweezers/tip visible as shadow in brightfield (reversible positioned in close proximity to cells or moved far away). GFP-MNPs and $NB^{GFP}$-mCherry relocalization to various subcellular locations, yellow arrows. GFP-MNPS delivered by microinjection. Timelapse covers total 20 min with frame taken every 3,79 s (approximately 114 s elapsed time per second of the movie). Bleaching of mCherry signal was automatically compensated for by Stack Contrast Adjustment plugin in Fiji. Scale bar, 10 µm. GFP-MNPs delivered by microinjection. Selected key frames from this movie are shown in fig. S2D. See also additional relevant extra movies S1; S12-13 accessible via https://doi.org/10.6084/m9.figshare.22155083.

### Supplementary movie S12_ extra_magnetic manipulation

Timelapse video (Andor Dragonfly spinning disk) of remote un-limited magnetic manipulation of GFP-MNPs (biop-SpringGreen LUT) and $NB^{GFP}$-mCherry (biop-Amber LUT) inside living A2780 cells stably expressing LifeAct-iRFP670 (F-actin, biob-Azure LUT) on FN-coated coverslips. Merge: GFP-MNPs/ $NB^{GFP}$-mCherry/F-actin, left; MNPs/ $NB^{GFP}$-mCherry/F-actin/Brightfield, right. Magnetic tweezers/tip visible as shadow in brightfield merge re-positioned from right to left across cells. Quantitative GFP-MNPs and $NB^{GFP}$-mCherry relocalization from a variety of subcellular locations, including across whole cells, in multiple cells simultaneously. GFP-MNPS delivered by microinjection. Note: In cells with GFP-MNPs microinjected levels < $NB^{GFP}$-mCherry levels, GFP-MNPs visible as aggregates (crosslinked) that are not amenable to movement with magnetic force (not observed with A2780 DExCon-modified $NB^{GFP}$-mCherry-Rab25 cells). Timelapse covers total 50 min with frame taken every 5 s (approximately 500 s elapsed time per second of the movie). Scale bar, 10 µm. See also additional relevant extra movies S1; S11; S13 accessible via https://doi.org/10.6084/m9.figshare.22155083.

### Supplementary movie S13_ fig. S2E

Timelapse video (3i Marianas spinning disk) showing magnetic re-localization of both GFP-MNPs and $NB^{GFP}$-mCherry inside living A2780 cells without any significant visible effect on F-actin protrusions despite sustained attraction of GFP-MNPs, red arrow. A2780 cells stably co-expressing $NB^{GFP}$-mCherry (biop-Amber LUT; right) and LifeAct-iRFP670 (F-actin, biob-Azure LUT) on FN-coated coverslips. GFP-MNPs (biop-SpringGreen LUT; middle) delivered by microinjection. Merge: F-actin/Brightfield, left. GFP-MNPs/$NB^{GFP}$-mCherry/F-actin, inset (middle). Magnetic tweezers/tip visible as shadow in brightfield merge (bottom right). Note: In cells with GFP-MNPs microinjected levels < $NB^{GFP}$-mCherry levels, some GFP-MNPs visible as aggregates (crosslinked) that are not amenable to movement with magnetic force (not observed with A2780 DExCon-modified $NB^{GFP}$-mCherry-Rab25 cells). Timelapse covers total 37 min with frame taken every 12s (time 0:00 – 1:44; approx. 252 s elapsed time per second of the movie) for magnet-free phase, every 4s (3:20 – 5:00 and 33:20 – 37:20; approx. 84 s elapsed time per second of the movie) for initial and following adjustment of the magnet tip position, every 60s for long term imaging (5:00 – 33:00; approx. 1260 s elapsed time per second of the movie). Scale bar, 20 µm. Selected key frames from this movie are shown in fig. S2E. See also additional relevant extra movies S1; S11; S12 accessible via https://doi.org/10.6084/m9.figshare.22155083.

### Supplementary movie S14_fig. S3A

Timelapse video (3i Marianas spinning disk) of magnetic re-localization of endogenous membrane-bound Rab25 in A2780 ovarian cancer cells with visible effect on F-actin protrusions which was not blocked by Arp2/3 inhibitor CK666 (100 µM; time indicated in the movie). A2780 DExCon-modified $NB^{GFP}$-mCherry-Rab25 (biop-Amber LUT) cells dox pre-treated (>94 h; 250 ng/ml) stably expressing LifeAct-iRFP670 (F-actin, gem LUT) on FN-coated coverslips. GFP-MNPs delivered by microinjection (biop-SpringGreen LUT). Individual or merged channels as indicated in the movie. Magnetic tweezers/tip visible as shadow in brightfield/GFP-MNPs (left; boxed area show zoomed insets) or brightfield/F-actin/GFP-MNPs (middle left) merge. GFP-MNPs (GFP-nanop.) and $NB^{GFP}$-mCherry (Nan-mCh) relocalization followed by local F-actin dependent protrusion changes, yellow arrows. Timelapse video comprises a total duration of 103 minutes, with a frame captured every 2s (time 0:00 – 6:30; approx. 120 s elapsed time per second of the movie) for initial and following adjustment of the magnet tip position (cells focused without magnetic field, then approached with magnetic tip at 01:28 with 1 minute lag not visualized on the time stamp) or 30s interval (6:30 – 74:00; approx. 1800 s elapsed time per second of the movie) for long term imaging followed with 2s interval prior and after CK666 treatment (74:00 – 104:56; magnetic tip re-adjustments and re-focus), see depicted stamped time interval. Scale bar, 20 µm. Selected key frames from this movie are shown in fig. S3A. See also additional relevant movies S2, S15 accessible via https://doi.org/10.6084/m9.figshare.22155083.

### Supplementary movie S15_fig. S3B

Timelapse video (3i Marianas spinning disk) of magnetic re-localization of endogenous membrane-bound Rab25 in A2780 ovarian cancer cells, treatment with formin inhibitor SMIFH2 (5 µM, later 25 µM; time indicated in the movie) blocked F-actin protrusion growth. A2780 DExCon-modified $NB^{GFP}$-mCherry-Rab25 (biop-Amber LUT) cells dox pre-treated (>94 h; 250 ng/ml) stably expressing LifeAct-iRFP670 (F-actin, gem LUT) on FN-coated coverslips. GFP-MNPs delivered by microinjection (gem or biop-SpringGreen LUT). Individual or merged channels as indicated in the movie. Magnetic tweezers/tip visible as shadow in brightfield/F-actin merge (middle). GFP-MNPs (GFP-nanop.), $NB^{GFP}$-mCherry (Nan-mCh) distribution changes and retraction of F-actin based protrusions indicated by red arrowheads. Timelapse video comprises a total duration of 42 minutes, with a frame captured every 30s (00:00 – 39:20; approx. 840 s elapsed time per second of the movie) for long term imaging followed by 4s interval (39:20 – 42:12; approx. 112 s elapsed time per second of the movie; magnetic tip re-adjustments and additional SMIFH2 treatment). Scale bar 20 µm. Selected key frames from this movie are shown in fig. S3B and fig. S4C. See also additional relevant movies S2, S14 accessible via https://doi.org/10.6084/m9.figshare.22155083.

### Supplementary movie S16_Fig. 1F

Tomogram animation with segmented recycling endosomes (light blue) and GFP-MNPs (orange) in A2780 DExCon-modified $NB^{GFP}$-mCherry-Rab25 cells (pre-treated with dox >94 h; 250 ng/ml). GFP-MNPs delivered by electroporation. Tomogram animation movie generated from STEM tomography relevant to Fig. 1F.

### Supplementary movie S17_fig. S5B

Timelapse video (Andor Dragonfly spinning disk) showing remote manipulation of Rab25 endosomes demonstrating a direct role in cell protrusion independent of initial cell polarity. A2780 stably co-expressing $NB^{GFP}$-mCherry-Rab25 (left) and LifeAct-iRFP670 (F-actin, right) on FN-coated coverslips. GFP-MNPs delivered by microinjection (middle). Magnetic tweezers/tip visible as shadow in brightfield/F-actin merge (right). GFP-MNPs and $NB^{GFP}$-mCherry relocalization followed by local F-actin dependent protrusion changes, blue arrowheads. The timelapse video comprises a total duration of 21 minutes, with a frame shown every 30 s (approximately 210 s elapsed time per second of the movie), as indicated by the depicted time interval. Note: Approximately 25% of $NB^{GFP}$-mCherry-Rab25 wt overexpressing cells exhibited a higher proportion of diffusively localized Rab25 (not observed with A2780 DExCon-modified $NB^{GFP}$-mCherry-Rab25 cells) with fast relocalization kinetics (not included in the analysis of Magnetic attraction and release kinetics). Scale bar, 10 µm. Selected key frames from this movie are shown in fig. S5B. See also additional relevant movies S2, S4, S5, S7 and S14 accessible via https://doi.org/10.6084/m9.figshare.22155083.

### Supplementary movie S18_fig. S7A

Timelapse video (3i Marianas spinning disk) showing magnetic manipulation of $NB^{GFP}$-mCherry and GFP-MNPs in living cells migrating in 3D-Cell-Derived Matrix (CDM). A2780 stably co-expressing $NB^{GFP}$-mCherry (biop-Amber LUT; middle right) with Lifeact-iRFP670 (F-actin; gem LUT). GFP-MNPs (gem or biop-SpringGreen LUT; left and middle left) delivered by microinjection. Magnetic tweezers/tip visible as shadow in brightfield/GFP-MNPs merge (left). Timelapse covers total 14 min (12:00 – 26:00) with frame taken every 30s (approximately 300 s elapsed time per second of the movie). Scale bar, 20 μm. Movie relevant to fig. S7A.

### Supplementary movie S19_fig. S7B

Timelapse video (Andor Dragonfly spinning disk) showing that cell adaptation to mechanosensing of magnetic force by MNP-bound Rab25 endosomes is perturbed by Blebbistatin treatment. A2780 DExCon-modified $NB^{GFP}$-mCherry-Rab25 (bottom left; dox treated > 94 h, 250 ng/ml) stably expressing Lifeact-iRFP670 (F-actin; bottom right, gem LUT) migrating in 3D-Cell-Derived Matrix (CDM). GFP-MNPs (upper left) delivered by microinjection. Magnetic tweezers/tip visible as shadow in brightfield/F-actin merge (top right) or as indicated in the movie. Cell shape changes: adaptation to magnetic force (time 6-18 min, yellow arrowhead), swelling (51 – 111 min, yellow arrowheads) induced by Blebbistatin (5 μM) treatment (23 – 111 min), no magnet in between 23:00 –51:30. Blue arrowheads, magnetic attraction. Timelapse covers total 111 min with frame taken every 30 s (00:00 –06:00 and 23:00 – 111:00; approximately 1500 s elapsed time per second of the movie) for interval before approaching cells with magnetic tip and for long term imaging or 5s (06:00 –22:45; approximately 250 s elapsed time per second of the movie) for initial adjustment of the magnet tip position and adaptation phase. Scale bar, 20 μm. For stress maps see relevant fig. S7B (time stamper here starts from 0, 6 min before approaching cells with the magnet).

### Supplementary movie S20_fig. S7C

Timelapse video (Andor Dragonfly spinning disk) showing magnetic manipulation of membrane-free Rab25 in A2780 ovarian cancer cells in 3D-Cell-Derived Matrix (CDM) with no effect on protrusion outgrowth (red arrowhead). A2780 stably co-expressing LifeAct-iRFP670 (F-actin, gem LUT) with $NB^{GFP}$-mCherry-Rab25 dC mutant (biop-Amber LUT). GFP-MNPs delivered by microinjection (biop-SpringGreen LUT). Individual or merged (GFP-MNPs/F-actin/brightfield) channels as indicated in the movie. Magnetic tweezers/tip visible as shadow in brightfield/F-actin (right). Gradient of $NB^{GFP}$-mCherry- Rab25 dC and GFP-MNPs induced by the magnet, blue arrowheads. Timelapse covers total 23 min with frame taken every 5 s (approximately 150 s elapsed time per second of the movie). Scale bar 20 μm. Movie relevant to fig. S7C.

### Supplementary movie S21_extra_actin tracks

Timelapse video (3i Marianas spinning disk) showing magnetic attraction of Rab25 positive endosomal cluster that promotes formation of actin polymerization hotspot. A2780 DExCon-modified $NB^{GFP}$-mCherry-Rab25 (Nan-mCh-R25; biop-Amber LUT; dox treated 72 h, 250 ng/ml) stably expressing Lifeact-iRFP670 (F-actin; gem or biob-Azure LUT) migrating in 3D-Cell-Derived Matrix (CDM). GFP-MNPs (GFP-nanop.; biop-SpringGreen LUT) delivered by microinjection. Individual or merged channels as indicated in the movie. Magnetic tweezers/tip visible as shadow in brightfield/F-actin merge (bottom left) or as indicated in the movie. Red arrowhead indicates shape changes (cell adaptation to the mechanosensing of magnetic forces upon the repositioning of a magnetic tip). Yellow arrowheads indicate the formation of a visible actin polymerization hot-spot that moved and highly colocalized with Rab25 and GFP-MNPs over >1 hour, co-attracted to the magnet, and returned after its removal. Timelapse covers total 75 min with frame taken every 20 s (approximately 420 s elapsed time per second of the movie). Scale bar 20 μm. Movie relevant to fig. S12.

### Supplementary movie S22_Fig. 6D

Timelapse video (Andor Dragonfly spinning disk) showing local modulation of RhoA activity by magnetic re-localization of endogenous membrane-bound Rab25 in A2780 ovarian cancer cells migrating in 3D-CDM. A2780 DExCon-modified $NB^{GFP}$-mCherry-Rab25 (nanob-mCh-Rab25; biop-Amber LUT) cells dox pre-treated (> 72 h; 250 ng/ml) stably co-expressing active RhoA probe $iRFP670_{3x}$-$RBD_{4x}$ ($RBD_{4x}$; gem LUT). GFP-MNPs delivered by microinjection (GFP-nanop.; biop-

SpringGreen LUT). Magnetic tweezers/tip visible as shadow in merge brightfield/GFP-MNPs (top left) and brightfield/ $RBD_{4x}$ (top middle). Individual or merged channels as indicated in the movie. Yellow arrowheads indicate redistribution of Rab25 endosomes via bound GFP-MNPs and co-redistribution of active RhoA probe in a punctate pattern towards magnetic tweezers. Transient cell contraction adaptation upon magnetic tip re-positioning visible as a local increase of active RhoA signal at the cell rear, cyan arrowhead. Timelapse covers total 30 min with frame taken every 5 s (00:00-11:22; approximately 104 s elapsed time per second of the movie) for initial adjustment of the magnet tip position or 30 s (11:22-29:10; approximately 630 s elapsed time per second of the movie) for long term imaging. Scale bar, 20 µm. Movie relevant to Fig. 6D (time stamper here starts from 0, 2 min 30 s before approaching cells with the magnet).
